# Little evidence for inbreeding depression for birth mass, survival and growth in Antarctic fur seal pups

**DOI:** 10.1101/2024.01.12.575355

**Authors:** A.J. Paijmans, A.L. Berthelsen, R. Nagel, F. Cristaller, N. Kröcker, J. Forcada, J.I. Hoffman

## Abstract

Inbreeding depression, the loss of offspring fitness due to consanguineous mating, is generally detrimental for individual performance and population viability. We therefore investigated inbreeding effects in a declining population of Antarctic fur seals at Bird Island, South Georgia. Here, localised warming has reduced the availability of the seal’s staple diet, Antarctic krill, leading to a temporal increase in the strength of viability selection against inbred offspring, which are increasingly failing to recruit into the adult breeding population. However, it remains unclear whether viability selection operates before or after nutritional independence at weaning. We therefore used microsatellite data from 884 pups and their mothers, and SNP array data from 100 mother-offspring pairs, to quantify the effects of individual and maternal inbreeding on three important neonatal fitness traits: birth mass, survival and growth. We did not find any clear or consistent effects of inbreeding on any of these traits. This suggests that viability selection filters inbred individuals out of the population as juveniles during the time window between weaning and recruitment. Our study brings into focus a poorly understood life-history stage and emphasises the importance of understanding the ecology and threats facing juvenile pinnipeds.

## Introduction

Genetic diversity is fundamental for species survival and adaptation, especially in the Anthropocene where environments are changing rapidly ^1^. Understanding the evolutionary mechanisms that shape genetic diversity is therefore essential for predicting species persistence and for informing conservation policies ^2,3^. Arguably, one of the most important of these mechanisms is inbreeding depression, the loss of offspring fitness that occurs when close relatives mate ^4^. Fitness is reduced because inbreeding increases genome-wide homozygosity, exposing recessive deleterious alleles to selection and, to a lesser extent, reducing heterozygote advantage ^4^. Studies of captive populations have documented strong effects of inbreeding on key fitness traits such as neonatal survival, longevity and reproductive success ^4–6^. However, inbreeding has been more challenging to study in wild populations ^7^, leaving numerous open questions about the importance of inbreeding depression and its dependence on life-history and environmental factors ^8,9^.

Historically, pedigrees were considered the gold standard for quantifying inbreeding and its effects on fitness ^10,11^. However, pedigrees are challenging to construct in wild populations as they require the intensive monitoring of entire populations over multiple generations. Hence, many studies have used the heterozygosity of small panels of genetic markers (typically around 10–20 microsatellites) as a proxy for inbreeding ^12,13^. This approach has uncovered heterozygosity fitness correlations (HFCs) for a variety of life-history, physiological and behavioural traits including birth weight and neonatal survival ^14^, resistance to parasites ^15,16^, aggressiveness ^17^ and even attractiveness ^18^.

Due to the relatively low cost and ease of genotyping microsatellites, the HFC approach can be readily scaled up to include hundreds or even thousands of individuals, facilitating large-scale comparisons over time and across different life-history stages ^e.g. 19–21^. However, small panels of microsatellites typically have limited power to capture genome-wide variation in inbreeding ^22–24^. Consequently, there has been a growing focus on genome-wide approaches capable of quantifying inbreeding with greater precision ^16,25–28^. In particular, many studies are now using single nucleotide polymorphism (SNP) arrays or whole genome resequencing to characterise runs of homozygosity (ROHs), long homozygous tracts that occur when an individual inherits the same identical by descent (IBD) haplotype from both of its parents ^27,29,30^. By summing up the ROHs within an individual’s genome and expressing this as a proportion of the total genome length, the genomic inbreeding coefficient *F*_ROH_ can be calculated ^28,31^.

Studies based on ROHs have confirmed that inbreeding occurs in many animal species and can have strong effects on individual fitness ^9,25,32,33^. However, less is known about the effects of maternal inbreeding on offspring fitness ^although^ ^see^ ^25,26,34^. Filling this knowledge gap is important because transgenerational inbreeding effects can potentially exacerbate individual inbreeding effects, especially in species where mothers provision their offspring ^34^. The interplay between individual and maternal inbreeding might therefore have important consequences for neonatal survival and growth, early-life traits that can influence population dynamics, especially under stressful environmental conditions which can exacerbate inbreeding depression ^35^.

A long-term study of Antarctic fur seals at Bird Island, South Georgia, provides an excellent opportunity to investigate the effects of individual and maternal inbreeding on early-life traits in a wild marine mammal. This species is polygynous ^36^ and adults of both sexes show strong site fidelity ^37,38^, behavioural traits that can promote inbreeding. In line with this, HFCs have already been documented for multiple traits in Antarctic fur seals including body size and reproductive success in males ^18,39,40^ and the propensity of female offspring to recruit into the adult breeding population ^21^. Furthermore, the population of Antarctic fur seals at Bird Island has been steadily declining since the mid 1980s, a pattern that has been linked to a long-term trend of decreasing local krill abundance ^21,41–43^. In parallel, the average heterozygosity of the breeding female population has been steadily increasing over time, implying that homozygous female offspring are increasingly being filtered out of the population prior to recruitment by viability selection ^21^.

In order to better understand the decline of the Antarctic fur seal population at South Georgia, we need to learn more about how viability selection operates against homozygous individuals and exactly when this filtering process takes place. We envisage two, non-mutually exclusive possibilities. First, viability selection may operate during the time-window from birth until weaning at around four months of age ^44^, in which case one would expect to observe a negative relationship between inbreeding and pup survival. Furthermore, as Antarctic fur seal pups are dependent on their mothers for nutrition and protection from predators during this period, one might also expect to observe maternal inbreeding effects on pup survival.

Alternatively, viability selection may operate during the time-window after weaning and prior to recruitment at around 4–6 years of age ^42^. Indeed, one might expect selection against inbred animals to be stronger during this period as the sub-adults are nutritionally independent from their mothers and have to fend for themselves. However, this does not necessarily mean that maternal inbreeding effects will be absent, as recruitment success has been linked in pinnipeds to birth mass ^21^ and mass at weaning ^45^, traits which are themselves reflections of the amount of maternal investment ^46–50^.

In this study, we investigated the above possibilities by analysing relationships between inbreeding and three important early life traits in Antarctic fur seals: pup birth mass, survival and growth. For this, we generated two datasets. The first of these comprised 884 pups and 342 mothers from a single breeding colony genotyped at 39 microsatellites. This dataset spanned four consecutive breeding seasons, including one of the worst years on record in terms of female breeding numbers, pup birth mass and food availability (Figure 1). To measure pup growth, the animals were weighed at birth and were subsequently recaptured and weighed again at around 50 days of age. The second dataset comprised 100 pups and their mothers sampled from two adjacent breeding colonies during two consecutive breeding seasons ^51^. These animals were fitted with VHF transmitters, which allowed the pups to be recaptured and weighed every ten days from birth until just before moulting at around 60 days of age, allowing the construction of individual growth curves. These pups and their mothers were genotyped on a recently developed 85k SNP array ^52^ for the quantification of genomic inbreeding.

**Figure 1.**
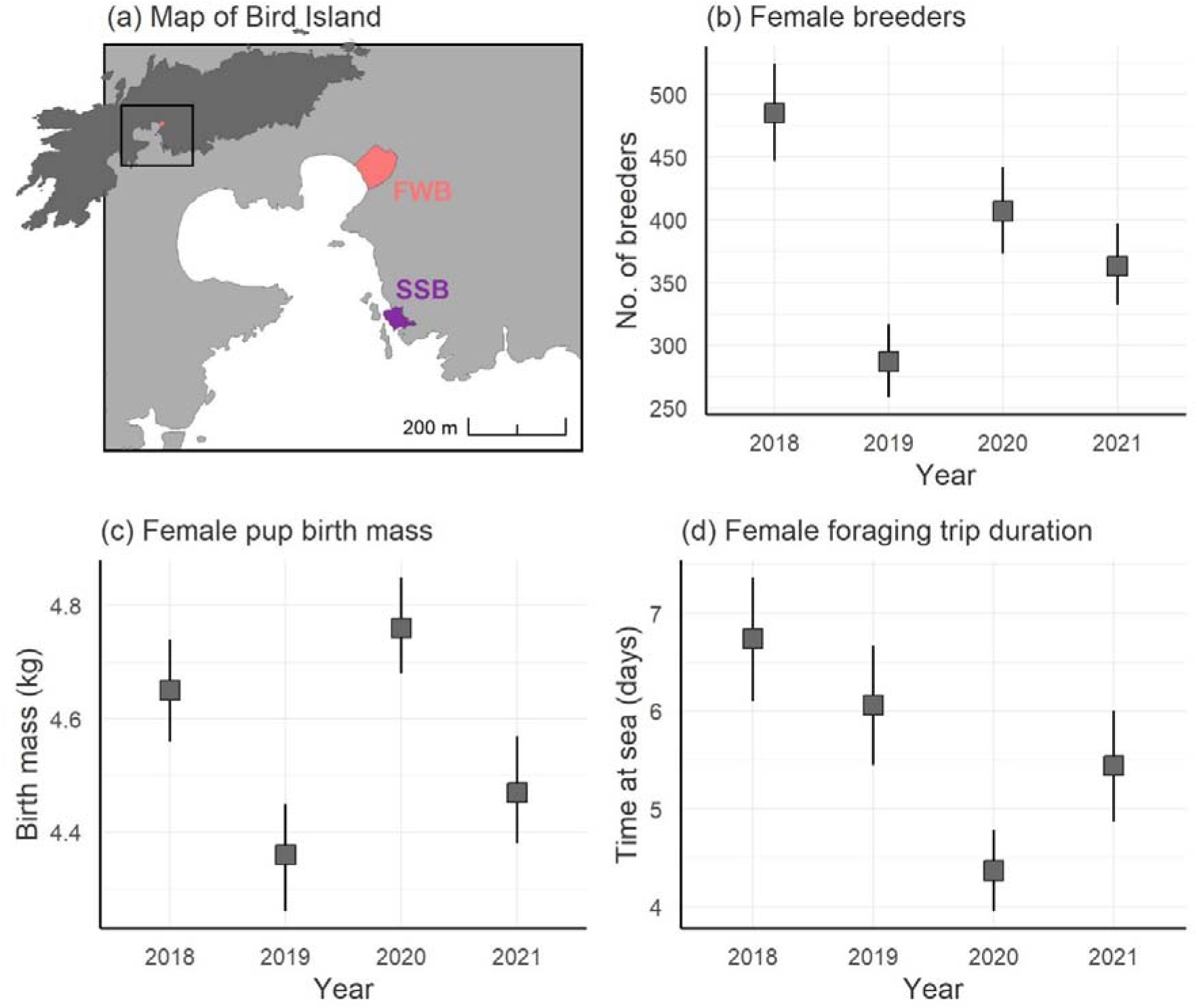
Interannual variation in three measures of season quality. (a) Map of Bird Island, South Georgia, showing the location of two adjacent breeding colonies, the Special Study Beach (SSB) and Freshwater Beach (FWB); (b) Annual numbers of breeding females on SSB; (c) The birth mass of female pups born on SSB; (d) The amount of time spent foraging at sea by mothers (data are from FWB). The squares show the means and the whiskers show the 95% confidence intervals. Data from 2019 and 2020 are already published by Nagel et al.^55^.

We hypothesised that (i) viability selection against inbred Antarctic fur seals occurs mainly after weaning, and hence that an individual pup’s survival will be unrelated to its level of inbreeding. Furthermore, because inbred pups are less likely to recruit ^21^ and recruitment in pinnipeds if often related to body mass ^21,45^, we hypothesised that (ii) inbred pups would exhibit slower growth. We additionally hypothesised that (iii) pups born to inbred mothers will have lower survival and gain less weight; and (iv) inbreeding effects will be more readily detected using ROHs based on tens of thousands of SNPs in comparison to 39 microsatellites.

## Results

To test for effects of individual and maternal inbreeding on pup birth mass, survival and growth, we analysed two datasets. The first of these comprised 884 pups (of which 431 were male and 453 were female) and 342 mothers sampled over four consecutive breeding seasons (2018–2021 inclusive) at the Special Study Beach (SSB; Fig. 1a) of which 721 (81%) survived until the end of the field season. These pups were weighed at birth and subsequently recaptured and weighed again at a mean age of 49 days (range = 12–89 days). Both the pups and their mothers were genotyped at 39 microsatellites that have previously been shown to be in Hardy-Weinberg equilibrium and linkage equilibrium in the study population ^53,54^. The second dataset comprised 100 mother-offspring pairs (i.e. a total of 200 individuals, comprising 51 male and 47 female pups and their mothers) sampled over two consecutive breeding seasons (2019 and 2020) from SSB and Freshwater Beach (FWB; Fig. 1a), of which 76 pups survived until the end of the field season. Repeated weight measurements were gathered from these pups from birth until around 60 days of age. To increase genetic resolution for this subset of animals, we genotyped them on a custom SNP array, resulting in a dataset of 196 individuals (100 pups and 96 mothers) genotyped at 75,101 SNPs.

### Seasonal variation

The four years of our study varied in three measures of breeding season quality (Fig. 1). The 2019 season was among the worst on record ^42^, as indicated by substantially lower numbers of breeding females (Fig. 1b), reduced pup birth mass (Fig. 1c) and relatively high foraging trip durations (Fig. 1d), which indicate that fewer food resources were available to the breeding females. By comparison, the 2020 season had the highest pup birth mass and the shortest foraging trip durations, although female numbers were higher in 2018.

### Molecular inference of inbreeding

Estimates of the two-locus disequilibrium *g*_2_ were positive and significant for both the microsatellite (*g*_2_ = 0.00053, 95% CI = -0.00010–0.00124, *p* = 0.040) and the SNP (*g*_2_ = 0.00012, 95% CI = 0.000086–0.00015, *p* = 0.001) datasets (see SI Fig. 1) indicating that both sets of markers capture variation in inbreeding in the study population. The genomic inbreeding coefficient *F*_ROH_ varied from 0.0417 to 0.1042 and averaged 0.0723 for the mother-pup pairs genotyped on the SNP array.

### Effects of microsatellite heterozygosity on pup birth mass, survival and growth

No effects of either individual or maternal sMLH were found on pup birth mass when controlling for the confounding effects of pup sex, maternal age and breeding season (Fig. 2a, SI Table 1a). Male pups were born heavier than females (*p* < 0.001, Fig 2a, SI Table 1a), older mothers gave birth to heavier pups (*p* < 0.001, Fig 2a, SI Table 1a) and pups were also born lighter in 2019 compared to 2018 (*p* = 0.008 Fig 2a, SI Table 1a). Similarly, when controlling for confounding effects, no effects of individual or maternal sMLH were found on pup survival (Fig. 2b, SI Table 2a), although heavier pups had higher survival (*p* = 0.003, Fig 2b, SI Table 2a) and male pups had lower survival than female pups (*p* = 0.013, Fig 2b, SI Table 2a). In our pup growth model, we found a weak positive effect of sMLH (*p* = 0.021, Fig 2c, SI Table 3a) on growth, and male pups gained significantly more weight than females (*p* = 0.023, Fig 2c, SI Table 3a). Older pups were also significantly heavier (*p* < 0.001, Fig 2c, SI Table 3a) and pups born in 2019, 2020 and 2021 were heavier than pups born in 2018 (*p* < 0.001, Fig 2c, SI Table 3a).

**Figure 2.**
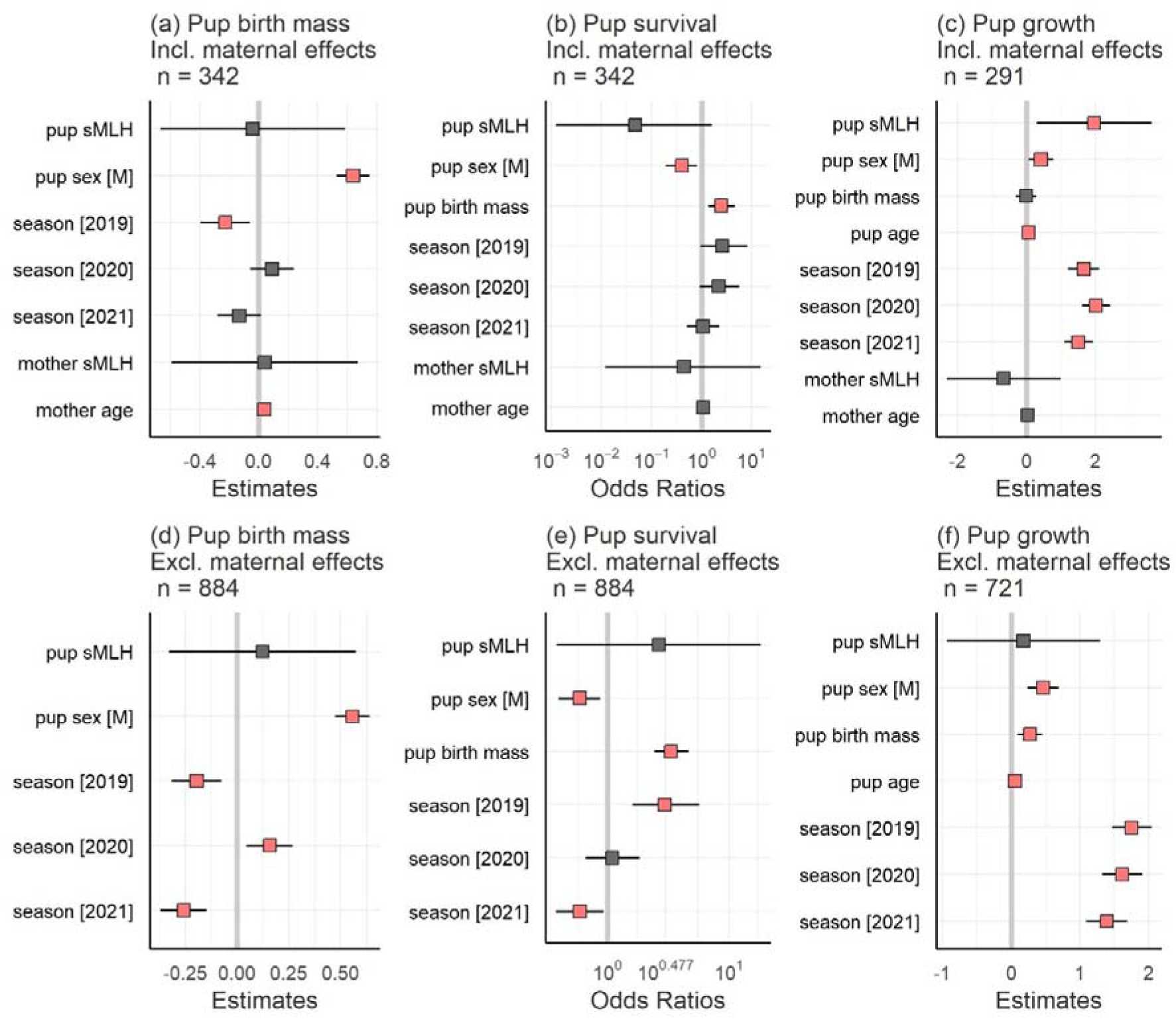
Model estimates and associated 95% confidence intervals for the fixed effects of (a) pup birth mass; (b) pup survival; and (c) pup growth in models including maternal effects; and (d) pup birth mass; (e) pup survival; and (f) pup growth in models excluding maternal effects. Statistically significant relationships are highlighted in salmon pink.

To investigate further, we repeated the above analyses using a larger dataset including many additional pups with unknown mothers (see Methods for details). Overall, the results were similar, with a small number of exceptions. For pup birth mass, all of the seasons showed significant differences compared to 2018 (*p* < 0.01, Fig 2d, SI Table 1b). For pup survival, we also found significant seasonal differences, with more pups surviving in 2019 compared to 2018 and fewer pups surviving in 2021 (*p* < 0.05, Fig 2e, SI Table 2b). For pup growth, we found a significant positive effect of pup birth mass (*p* < 0.01, Fig 2f, SI Table 3b) but the effect of pup sMLH was not significant.

### Effects of genomic inbreeding on pup birth mass, survival and growth

When controlling for the confounding effects of pup sex, breeding season and colony, no effects of individual or maternal *F*_ROH_ were found on pup birth mass (Fig 3a, SI Table 4), nor were there any differences between the two breeding colonies or seasons (Fig 3a, SI Table 4), although male pups were born heavier than females (*p* = 0.021, Fig 3a, SI Table 4). We also found no effects of individual or maternal *F*_ROH_ on pup survival (Fig 3b, SI Table 5), although survivorship was higher at SSB (*p* = 0.028, Fig 3b, SI Table 5) and heavier born pups were more likely to survive (*p* = 0.042, Fig 3b, SI Table 5) when other model components were kept constant. Maternal *F*_ROH_ was positively associated with pup growth (*p* = 0.002, Fig 3c, SI Table 6) and male pups gained more weight than females (*p* < 0.001, Fig 3c, SI Table 6). Pup *F*_ROH_, birth mass, age, season and breeding colony had no effect on growth (Fig 3c, SI Table S6).

**Figure 3.**
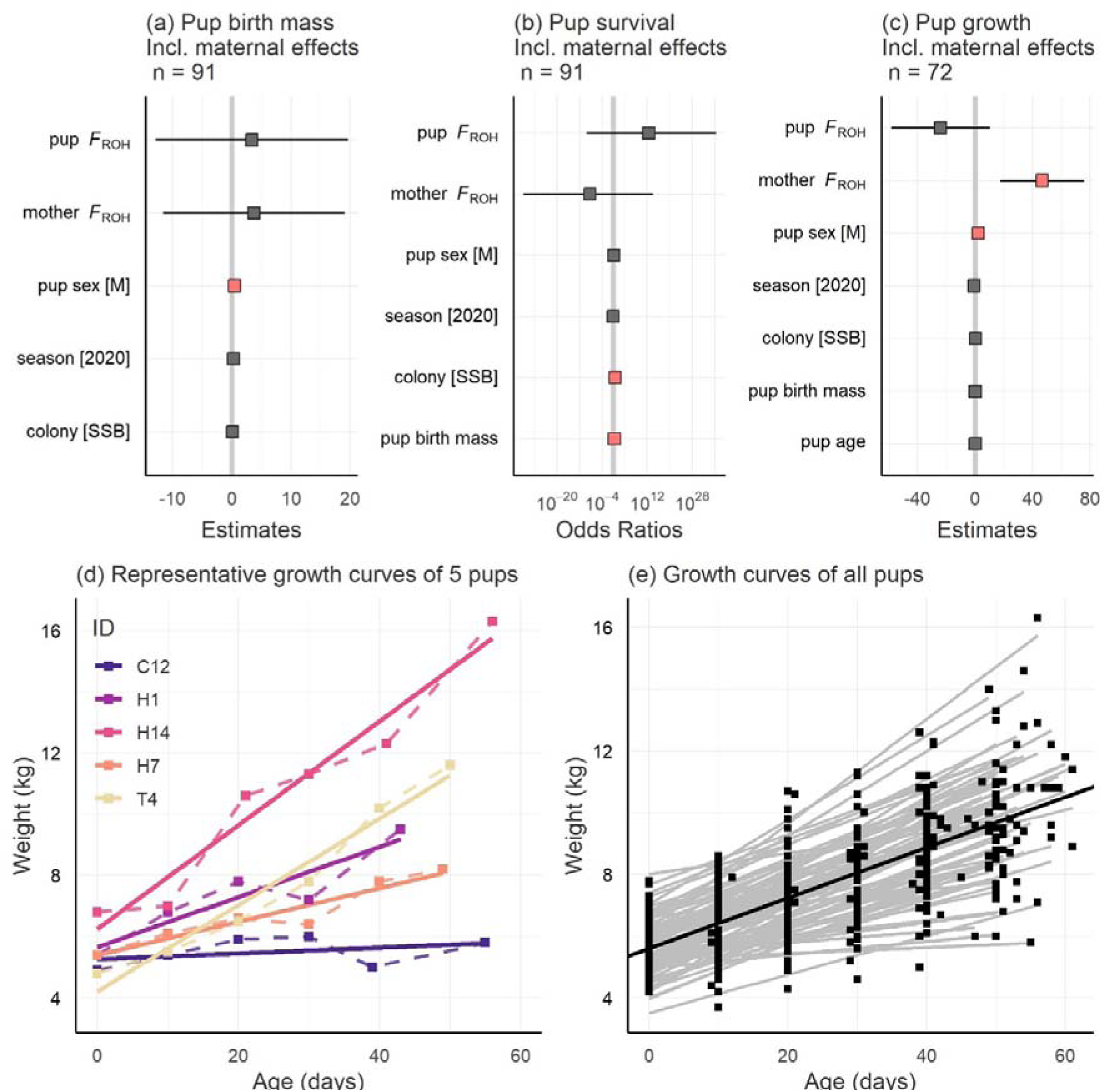
Results of genomic inbreeding analyses, including model estimates and associated 95% confidence intervals for the fixed effects of (a) pup birth mass; (b) pup survival; and (c) pup growth. Statistically significant relationships are highlighted in salmon pink. Panels (d) and (e) depict approximately linear increases in pup mass over time for a representative subset of five pups and for all 100 pups respectively. The dark line in panel (e) indicates the global average rate of change over time (y = 0.082 x + 5.598).

## Discussion

We used molecular and life-history data from an intensively studied Antarctic fur seal population to test for inbreeding depression for three important early acting traits. Using microsatellite and SNP array data, we found no effects of individual or maternal inbreeding on pup birth mass and survival. The results for pup growth were less clear as they depended on the dataset, but again we did not find any consistent evidence for inbreeding depression. Taken together, our results suggest that viability selection against inbred pups is unlikely to be important during the first three months of life. By implication, viability selection against inbred offspring likely operates during the juvenile stage after nutritional independence.

### Study design and comparison to previous studies

Our study complements and builds upon two previous studies of inbreeding depression in Antarctic fur seal pups. The first of these found no effects of individual or maternal heterozygosity at nine microsatellites on pup birth mass and survival ^56^ and the second found no effects of individual heterozygosity at 48 microsatellites on neonatal mortality due to bacterial infection ^53^. However, both of these studies had important limitations. The first had a relatively large sample size of individuals and incorporated maternal effects, but simulations and empirical studies have since shown that nine microsatellites provide a poor estimate of inbreeding in all but the most inbred of populations ^22^. The second study, on the other hand, used a larger panel of microsatellites but focused on a more narrowly defined trait and did not incorporate maternal effects. The current study aimed to produce a more comprehensive and detailed picture of how selection acts in early life by incorporating three main improvements. First, we broadened the focus to include not only pup birth weight and survival, but also growth, inferred from repeated measurements of the same individuals. Understanding the effects of inbreeding on early growth could be important because body size at weaning is known to influence subadult survival and recruitment success in several pinniped species ^45,50,57–59^. Second, genetic parentage analysis allowed us to confirm the maternity of the majority of pups, allowing us to jointly analyse individual and maternal inbreeding effects for all three traits. Third, we genotyped a sufficiently large number of microsatellites to capture a clear signal of identity disequilibrium, as indicated by a statistically significant *g*_2_ statistic, indicating that these markers capture variation in genome-wide inbreeding. We furthermore used a SNP array to quantify genomic inbreeding for a smaller number of animals that were radio tracked from birth until moulting, allowing the construction of detailed individual growth curves.

### Pup birth mass and survival

We found that male pups were born heavier but had lower survival than female pups, older mothers gave birth to heavier pups, and heavier pups had higher survival. Many of these relationships have previously been reported for the study population ^47,60^. However, we did not find any effects of individual or maternal inbreeding on pup birth mass or survival after controlling for the confounding effects of pup sex and breeding season. The clear absence of inbreeding depression for these traits in the current study as well as in two previous studies ^53,56^ implies that viability selection against inbred pups is weak or absent during the first three months of life. This is in contrast to the situation with harbour seals ^14^, grey seals ^61^ and California sea lions ^62^, where microsatellite heterozygosity is strongly associated with neonatal survival. The most likely explanation for this difference is that most Antarctic fur seal pups die of causes that may not have a genetic basis including starvation, trauma ^56^ and predation ^55,63,64^. These findings do not support the hypothesis that viability selection operates to filter inbred pups out of the population prior to weaning. However, it is important to note that we could only follow the pups until they began to moult at around 50–60 days of age, which is around two months before weaning takes place ^44^. Nevertheless, 50% of the pup mortality that we observed in our study occurred in the first seven days of life, and 90% of the pups had died before 35 days, similar to previously reported by ^65^. Consequently, although our data do not extend up to weaning, it seems unlikely that many of our focal pups died between moulting and weaning.

### Pup growth

As body mass at weaning is an important predictor of juvenile survival in pinnipeds ^45,50,57–59^, we hypothesised that the higher recruitment success of outbred individuals might be explained by increased growth rates during early life. We therefore used two approaches to quantify pup growth. First, for a large dataset of pups spanning four consecutive seasons and genotyped at 39 microsatellites, we captured pups shortly after birth and subsequently at tagging 12–89 (mean = 49) days later. We then controlled for the variation in recapture time by including age as a fixed effect in our models. Second, we made use of more detailed growth information available for 100 pups that were recaptured every 10 days from birth until moulting. This revealed growth trajectories to be approximately linear during the first 60 days of life, which justifies our approach of applying linear models of weight gain for both datasets.

Starting with the microsatellite dataset, the model including maternal effects revealed a significant positive association between individual heterozygosity and pup growth, but this was not significant in the model excluding maternal effects. One possible explanation for this discrepancy could be collider bias ^66^, which is a type of selection bias where the likelihood of a sample’s inclusion in a given dataset is influenced by an unknown variable (the “collider”), which is affected by both the response variable (x) and the predictor (y). When the collider is included in the model, it could induce an association between x and y that does not exist, or flip the estimate in the opposite direction if an association does exist. It is possible that this could have occurred in our study, as pups for which maternal information was available were born to tagged females in the study population, who tend to be older and thus more experienced than untagged females. Hence, pups born to untagged females might experience weaker inbreeding depression due to their fitness being primarily determined by the lack of experience of their mothers. However, this seems unlikely given that we did not find a significant effect of mother’s age on pup growth. Alternatively, including maternal sMLH in the model might have masked a significant effect of pup sMLH. However, this is again not supported by our data as the association between pup sMLH and growth remained significant even after removing maternal sMLH from the model (data not shown). Given that no relationship between inbreeding and pup growth was found for the larger microsatellite dataset as well as for the genomic dataset (see below), a final possibility could be type I error.

### Genomic inbreeding

To quantify inbreeding with greater precision, we genotyped 100 pups and their mothers on an 85k SNP array ^52^. These individuals were sampled in two consecutive years from two neighbouring breeding colonies, SSB and FWB ^55^. In line with the results of the microsatellite analyses, no effects of individual or maternal *F*_ROH_ were found on pup birth mass and survival. In addition, a significant effect of colony was found on pup survival, with pups from SSB being more likely to survive. This has been shown previously ^55^ and is due to predation being higher at the lower density FWB colony ^64^.

In contrast to the results of the microsatellite analyses, we found a significant effect of maternal *F*_ROH_ on pup growth, although the direction of the relationship was the opposite to what we originally hypothesised, with inbred mothers tending to have pups that gained more weight. Again, this could potentially be due to type I error given our relatively small sample size for this analysis. However, taking this result at face value, we can think of two alternative explanations. First, because inbreeding tends to reduce longevity ^7,26^, inbred mothers may trade-off current versus future reproduction and invest more heavily into their current offspring. However, theoretical models suggest maternal inbreeding is unlikely to affect optimal parental investment ^67^, while an empirical study of zebra finches found that inbred mothers showed reduced, rather than increased, maternal care ^68^. Another possibility is that Antarctic fur seal pups may be better able to extract resources from inbred mothers, for example via more effective food solicitation. In line with this, it has been shown that maternal provisioning in Antarctic fur seals varies over time and depends on a combination of both maternal and offspring traits, with heavier pups receiving more milk at around one month of age, but maternal mass being the primary determinant of energy allocation in newborn and two-month old pups ^69^. This suggests that who is in control of parental investment is dynamic across the investment period and that maternal traits are more important determinants of maternal care overall.

### Implications

Our findings lend support to the hypothesis that poor environmental conditions select against inbred offspring mainly after nutritional independence in Antarctic fur seals. In line with this, survival has been shown to decline after weaning and is strongly related to sea surface temperature during the first six months of nutritional independence in a closely related pinniped, the subantarctic fur seal ^58^. This environmental dependence of early survival mirrors the situation in Antarctic fur seals, where Forcada et al. ^21^ showed that the strength of viability selection against inbred animals depends on the Southern Annular Mode, a measure of climate variability in the Southern Ocean, which is positively correlated with SST, low krill supply and reduced fur seal viability ^21,42^. Taken together, these studies suggest that juvenile pinnipeds may be particularly vulnerable to the selection pressures imposed by changing environments. Hence, we urgently need to learn more about the ecology of juvenile pinnipeds and the threats facing them during this critical life history stage.

## Conclusion

To summarise our main results, we did not find any significant effects of either individual or maternal inbreeding on pup birth mass and survival. Furthermore, our results for pup growth were not consistent across datasets and methods, leading us to conclude that there is little clear evidence for inbreeding depression for pup growth. While larger sample sizes would be required to reach more definitive conclusions, our results suggest that viability selection against inbred Antarctic fur seals operates mainly after nutritional independence at weaning. Our study therefore brings into focus a life-history stage that is little studied and poorly understood.

## Methods

### Field methods

This study was conducted at an intensively studied breeding population of Antarctic fur seals at Bird Island, South Georgia (54°00024.800 S, 38°03004.100 W) during the austral summers of 2017–2018 to 2020–2021 (hereafter, breeding seasons are referred to by the year in which they ended). Our main study colony (Special Study Beach; SSB, see Fig 1a) was located at a small cobblestone breeding beach (approximately 440 m^2^ at high tide) where a scaffold walkway ^65^ provides safe access to the animals while minimizing disturbance. A second breeding colony, referred to as Freshwater Beach (FWB, see Fig 1a), was located approximately 200 meters to the north.

The seals were captured and restrained following protocols that have been established over more than 40 consecutive years of the long-term monitoring and survey program of the British Antarctic Survey (BAS). As part of this long-term monitoring program, almost a thousand adult females were tagged using cattle ear tags (Dalton Supplies, Henley on Thames, UK) placed in the trailing edge of the foreflipper ^70,71^. The majority of these females were aged from canine tooth sections ^72,73^. Pups were captured on the day of birth, sexed, and weighed. Piglet ear notching pliers were used to collect a small skin sample from the interdigital margin of the foreflipper, which was stored individually in 20% dimethyl sulphoxide (DMSO) saturated with salt ^74^ at −20°C. The pups were then marked with temporary serial numbers by bleaching the fur on their backs before returning them to their mothers, which were tissue sampled later in the season (January–March).

Twice-daily surveys were made of all females and their pups present in the colony from the beginning of November until the end of January. To gather data on growth and survival, we recaptured the pups and weighed them again after ∼49 days (min: 12 days, max: 89 days) and we recorded the identities of any pups that died during this period.

To provide more detailed insights into pup growth, we also collected serial weight measurements from a subset of pups as described by Nagel et al. ^75^. Briefly, during the breeding seasons of 2019 and 2020, a total of 100 unique mother–pup pairs (*n* = 200 individuals) were captured, 50 from SSB and 50 from FWB. Because mothers and their offspring become increasingly mobile as the pups mature ^51^, VHF transmitters were attached to the animals to allow them to be located, recaptured and weighed every ten days from birth until they started to moult (ca. 60 days).

### Genetic analyses

#### Microsatellite genotyping

Total genomic DNA was extracted using an adapted chloroform-isoamylalcohol protocol ^75^ and genotyped for 39 microsatellite loci as described by Paijmans et al. ^54^. Briefly, the microsatellite loci were PCR[amplified in five separate multiplexed reactions using a Type It Kit (Qiagen). Fluorescently labelled PCR products were then resolved by electrophoresis on an ABI 3730xl capillary sequencer (Applied Biosystems, Waltham, MA, USA). Allele sizes were scored automatically using GeneMarker v. 2.6.2 (SoftGenetics, LLC., State College, PA, USA) and the resulting genotypes were manually inspected and corrected where necessary. Those genotypes with fewer than five missing loci were then used to quantify individual standardized multilocus heterozygosity (sMLH) ^15^ using the *sMLH* function of the InbreedR package ^76^. The same package was also used to quantify identity disequilibrium (using the *g*_2_ statistic^77^, with 1,000 permutations).

Up to around ten percent of mother-offspring pairs identified in the field are known to genetically mismatch, probably due to a combination of fostering and milk-stealing ^78^. We therefore used NEWPAT ^79^ to check the maternity of all pups. Any pairs of genotypes with up to three mismatching loci were visually inspected as described by Hoffman et al. ^80^. Mismatches that could be clearly attributed to scoring errors were then corrected. Mothers with zero (*n* = 500, 90%) or one mismatching locus (*n* = 29, 5%) were considered to be biological mothers and were retained in the final dataset, while the remaining mothers (*n* = 26, 5%) were removed.

#### SNP genotyping

Additional genotyping was performed for the 100 pups and their mothers for which detailed growth data were available. These animals were genotyped using a custom 85k Affymetrix SNP array (for details, see Humble et al. ^52^). Quality control was performed in the Axiom Analysis Suite (5.0.1.38, Affymetrix) using the standard parameter threshold settings for diploid organisms. To recover SNPs that were initially classified as “off-target variants”, we used the “Run OTV caller” function in the Axiom Analysis Suite. This resulted in a dataset of 77,895 SNPs (97% of the 85,359 SNPs tiled on the array), of which 75% were categorised as “polymorphic high resolution”, 14% as “no minor homozygote” and 2.5% as “monomorphic high resolution”. SNPs showing a high Mendelian error rate were removed (*n* = 240) using the *--me* flag in PLINK version 1.9 ^81^, with a per-sample error rate of at most 0.05 and a per-variant error rate of at most 0.1. In addition, SNPs that departed significantly from Hardy Weinberg equilibrium (HWE, *n* = 238) were removed using the *--hwe* flag with a *p*-value threshold of 0.001 and the *midp* modifier in PLINK. Finally, we removed SNPs that did not map to the genome and filtered the dataset to only include autosomal SNPs, resulting in a final filtered dataset of 75,101 SNPs.

We then used the SNP data to calculate each individual’s genomic inbreeding coefficient, *F*_ROH_. For this, we first called ROHs on autosomes using the PLINK function *--homozyg* with the parameter settings described by Humble et al. ^52^. Briefly, we called ROH with a minimum length of 1,000 kb and containing at least 20 SNPs while allowing no more than one heterozygous site and a maximum gap of 1,000 kb using the command *-- homozygwindow-snp* 20 *--homozyg-snp* 20 *--homozyg-kb* 1000 --*homozyg-gap* 1000 -- *homozyg-density* 100 *--homozyg-window-missing* 5 *--homozyg-het* 1 *--homozyg-window-het* 1 *--homozyg-window-threshold* 0.05. The sum of the calculated ROH lengths was then divided over the total autosome length (2.28 Gb) to obtain *F*_ROH_. The two-locus heterozygosity disequilibrium (*g_2_*) was also calculated for the SNP data as described above.

### Statistical analyses

We implemented a series of statistical models to investigate whether individual and / or maternal heterozygosity explain a significant proportion of the variation in (i) pup birth mass, (ii) pup survival, and (iii) pup growth. Models of pup growth could only be implemented for surviving pups because the majority of dead pups could either not be recovered or were scavenged by skuas and giant petrels. In order to allow the joint analysis of individual and maternal effects, we initially focused on the subset of pups with known, genetically assigned mothers (342 / 884 pups, 39%). To maximise our sample size, we also ran the same models for the full dataset while excluding maternal effects. These analyses were performed separately for the microsatellite and SNP datasets.

#### Microsatellite data analyses

To test for effects of individual and maternal microsatellite heterozygosity on pup birth mass, we constructed a linear model. Pup and mother sMLH were included as continuous predictor variables. Pup sex (male / female) and season (2017 / 2018 / 2019 / 2020) were included as additional covariates (factors with two and four levels respectively) together with mother’s age (as a continuous variable):

Model 1: Pup birth mass*_i_* = β_Int_ + β_1_ * pup sMLH*_i_* + β_2_ * mother sMLH*_i_* + β_3_ * pup sex*_i_* + β_4_ * season*_i_* + β_5_ * mothers age*_i_* + ε*_i_*

where:

pup birth mass represents the observed value of the i-th individual in the sample, β_Int_, β_1_–β_5_ are regression coefficients for the intercept and the predictor variables, and ε is the random error.

To test for effects of individual and maternal microsatellite heterozygosity on pup survival, we constructed a generalized linear model (GLM) with a binomial error structure. Pup survival was encoded as 1 = survived and 0 = dead. Pup and mother sMLH were included as continuous predictor variables. Pup sex, season and mother’s age were included as additional covariates, together with birth mass (as a continuous variable):

Model 2: 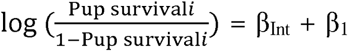 * mother sMLH*_i_*+ β_3_ * pup sex*_i_*+ β_4_ * pup birth mass*_i_* + β_5_ * season*_i_* + β_6_ * mothers age*_i_* + ε*_i_*

where:

Pup survival*_i_* represents the probability that survival is equal to 1 for the *i*-th individual in the sample, β_Int_, β_1_–β_6_ are regression coefficients for the intercept and the predictor variables, and ε is the random error.

To test for effects of individual and maternal microsatellite heterozygosity on pup growth, we constructed a linear model. Growth was calculated as the difference in body mass between birth and recapture. To correct for variation in the number of days between birth and recapture, pup age (defined as the number of days between birth and recapture) was included as a fixed effect in the model. Pup and mother sMLH were included as continuous predictor variables and pup sex, birth mass, season and mother’s age were included as additional covariates:

Model 3: Pup growth*_i_*= β_Int_ + β_1_ * pup sMLH*_i_* + β_2_ * mother sMLH*_i_* + β_3_ * pup sex*_i_* + β_4_ * pup birth mass*_i_*+ β_5_ * pup age*_i_* + β_6_ * season*_i_* + β_7_ * mothers age*_i_* + ε*_i_*

where:

pup survival represents the observed value of the i-th individual in the sample, β_Int_, β_1_–β_7_ are regression coefficients for the intercept and the predictor variables, and ε is the random error.

#### SNP data analysis

To test for effects of individual and maternal inbreeding on birth mass, we constructed a linear model. Pup and mother *F*_ROH_ were included as continuous predictor variables. Pup sex (male / female), colony (SSB / FWB) and season (2019 / 2020) were included as additional two-level covariates:

Model 4: Pup birth mass*_i_* = β_Int_ + β_1_ * pup *F*_ROH_ *_i_* + β_2_ * mother *F*_ROH_ *_i_*+ β_3_ * pup sex*_i_* +β_4_ * colony*_i_* + β_5_ * season*_i_* + ε*_i_*

where:

pup birth mass represents the observed value of the i-th individual in the sample, β_Int_, β_1_–β_5_ are regression coefficients for the intercept and the predictor variables, and ε is the random error.

To test for effects of individual and maternal inbreeding on pup survival, we constructed a generalized linear model (GLM) with a binomial error structure. Pup survival was encoded as 1 = survived and 0 = dead. Pup and mother *F*_ROH_ were included as continuous predictor variables. Pup sex, colony and season were included as two-level covariates as described above and birth mass was included as a continuous covariate.

Model 5: 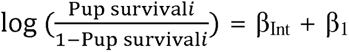 * pup *F*_ROH_ *_i_* + β_2_ * mother *F*_ROH_ *_i_*+ β_3_ * pup sex*_i_*+ β_4_ * pup birth mass*_i_* + β_5_ * colony*_i_* + β_6_ * season*_i_* + ε*_i_*

where:

Pup survival*_i_* represents the probability that survival is equal to 1 for the *i*-th individual in the sample, β_Int_, β_1_–β_6_ are regression coefficients for the intercept and the predictor variables, and ε is the random error.

To test for effects of individual and maternal inbreeding on pup growth, we used the repeated weight measurements of individual pups to model growth curves. In a preliminary analysis, we investigated the fit of linear, logistic and gompertz models to the growth data, and found that pup growth was best described by a linear model (see supplementary R markdown file). We therefore constructed a linear model of pup growth (calculated as the difference in body mass between birth and last capture). The predictor variables in this model were pup *F*_ROH_, mother *F*_ROH_ and the covariates pup sex, birth mass, age, colony and season:

Model 6: Pup growth*_i_*= β_Int_ + β_1_ * pup *F*_ROH_ *_i_* + β_2_ * mother *F*_ROH_ *_i_*+ β_3_ * pup sex*_i_* + β_4_ * pup birth mass*_i_* + β_5_ * pup age*_i_* + β_6_ * season*_i_* + β_7_ * colony*_i_* + ε*_i_*

where:

pup survival represents the observed value of the i-th individual in the sample, β_Int_, β_1_–β_7_ are regression coefficients for the intercept and the predictor variables, and ε is the random error.

For all of the models, the residuals were visually inspected for linearity and equality of error variances (using plots of residuals versus fits) and normality (using Q-Q plots). Testing for under- or over-dispersion was done by comparing the dispersion of simulated and observed residuals. Model inspection was performed using DHARMa ^82^. Analyses and visualisations were implemented in R version 4.0.2 ^83^ using the integrated development environment RStudio ^84^.

## Animal ethics

Fur seal samples were collected as part of the Polar Science for Planet Earth program of the British Antarctic Survey, under permits from the Government of South Georgia and the South Sandwich Islands (GSGSSI, Wildlife and Protected Areas Ordinance (2011), RAP permit numbers 2018/024 and 2019/032). Samples originating from South Georgia Islands were imported into the United Kingdom under permits from the Department for Environment, Food, and Rural Affairs (Animal Health Act, import license number ITIMP18.1397) and from the Convention on International Trade in Endangered Species of Wild Fauna and Flora (import numbers 578938/01-15 and 590196/01-18), and exported under permits issued by the GSGSSI and the UK Department for Environment, Food and Rural Affairs, under European Communities Act 1972. All procedures used were approved by the British Antarctic Survey Animal Welfare and Ethics Review Body (reference no. PEA6, AWERB applications 2018/1050 and 2019/1058).

## Data and code availability

Scripts are provided in the form of an R Markdown file. The scripts and data needed to reproduce all of the analyses and figures can also be accessed via GitHub https://github.com/apaijmans/inbreeding-pup-growth). Microsatellite and SNP data are available via the Zenodo repository, doi:XXX

## Supporting information

R markdown file

## Acknowledgements

This research was supported by the Deutsche Forschungsgemeinschaft (DFG, German Research Foundation) priority programme “Antarctic Research with Comparative Investigations in Arctic Ice Areas” SPP 1158 (project number 424119118) and the SFB TRR 212 (NC³) (Project Numbers 316099922 & 396774617). This work contributes to the Ecosystems project of the British Antarctic Survey, Natural Environmental Research Council, and is part of the Polar Science for Planet Earth Programme.

## Author contributions

Conceived the study: J.I.H. Sample collection and logistics: R.N., J.F. Laboratory work: A.J.P., A.L.B., F.C., N.K. and R.N. Analysed data: A.J.P., A.L.B, J.I.H. Wrote the paper: A.J.P., A.L.B. and J.I.H. All of the authors commented upon and approved the final manuscript.

## Additional information

Correspondence and requests for materials should be addressed to A.J.P. or J.I.H.

