## Supplementary information for "Little evidence for inbreeding depression for birth mass, survival and growth in Antarctic fur seal pups"

Anneke J. Paijmans, Ane Liv Berthelsen, Rebecca Nagel, Felicitas Cristaller, Nicole Kröcker, Jaume Forcada, and Joseph I. Hoffman

**Supplementary Figures**

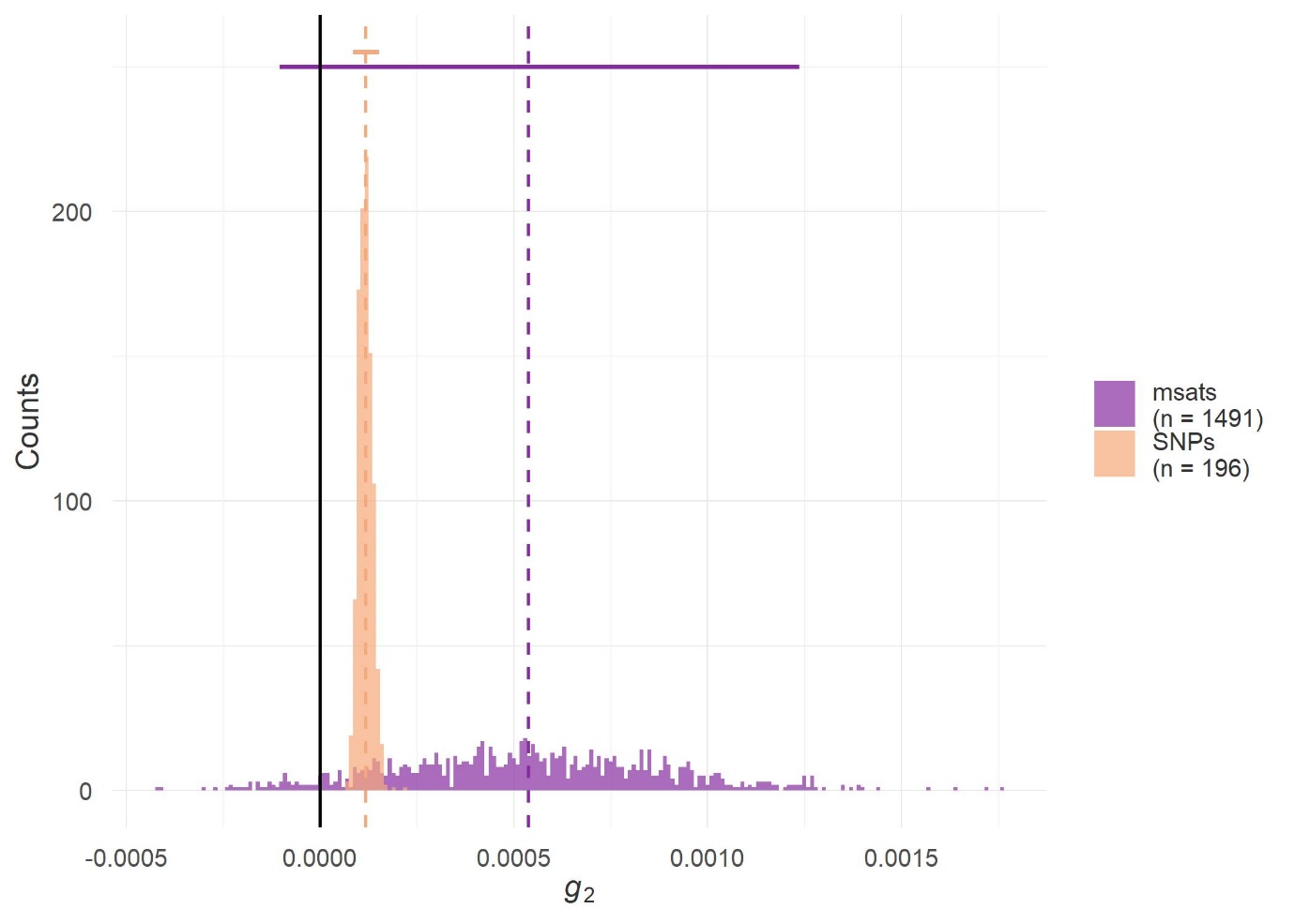

Figure S1. Distribution of identity disequilibrium estimates for both microsatellite data (purple, 1,491 individuals, 39 loci) and SNP array data (orange, 196 individuals, 75,101 autosomal variants). Estimates were based on 1,000 bootstrap replicates and 1,000 permutations. The vertical dashed lines show the empirical *g_2_* estimates and the horizontal bars show the 95% confidence intervals for each dataset. For both datasets, the empirical *g_2_* was significantly larger than zero (microsatellite data: *p* = 0.040, SNP array data: *p* = 0.001).

**Supplementary Tables**

Table S1. Parameter estimates from the linear models testing for the effect of individual and maternal heterozygosity (sMLH) on pup birth mass (a) including maternal genetic diversity and mother age; and (b) excluding maternal effects. Estimates are shown together with confidence intervals (CI), significant *p*-values are in bold. For both models, total number of observations, as well as the variance explained by the predictors (R^2^) and variance adjusted for the number of predictors (R^2^ adjusted) are reported.

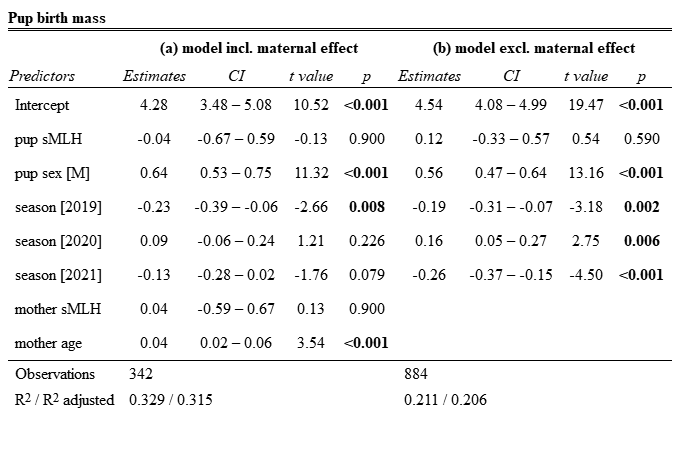

Table S2. Parameter estimates from the generalized linear models (GLM) with a binomial error distribution testing for the effect of individual and maternal heterozygosity (sMLH) on pup survival (a) including maternal genetic diversity and mother age, and (b) excluding maternal effects. Log-odd ratios are shown together with confidence intervals (CI), significant *p*-values are in bold. Total number of observations, as well as the variance explained (R^2^ Tjur) are reported for both models.

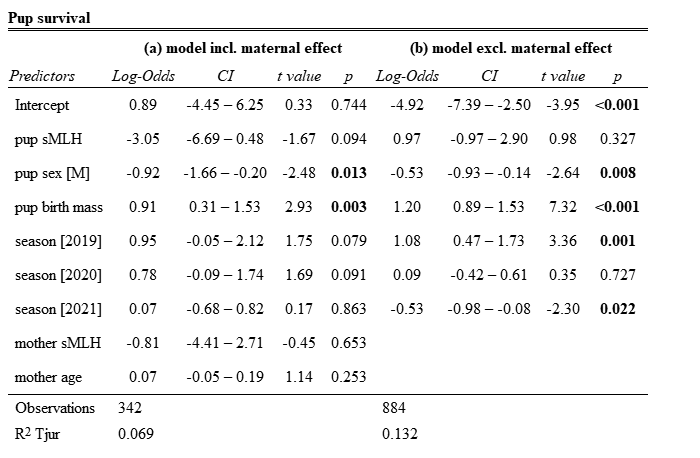

Table S3. Parameter estimates from the linear models testing for the effect of individual and maternal heterozygosity (sMLH) on pup growth (a) including maternal genetic diversity and mother age, and (b) excluding maternal effects. Estimates are shown together with confidence intervals (CI), significant *p*-values are in bold. For both models, total number of observations, as well as the variance explained by the predictors (R^2^) and variance adjusted for the number of predictors (R^2^ adjusted) are reported.

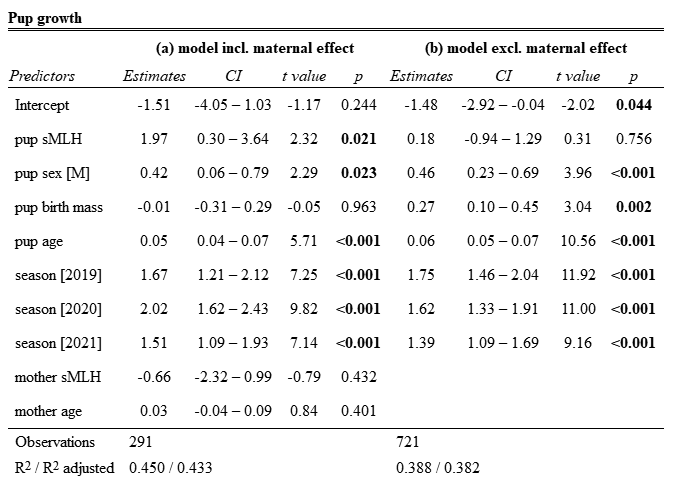

Table S4. Parameter estimates from the linear model testing for the effect of individual and maternal inbreeding (*F*_ROH_) on pup birth mass. Estimates are shown together with confidence intervals (CI), significant *p*-values are in bold. Total number of observations, as well as the variance explained by the predictors (R^2^) and variance adjusted for the number of predictors (R^2^ adjusted) are given.

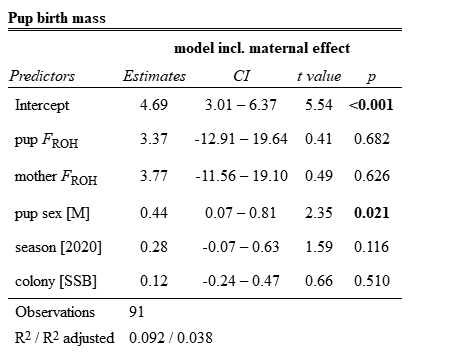

Table S5. Parameter estimates from the generalized linear model (GLM) with a binomial error distribution testing for the effect of individual and maternal inbreeding (*F*_ROH_) on pup survival. Estimates are shown together with confidence intervals (CI), significant *p*-values are in bold. Total number of observations, as well as the variance explained (R^2^ Tjur) are given.

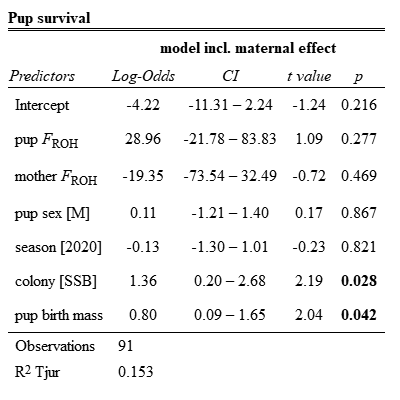

Table S6. Parameter estimates from the linear model testing for the effect of individual and maternal inbreeding (*F*_ROH_) on growth. Estimates are shown together with confidence intervals (CI), significant *p*-values are in bold. Total number of observations, as well as the variance explained by the predictors (R^2^) and variance adjusted for the number of predictors (R^2^ adjusted) are given.

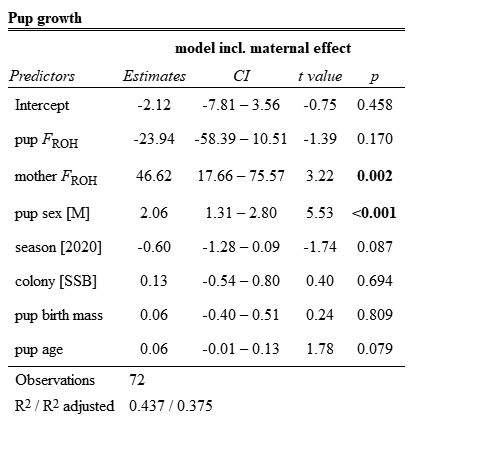
