## Supplementary material for "Little evidence for inbreeding depression for birth mass, survival and growth in Antarctic fur seal pups": R markdown file

Compiled by A.J. Paijmans and A.L. Berthelsen

12 January, 2024

#### Contents

**Packages and libraries** **2**

**Microsatellite dataset** **3**
Maternity check . . . 3
Data preparation . . . 3

**SNP dataset** **5**

**Variance in inbreeding** **5**

**Statistical models - microsatellite heterozygosity** **8**

---

This document contains all the `R code` used in the workflow for the manuscript *Little evidence for inbreeding depression for birth mass, survival and growth in Antarctic fur seal pups* by Anneke J. Paijmans, Ane Liv Berthelsen, Rebecca Nagel, Felicitas Cristaller, Nicole Kröcker, Jaume Forcada, and Joseph I. Hoffman. The R Markdown file and the raw data can all be downloaded via Zenodo, *LINK*. Additional scrips are available on Github, https://github.com/apaijmans/inbreeding-pup-growth/. Please, don't hesitate to contact me if you have any questions: a.paijmans[@]uni-bielefeld.de.

---

### Packages and libraries

```
packages <- c("here",
              "readxl",
              "tidyverse",
              "inbreedR",
              "lme4",
              "lmerTest",
              "DHARMa",
              "sjPlot",
              "cowplot",
              "ggtext",
              "nlme",
              "car",
              "AICcmodavg",
              "data.table",
              "rcartocolor",
              "dotwhisker",
              "merDeriv",
              "sf",
              "chron",
              "ggspatial",
              "ggsn")

# # Install packages needed not yet installed
# installed_packages <- packages %in% rownames(installed.packages())
#   if (any(installed_packages == FALSE)) {
```

```r
#   install.packages(packages[!installed_packages])
#   }

# Load packages
invisible(lapply(packages, library, character.only = TRUE))
```

### Microsatellite dataset

#### Maternity check

Before starting the analysis, mother-pup pairs were checked for genetic matching using the excel NEWPAT macro. The genotypes of pairs with a maximum of 3 mismatching loci were visually inspected. If a scoring mistake was identified, the genotype was corrected. The maternity analysis was then rerun on the updated microsatellite data. Mother-pup pairs with a maximum of 1 mismatching loci were considered a genetic match.

Preparing the files for the maternity analysis and postprocessing of the results were done in separate R scripts. These scripts and the NEWPAT excel macro files can be found on Github; scripts 1a-1e.

#### Data preparation

First, we load the fitness data including the results of the maternity analysis, and the microsatellite data for all our individuals in two separate dataframes.

```r
#~~ Load pup data
pup_data <- openxlsx::read.xlsx(here("Data", "Processed",
 ↪  "pup_growth_mums_checked.xlsx"), detectDates = T) %>%
  mutate(Pup_Sex = as.factor(Pup_Sex)) %>%
  mutate(Year = as.factor(Year)) %>%
  mutate(Age_Tag = as.integer(Age_Tag))
```

```r
#~~ Load msats for all individuals in this study
msats_all <- openxlsx::read.xlsx(here("Data", "Processed",
 ↪  "msats_growth_individuals.xlsx"), detectDates = T) %>%
  mutate(Gaps = as.numeric(Gaps))
```

Genotypes with more than 4 missing loci out of 39 were removed.

```r
#~~ Keep only individuals genotyped for all 39 loci, with a maximum of 4 missing loci
msats_all <- msats_all %>%
  filter(Gaps < 5)
```

We then calculated sMLH for all remaining individuals and added this to the fitness data.

```r
#~~ Convert msat data to inbreedR input
msat_genotypes39 <-  msats_all %>% select(Pv9.a:Mang36.b)
msat_ids <- msats_all %>% select(uniqueID)

msat_genotypes39_raw <- convert_raw(msat_genotypes39)

#~~ Check if data is in right format for InbreedR
check_data(msat_genotypes39_raw, num_ind = nrow(msat_ids), num_loci =
 ↪  length(msat_genotypes39_raw))
```

```
## [1] TRUE
```

```r
#~~ Calculate sMLH (incl FWB)
het39 <- sMLH(msat_genotypes39_raw)

sMLH_msat39 <- cbind(msat_ids, het39)
colnames(sMLH_msat39) <- c("uniqueID", "sMLH_msat39")

#~~ Add sMLH to pup data
pup_data <- left_join(pup_data, sMLH_msat39, by = c("uniqueID_pup" = "uniqueID")) %>%
  rename(sMLH_msat39_pup=sMLH_msat39)

pup_data <- left_join(pup_data, sMLH_msat39, by = c("uniqueID_mum" = "uniqueID"))%>%
  rename(sMLH_msat39_mum=sMLH_msat39)
```

We then removed all mothers and maternal information for the mothers that were not a genetic match (ie more than 1 mismatching locus).

```r
#~~ Remove mums that are not a genetic match, but keep pups (we can still use them in the
↪  analysis without mums)
# This means we also have to remove mum birth year and sMLH for the mismatching mums
pup_data <- pup_data %>%
  mutate(ID_Mum = ifelse(gen_mum == "match", ID_Mum, NA)) %>%
  mutate(uniqueID_mum = ifelse(gen_mum == "match", uniqueID_mum, NA)) %>%
  mutate(MumBirthYear = ifelse(gen_mum == "match", MumBirthYear, NA)) %>%
  mutate(Mum_Age = ifelse(gen_mum == "match", Mum_Age, NA)) %>%
  mutate(sMLH_msat39_mum = ifelse(gen_mum == "match", sMLH_msat39_mum, NA)) %>%
  select(-c(gen_mum, MISMATCHES))
```

And finally added a column with survival information (1 = survived until tagging, 0 = died).

```r
pup_data <- pup_data %>%
  mutate(Survival = ifelse(!is.na(Cat_Death) | !is.na(Pup_Death), "0",
                           ifelse(!is.na(Pup_TagWeight) & !is.na(Pup_Death), "0",
                                  ifelse(!is.na(Pup_TagWeight) & is.na(Pup_Death), "1",
                                   ↪  NA)))) %>%
  mutate(Survival = as.factor(Survival))

# All pups that do not have a tagging weight AND also no death date will now have an NA
↪  in the Survival column.
# These will be removed: we wont be able to use them in the growth model nor the survival
↪  model (as we do not know whether they were dead or alive).
# So for consistency, we will also not use them in the birth mass model (n=231)
# nrow(pup_data %>% filter(is.na(Survival)))
# All pups with a 2nd weight and no death date are assumed to have survived at least
↪  until the end of the season

pup_data <- pup_data %>%
  filter(!is.na(Survival))
```

Making sure all variables have the right categories

```r
#~~ Fix column categories
pup_data <- pup_data %>%
  mutate(Pup_Sex = as.factor(Pup_Sex)) %>%
  mutate(Year = as.factor(Year)) %>%
```

```
  mutate(Survival = as.factor(Survival))
```

### SNP dataset

Datasets for birth weight, survival and repeated measures growth analysis. Scripts for SNP data filtering, calling ROH and calculating $F_{\text{ROH}}$ are available on Github.

```
DataRM_Day60 <- read.csv(here("Data", "Raw", "GrowthRM_BI1820_Day60.new.csv"))

Unique_Day60 <- subset(DataRM_Day60, DataRM_Day60$Age_Days < 62 &
                         !DataRM_Day60$ID == 'H2' &
                         !DataRM_Day60$ID == 'H5') %>%
  mutate(Sex = as.factor(Sex)) %>%
  mutate(Season = as.factor(Season)) %>%
  mutate(Beach = as.factor(Beach)) %>%
  distinct(ID, .keep_all = TRUE)
#98 pups

UniqueSurvivors_Day60 <- subset(Unique_Day60, Unique_Day60$Death == 'N')
#76 pups

SurvivorsRM_Day60 <- subset(DataRM_Day60, DataRM_Day60$Death == 'N' &
                              DataRM_Day60$Age_Days < 62 &
                              !DataRM_Day60$ID == 'H2' &
                              !DataRM_Day60$ID == 'H5') %>%
  mutate(Sex = as.factor(Sex)) %>%
  mutate(Season = as.factor(Season)) %>%
  mutate(Beach = as.factor(Beach)) %>%
  mutate(Age_Days = as.numeric(Age_Days))
```

### Variance in inbreeding

To explore the variance in inbreeding, we calculated $g_2$ (n permumations = 1000, n bootstraps = 1000). Since the $g_2$ calculations can take some time, we calculated it once, stored it as a Gdata object and load the data from the saved object. The code to calculate the $g_2$ can be found on Github for the microsatellites and SNP data, as well as the resulting Rdata objects.

```
load(here("Data", "Processed", "g2_39loci_after_checks.RData"))

# summary
g2_39loci
```

```
##
##
## Calculation of identity disequilibrium with g2 for microsatellite data
## ----------------------------------------------------------------------
##
## Data: 1491 observations at 39 markers
## Function call = g2_microsats(genotypes = msat_genotypes39_raw, nperm = 1000,      nboot = 1000, CI = 
##
## g2 = 0.0005367851, se = 0.0003419581
##
```

```
## confidence interval
##          2.5%         97.5%
## -0.0001036011  0.0012367364
##
## p (g2 > 0) = 0.04 (based on 1000 permutations)
```

```r
# SNP g2 values (file generated in "calculate_g2.R" script that was ran on the server)
load(here("Data", "Processed", "g2_snps.RData"))

# summary SNPs
g2_snp_geno
```

```
##
##
## Calculation of identity disequilibrium with g2 for SNP data
## ----------------------------------------------------------
##
## Data: 196 observations at 75101 markers
## Function call = g2_snps(genotypes = snp_genotypes, nperm = 1000, nboot = 1000,     CI = 0.95, paralle
##
## g2 = 0.0001179302, se = 1.753435e-05
##
## confidence interval
##         2.5%        97.5%
## 8.597162e-05 1.523841e-04
##
## p (g2 > 0) = 0.001 (based on 999 permutations)
```

```r
# Make df for plotting
g2_plot <- rbind(data.frame(g2_boot = g2_snp_geno$g2_boot, gen_data = "snps"),
                 data.frame(g2_boot = g2_39loci$g2_boot, gen_data = "msats"))

lcl_snps <- g2_snp_geno$CI_boot[1]
ucl_snps <- g2_snp_geno$CI_boot[2]
g2_boot_summary_snps <- data.frame(lcl_snps, ucl_snps)

lcl_msat <- g2_39loci$CI_boot[1]
ucl_msat <- g2_39loci$CI_boot[2]
g2_boot_summary_msat <- data.frame(lcl_msat, ucl_msat)

# Colors
col1 <- "#872ca2"
col2 <- "#f6a97a"

# Use Martin Stoffel's GGplot theme as a base
source(here("Rcode", "anneke_theme.R"))

ggplot(g2_plot, aes(x = g2_boot, fill = gen_data)) +
  geom_histogram(alpha = 0.7, position = "identity", binwidth = 0.00001) + # 0.00001 or
  ↪  0.00005 ?
  scale_fill_manual(values = c(col1, col2), labels = c("msats\n(n = 1491)", "SNPs\n(n =
  ↪  196)")) +
  # Add CI bars and g2 line for msats
  geom_errorbarh(aes(xmin = g2_boot_summary_msat$lcl_msat , xmax =
  ↪  g2_boot_summary_msat$ucl_msat , y = 250),
```

```
                 linewidth = 0.8, color = col1, linetype = "solid", height = 0) +
  geom_vline(xintercept = g2_39loci$g2, linewidth = 0.6, color = col1, linetype =
   ↪  "dashed") +
  # Add CI bars and g2 line for SNPs
  geom_errorbarh(aes(xmin = g2_boot_summary_snps$lcl_snps , xmax =
   ↪  g2_boot_summary_snps$ucl_snps , y = 255),
                 linewidth = 0.8, color = col2, linetype = "solid", height = 0) +
  geom_vline(xintercept = g2_snp_geno$g2, linewidth = 0.6, color = col2, linetype =
   ↪  "dashed") +
  # Add zero line
  geom_vline(xintercept = 0, linewidth = 0.6, linetype = "solid") +
  # Add other labs and theme
  labs(y = "Counts", x = expression(italic(g[2]))) +
  theme_anneke() +
  theme(legend.title=element_blank())
```

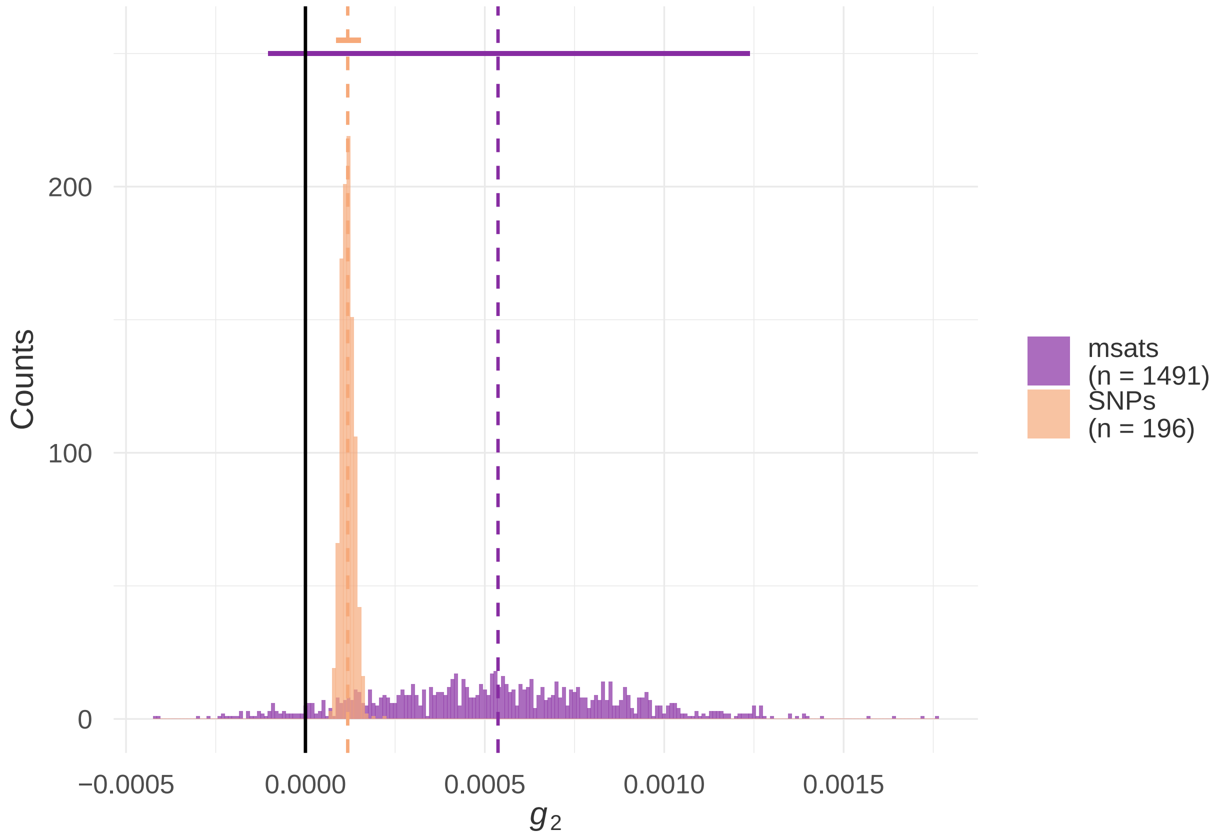

We then plotted the bootstrapped $g_2$ values for both the microsatellite data as well as the SNP array data. For both datasets, the $g_2$ was significantly different from zero (microsatellite data: $p = 0.04$, SNP array data: $p = 0.001$).

### Statistical models - microsatellite heterozygosity

#### Microsatellite dataset: pup birth mass

We tested for an effect of individual or maternal heterozygosity (sMLH) on pup birth mass with a linear model. Pup and maternal sMLH were included as continuous variables. Covariates were pup sex and year (as factors) and mother age (as continuous variable) (see Model 1).

To make use of a bigger sample size (including pups with known and unknown mothers) we ran the same model while excluding maternal effects (see Model 2). The results for the predictors of main interest (pup sMLH/mum sMLH) did not differ (see parameter estimates)). In the model without maternal effects there seemed to be a year effect for 2020 and 2021 that was not present in the model including maternal effects.

##### Model 1: including maternal effects

```
#~~ Birth mass model incl maternal effects
m1birthmass <- lm(Pup_BirthWeight ~ sMLH_msat39_pup
                  + Pup_Sex
                  + Year
                  + sMLH_msat39_mum
                  + Mum_Age,
                  # + (1 | uniqueID_mum), # including mother ID as a random effect did
                   ↪  not change the results significantly and the model was a poorer fit
                  # In addition, adding mother ID as random effect may make it
                   ↪  problematic as sMLH mum would be a confounding factor.
                  data = pup_data)

summary(m1birthmass)
```

```
##
## Call:
## lm(formula = Pup_BirthWeight ~ sMLH_msat39_pup + Pup_Sex + Year +
##     sMLH_msat39_mum + Mum_Age, data = pup_data)
##
## Residuals:
##      Min       1Q   Median       3Q      Max
## -2.40510 -0.35600  0.00611  0.28943  1.78262
##
## Coefficients:
##                  Estimate Std. Error t value Pr(>|t|)
## (Intercept)       4.27721    0.40642  10.524  < 2e-16 ***
## sMLH_msat39_pup -0.03999    0.31888  -0.125 0.900272
## Pup_SexM         0.64099    0.05660  11.324  < 2e-16 ***
## Year2018        -0.22695    0.08521  -2.663 0.008108 **
## Year2019         0.09053    0.07463   1.213 0.225924
## Year2020        -0.13158    0.07470  -1.761 0.079073 .
## sMLH_msat39_mum  0.04050    0.32059   0.126 0.899539
## Mum_Age          0.03751    0.01059   3.541 0.000455 ***
## ---
## Signif. codes:  0 '***' 0.001 '**' 0.01 '*' 0.05 '.' 0.1 ' ' 1
##
## Residual standard error: 0.5193 on 334 degrees of freedom
##   (742 observations deleted due to missingness)
## Multiple R-squared:  0.3291, Adjusted R-squared:  0.315
## F-statistic:  23.4 on 7 and 334 DF,  p-value: < 2.2e-16
```

##### Residual check of model

We used the DHARMa package to check three model assumptions: the first figure shows a test for under/overdispersion, the second figure, a QQplot, checks for normality, and the third figure, residuals versus the predictions, allows to check for issues with linearity and equality of error variances. Since the outlier test was significant (meaning that the simulated values are higher or lower than the observed data), we investigated these in the fourth figure.

```
#~~ Model assumptions
testDispersion(m1birthmass) # tests for over/underdispersion
```

```
##
##  DHARMa nonparametric dispersion test via sd of residuals fitted vs.
##  simulated
##
## data:  simulationOutput
## dispersion = 0.97627, p-value = 0.712
## alternative hypothesis: two.sided
```

```
plotQQunif(m1birthmass) # detect overall deviations from the expected distribution. Is
↪  data normally distributed?
plotResiduals(m1birthmass) # plot residuals against the predicted value
# Outlier test is significant, although the QQplot looks good

#~~ Further investigation of the outliers
simulationOutput <- simulateResiduals(fittedModel = m1birthmass)
testOutliers(simulationOutput)
```

```
##
##  DHARMa outlier test based on exact binomial test with approximate
##  expectations
##
## data:  simulationOutput
## outliers at both margin(s) = 7, observations = 342, p-value = 0.021
## alternative hypothesis: true probability of success is not equal to 0.007968127
## 95 percent confidence interval:
##  0.008267784 0.041715094
## sample estimates:
## frequency of outliers (expected: 0.00796812749003984 )
##                                              0.02046784
```

```
# There are 7 outliers out of 342 obs. The frequency of outliers is significantly higher
↪  than expected (2.04% against 0.79% expected)
# According to the vignette, a certain number of outliers are expected 'at random'. It
↪  could also hint at overdispersion, but the test for dispersion
# showed no indication for that. The DHARMa vignette also mentions that the visual
↪  indicators of the residuals are a lot more informative than the p-values.
# Taken together with the fact that all plots look good, it is not likely to be a
↪  problem.
```

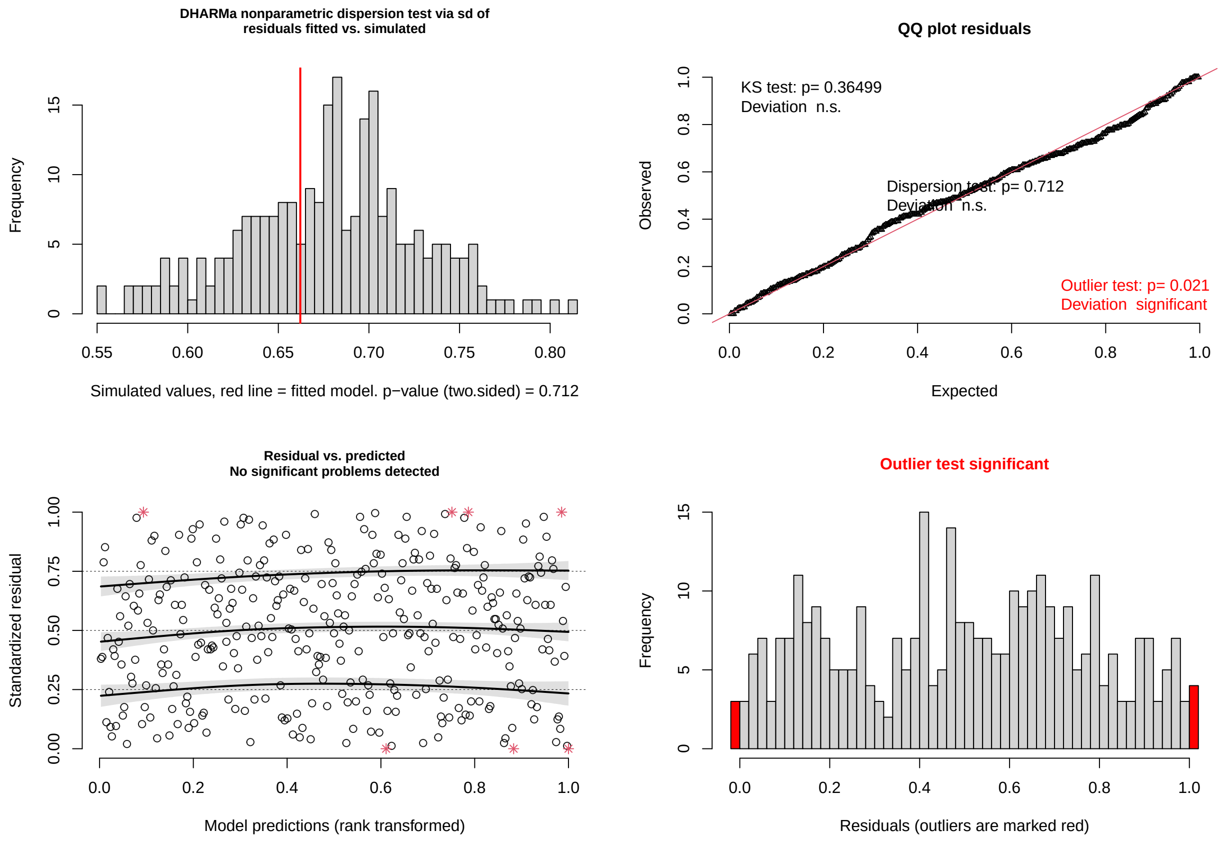

##### Model 2: excluding maternal effects

```r
#~~ Birth mass model excl maternal effects
m2birthmass <- lm(Pup_BirthWeight ~ sMLH_msat39_pup
                  + Pup_Sex
                  + Year,
                  data = pup_data)

summary(m2birthmass)
```

```
##
## Call:
## lm(formula = Pup_BirthWeight ~ sMLH_msat39_pup + Pup_Sex + Year,
##     data = pup_data)
##
## Residuals:
##     Min      1Q  Median      3Q     Max
## -2.5190 -0.3705 -0.0043  0.3824  2.3107
##
## Coefficients:
##                 Estimate Std. Error t value Pr(>|t|)
## (Intercept)      4.53703    0.23306  19.467  < 2e-16 ***
## sMLH_msat39_pup  0.12353    0.22927   0.539  0.59016
## Pup_SexM         0.55663    0.04228  13.164  < 2e-16 ***
## Year2018        -0.19440    0.06122  -3.175  0.00155 **
```

```
## Year2019         0.15770    0.05729   2.753  0.00603 **
## Year2020        -0.25740    0.05724  -4.497 7.83e-06 ***
## ---
## Signif. codes:  0 '***' 0.001 '**' 0.01 '*' 0.05 '.' 0.1 ' ' 1
##
## Residual standard error: 0.6274 on 878 degrees of freedom
##   (200 observations deleted due to missingness)
## Multiple R-squared:  0.2106, Adjusted R-squared:  0.2061
## F-statistic: 46.85 on 5 and 878 DF,  p-value: < 2.2e-16
```

##### Residual check of model

```r
#~~ Model assumptions
testDispersion(m2birthmass)
```

```
##
##  DHARMa nonparametric dispersion test via sd of residuals fitted vs.
##  simulated
##
## data:  simulationOutput
## dispersion = 0.99112, p-value = 0.888
## alternative hypothesis: two.sided
```

```r
plotQQunif(m2birthmass)
plotResiduals(m2birthmass)

# Outlier test is significant, although the QQplot looks good
#~~ Further investigation of the outliers
simulationOutput <- simulateResiduals(fittedModel = m2birthmass)
testOutliers(simulationOutput)
```

```
##
##  DHARMa outlier test based on exact binomial test with approximate
##  expectations
##
## data:  simulationOutput
## outliers at both margin(s) = 14, observations = 884, p-value = 0.01997
## alternative hypothesis: true probability of success is not equal to 0.007968127
## 95 percent confidence interval:
##  0.008684635 0.026429330
## sample estimates:
## frequency of outliers (expected: 0.00796812749003984 )
##                                             0.0158371
```

```r
# There are 14 outliers out of 884 obs.
# Also here, the test for dispersion shows no indication for overdispersion, and the
↪ QQplot looks good.
# Therefore, the model fit is likely to be OK.
```

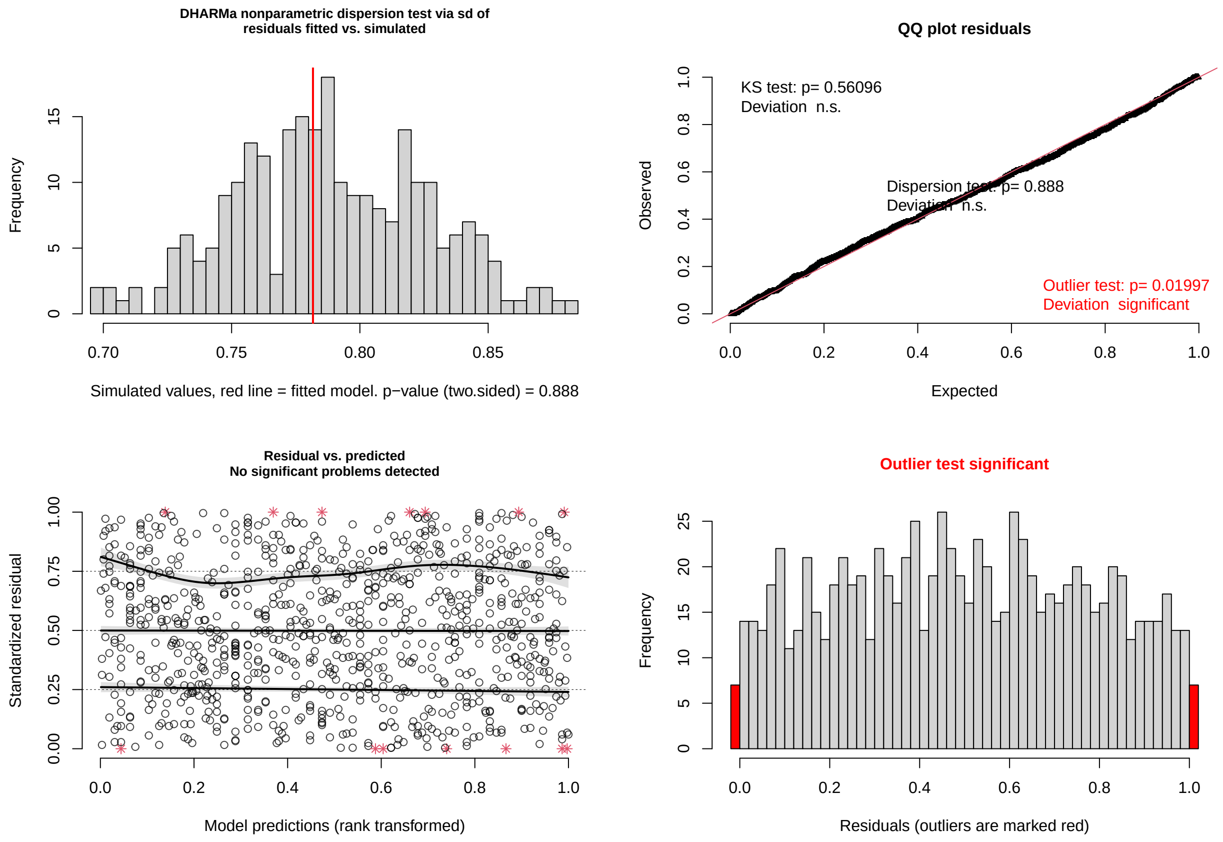

#### Parameter estimates pup birth mass models

Parameter estimates from: (a) the linear model including maternal genetic diversity (mother sMLH) and mother age, and (b) the linear model excluding maternal effects. Estimates are shown together with confidence intervals (CI), significant *p*-values are in bold. For both models, total number of observations, as well as the variance explained by the predictors ($R^2$) and variance adjusted for the number of predictors ($R^2$ adjusted) are reported. The results of the model including maternal effect (a) compared to the results of the model excluding maternal effects (b) are very similar, even though the sample size is more than double (342 vs. 884, respectively).

```
# Statistical table showing parameter estimates for both models
# Labels
tab_label <- c(
  `(Intercept)` = "Intercept",
  sMLH_msat39_pup = "pup sMLH",
  Pup_SexM = "pup sex [M]",
  sMLH_msat39_mum = "mother sMLH",
  Mum_Age = "mother age",
  Year2018 = "season [2019]",
  Year2019 = "season [2020]",
  Year2020 = "season [2021]")

# Table
# print so it saves but doesn't show the html table,
# which doesn't display nicely in the pdf generated by Rmarkdown
print(tab_model(m1birthmass, m2birthmass,
```

```
                 pred.labels = tab_label,
                 title = "Pup birth mass",
                 dv.labels = c("(a) model incl. maternal effect",
                               "(b) model excl. maternal effect"),
                 show.stat=T,
                 string.stat = "t value",
                 file = here("Tables", "Table_BW_full_model_vs_no_mat_NEW.html")))

# Make a screenshot of saved htlm table and save as a png
# so that it can be shown in Rmarkdown pdf
webshot::webshot(here("Tables", "Table_BW_full_model_vs_no_mat_NEW.html"),
                 file=here("Tables", "Table_BW_full_model_vs_no_mat_NEW.png"), delay=2,
                  ↪  vheight = 450, vwidth = 700)
```

**Pup birth mass**

| | **(a) model incl. maternal effect** | | | | **(b) model excl. maternal effect** | | | |
|---|---|---|---|---|---|---|---|---|
| *Predictors* | *Estimates* | *CI* | *t value* | *p* | *Estimates* | *CI* | *t value* | *p* |
| Intercept | 4.28 | 3.48 – 5.08 | 10.52 | **<0.001** | 4.54 | 4.08 – 4.99 | 19.47 | **<0.001** |
| pup sMLH | -0.04 | -0.67 – 0.59 | -0.13 | 0.900 | 0.12 | -0.33 – 0.57 | 0.54 | 0.590 |
| pup sex [M] | 0.64 | 0.53 – 0.75 | 11.32 | **<0.001** | 0.56 | 0.47 – 0.64 | 13.16 | **<0.001** |
| season [2019] | -0.23 | -0.39 – -0.06 | -2.66 | **0.008** | -0.19 | -0.31 – -0.07 | -3.18 | **0.002** |
| season [2020] | 0.09 | -0.06 – 0.24 | 1.21 | 0.226 | 0.16 | 0.05 – 0.27 | 2.75 | **0.006** |
| season [2021] | -0.13 | -0.28 – 0.02 | -1.76 | 0.079 | -0.26 | -0.37 – -0.15 | -4.50 | **<0.001** |
| mother sMLH | 0.04 | -0.59 – 0.67 | 0.13 | 0.900 | | | | |
| mother age | 0.04 | 0.02 – 0.06 | 3.54 | **<0.001** | | | | |
| Observations | 342 | | | | 884 | | | |
| $R^2$ / $R^2$ adjusted | 0.329 / 0.315 | | | | 0.211 / 0.206 | | | |

#### Microsatellite dataset: survival

Here, we tested for an effect of individual or maternal heterozygosity on pup survival with a generalized linear model (GLM) with a binomial error structure. Survival ( 1 = survived and 0 = dead) was included as a factor, pup and maternal sMLH were included as continuous variables. Also in this model the additional covariates were pup sex, year (as factors) and mother age (as continuous variable) (see Model 1).

Again, we compared the results with the results generated by running the same model on the bigger dataset (including pups with known and unknown mothers) while excluding maternal effects (see Model 2). The results for the predictors of main interest (pup sMLH/mum sMLH) did not differ (see parameter estimates)). In the model without maternal effects there seemed to be a year effect that was not present in the model including maternal effects. Pups were more likely to survive in 2019 compared to 2018 but less likely to survive in 2021 compared to 2018.

##### Model 1: including maternal effects

```r
#~~ Survival incl maternal effects
m1survival <- glm(Survival ~ sMLH_msat39_pup
                  + Pup_Sex
                  + Pup_BirthWeight
                  + Year
                  + sMLH_msat39_mum
                  + Mum_Age,
                  #+ (1 | uniqueID_mum), # including mother ID as a random effect did not
                   ↪ change the results significantly, and might be confounded with mum
                   ↪ sMLH
                  family=binomial,
                  data = pup_data)
# # Because of convergence issues for the model incl mother ID as a random effect, I had
 ↪ to re-run it with more iterations
#
# ss <- getME(m1survival, c("theta", "fixef"))
# m1survival.m2 <- update(m1survival, start = ss, control = glmerControl(optimizer =
 ↪ "bobyqa", optCtrl = list(maxfun = 2e+05)))

summary(m1survival) # conclusions are the same as for the model excl mum ID as a random
 ↪ factor
```

```
##
## Call:
## glm(formula = Survival ~ sMLH_msat39_pup + Pup_Sex + Pup_BirthWeight +
##     Year + sMLH_msat39_mum + Mum_Age, family = binomial, data = pup_data)
##
## Deviance Residuals:
##     Min       1Q   Median       3Q      Max
## -2.4790   0.3314   0.4559   0.5976   1.1730
##
## Coefficients:
##                 Estimate Std. Error z value Pr(>|z|)
## (Intercept)      0.88722    2.72121   0.326  0.74439
## sMLH_msat39_pup -3.05224    1.82370  -1.674  0.09420 .
## Pup_SexM        -0.92138    0.37083  -2.485  0.01297 *
## Pup_BirthWeight  0.90808    0.30995   2.930  0.00339 **
## Year2018         0.94972    0.54154   1.754  0.07947 .
## Year2019         0.77948    0.46104   1.691  0.09090 .
## Year2020         0.06544    0.38034   0.172  0.86339
## sMLH_msat39_mum -0.81441    1.81066  -0.450  0.65287
## Mum_Age          0.06996    0.06123   1.143  0.25323
## ---
## Signif. codes:  0 '***' 0.001 '**' 0.01 '*' 0.05 '.' 0.1 ' ' 1
##
## (Dispersion parameter for binomial family taken to be 1)
##
##     Null deviance: 288.09  on 341  degrees of freedom
## Residual deviance: 264.90  on 333  degrees of freedom
##   (742 observations deleted due to missingness)
## AIC: 282.9
##
```

```
## Number of Fisher Scoring iterations: 5
```

##### Residual check of model

```r
#~~ Model assumptions
testDispersion(m1survival)
```

```
##
##  DHARMa nonparametric dispersion test via sd of residuals fitted vs.
##  simulated
##
## data:  simulationOutput
## dispersion = 1.0157, p-value = 0.824
## alternative hypothesis: two.sided
```

```r
plotQQunif(m1survival)
plotResiduals(m1survival)
```

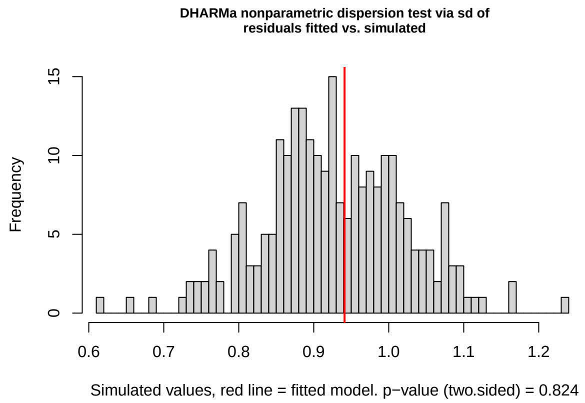

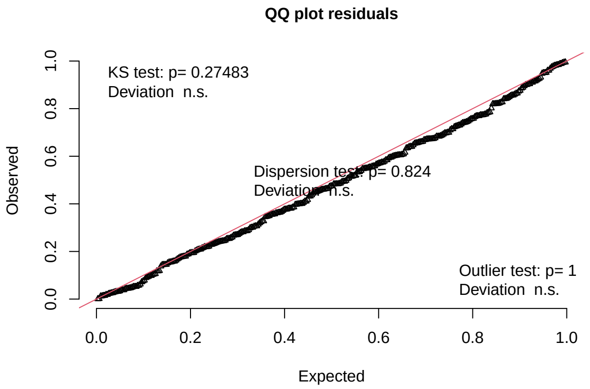

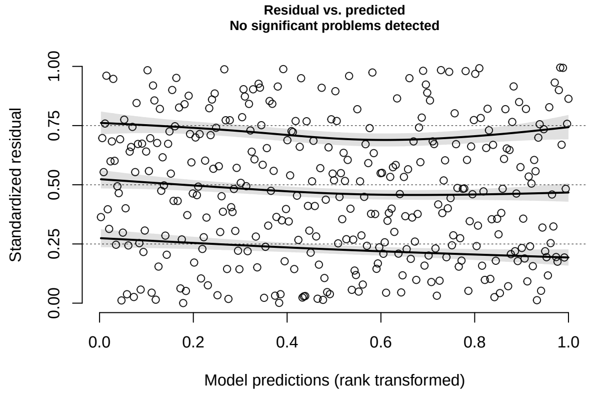

##### Model 2: excluding maternal effects

```r
#~~ Survival excl maternal effects
m2survival <- glm(Survival ~ sMLH_msat39_pup
                  + Pup_Sex
                  + Pup_BirthWeight
                  + Year,
                  family=binomial,
```

```r
                   data = pup_data)

summary(m2survival)
```

```
##
## Call:
## glm(formula = Survival ~ sMLH_msat39_pup + Pup_Sex + Pup_BirthWeight +
##     Year, family = binomial, data = pup_data)
##
## Deviance Residuals:
##     Min       1Q   Median       3Q      Max
## -2.6930   0.2944   0.4941   0.6493   1.4802
##
## Coefficients:
##                   Estimate Std. Error z value Pr(>|z|)
## (Intercept)        -4.9198     1.2444  -3.954 7.69e-05 ***
## sMLH_msat39_pup     0.9654     0.9857   0.979 0.327359
## Pup_SexM           -0.5317     0.2017  -2.636 0.008380 **
## Pup_BirthWeight     1.2004     0.1640   7.319 2.49e-13 ***
## Year2018            1.0788     0.3212   3.358 0.000785 ***
## Year2019            0.0915     0.2624   0.349 0.727332
## Year2020           -0.5269     0.2294  -2.297 0.021634 *
## ---
## Signif. codes:  0 '***' 0.001 '**' 0.01 '*' 0.05 '.' 0.1 ' ' 1
##
## (Dispersion parameter for binomial family taken to be 1)
##
##     Null deviance: 845.08  on 883  degrees of freedom
## Residual deviance: 745.04  on 877  degrees of freedom
##   (200 observations deleted due to missingness)
## AIC: 759.04
##
## Number of Fisher Scoring iterations: 5
```

##### Residual check of model

```r
#~~ Model assumptions
testDispersion(m2survival)
```

```
##
##  DHARMa nonparametric dispersion test via sd of residuals fitted vs.
##  simulated
##
## data:  simulationOutput
## dispersion = 0.9758, p-value = 0.632
## alternative hypothesis: two.sided
```

```r
plotQQunif(m2survival)
plotResiduals(m2survival)
```

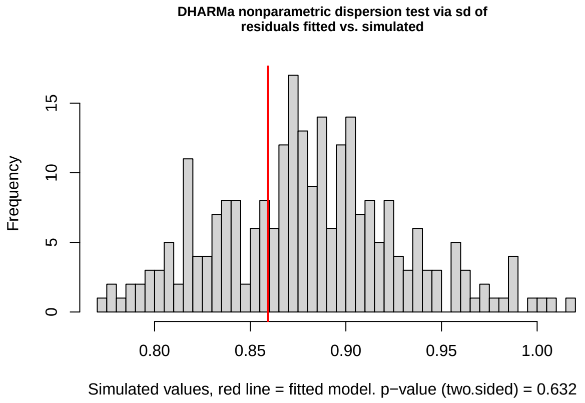

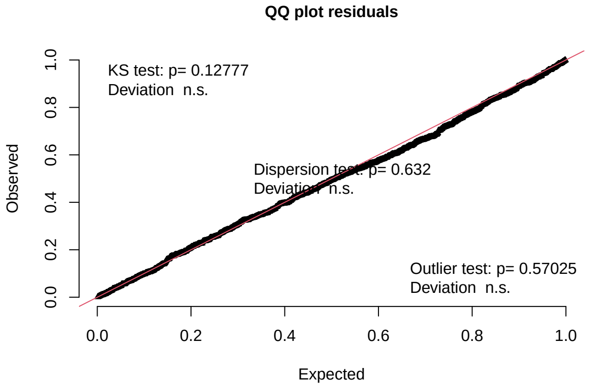

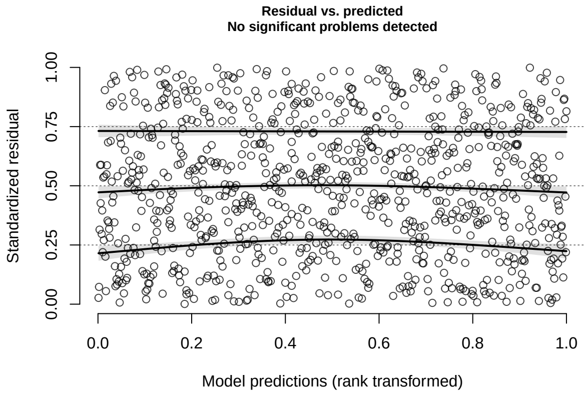

##### Parameter estimates pup survival models

Parameter estimates from: (a) the GLM with a binomial error distribution including maternal genetic diversity, and (b) excluding maternal effects. Log-Odd ratios are shown together with confidence intervals (CI), significant *p*-values are in bold. Total number of observations, as well as the variance explained ($R^2$ Tjur) are reported for both models. The results of the model including maternal effect (a) compared to the results of the model excluding maternal effects (b) are very similar, even though the sample size is more than double (342 vs. 884, respectively).

```
#~~ Labels
tab_labSurvival <- c(
  `(Intercept)` = "Intercept",
  sMLH_msat39_pup = "pup sMLH",
  Pup_BirthWeight = "pup birth mass",
  Pup_SexM = "pup sex [M]",
  sMLH_msat39_mum = "mother sMLH",
  Mum_Age = "mother age",
  Year2018 = "season [2019]",
  Year2019 = "season [2020]",
  Year2020 = "season [2021]")

#~~ Table
print(tab_model(m1survival, m2survival,
                pred.labels = tab_labSurvival,
                title = "Pup survival",
                dv.labels = c("(a) model incl. maternal effect",
```

```r
                              "(b) model excl. maternal effect"),
                  #order.terms = c(1, 2, 4, 3, 7, 5, 6),
                  transform = NULL,
                  show.stat=T,
                  string.stat = "t value",
                  file = here("Tables", "Table_survival_full_model_vs_no_mat.html")))

#~~ Makes a screenshot of saved htlm table and saves as a png
webshot::webshot(here("Tables", "Table_survival_full_model_vs_no_mat.html"),
                   file=here("Tables", "Table_survival_full_model_vs_no_mat.png"), delay=2,
                   ↪  vheight = 400, vwidth = 700)
```

**Pup survival**

| | (a) model incl. maternal effect | | | | (b) model excl. maternal effect | | | |
|---|---|---|---|---|---|---|---|---|
| *Predictors* | *Log-Odds* | *CI* | *t value* | *p* | *Log-Odds* | *CI* | *t value* | *p* |
| Intercept | 0.89 | -4.45 – 6.25 | 0.33 | 0.744 | -4.92 | -7.39 – -2.50 | -3.95 | **<0.001** |
| pup sMLH | -3.05 | -6.69 – 0.48 | -1.67 | 0.094 | 0.97 | -0.97 – 2.90 | 0.98 | 0.327 |
| pup sex [M] | -0.92 | -1.66 – -0.20 | -2.48 | **0.013** | -0.53 | -0.93 – -0.14 | -2.64 | **0.008** |
| pup birth mass | 0.91 | 0.31 – 1.53 | 2.93 | **0.003** | 1.20 | 0.89 – 1.53 | 7.32 | **<0.001** |
| season [2019] | 0.95 | -0.05 – 2.12 | 1.75 | 0.079 | 1.08 | 0.47 – 1.73 | 3.36 | **0.001** |
| season [2020] | 0.78 | -0.09 – 1.74 | 1.69 | 0.091 | 0.09 | -0.42 – 0.61 | 0.35 | 0.727 |
| season [2021] | 0.07 | -0.68 – 0.82 | 0.17 | 0.863 | -0.53 | -0.98 – -0.08 | -2.30 | **0.022** |
| mother sMLH | -0.81 | -4.41 – 2.71 | -0.45 | 0.653 | | | | |
| mother age | 0.07 | -0.05 – 0.19 | 1.14 | 0.253 | | | | |
| Observations | 342 | | | | 884 | | | |
| $R^2$ Tjur | 0.069 | | | | 0.132 | | | |

#### Microsatellite dataset: pup growth

Pup growth was defined as the difference in weight between birth and recapture. Pups that did not survive the first months were not recaptured, meaning weight at recapture was not available for these pups and growth could not be calculated. Because of this, we filtered the data to include only surviving pups.

We then tested for an effect of individual or maternal heterozygosity on pup survival with a linear model. Growth was included as a continuous variable. Addition covariates were pup age (number of days between birth and recapture), pup sex, year and mother age (see Model 1).

Again, here we compared the results with the results generated by running the same model on the bigger dataset (including pups with known and unknown mothers) while excluding maternal effects (see Model 2). The results were comparable between the two models (see parameter estimates)), although we found a significant effect of pup sMLH in the model with maternal effect, which is not present in the model without maternal effects. In the model without maternal effects there was an effect of pup birth mass, where pups that were born heavier also gained more weight, however, this effect was absent in the model with maternal effects.

```r
# Keep only pups that survived
pup_alife <- pup_data %>%
  filter(Survival == "1") %>%
  select(-c(Pup_Death, Survival))
```

##### Model 1: including maternal effects

```r
#~~ Weight gain incl maternal effects
m1growth <- lm(WeightGain ~ sMLH_msat39_pup
               + Pup_Sex
               + Pup_BirthWeight
               + Age_Tag
               + Year
               + sMLH_msat39_mum
               + Mum_Age,
               #+ (1 | uniqueID_mum), # including mother ID as a random effect did not
               ↪   change the results significantly, and might be confounded with mum
               ↪   sMLH
               data = pup_alife)

summary(m1growth)
```

```
##
## Call:
## lm(formula = WeightGain ~ sMLH_msat39_pup + Pup_Sex + Pup_BirthWeight +
##     Age_Tag + Year + sMLH_msat39_mum + Mum_Age, data = pup_alife)
##
## Residuals:
##     Min      1Q  Median      3Q     Max
## -4.6704 -0.7894  0.0427  0.8033  3.7300
##
## Coefficients:
##                  Estimate Std. Error t value Pr(>|t|)
## (Intercept)     -1.509259   1.291837  -1.168   0.2437
## sMLH_msat39_pup  1.966734   0.847586   2.320   0.0210 *
## Pup_SexM         0.423281   0.184809   2.290   0.0227 *
## Pup_BirthWeight -0.007078   0.152805  -0.046   0.9631
## Age_Tag          0.054574   0.009551   5.714 2.81e-08 ***
## Year2018         1.666965   0.230027   7.247 4.13e-12 ***
## Year2019         2.022889   0.206037   9.818  < 2e-16 ***
## Year2020         1.511597   0.211733   7.139 8.04e-12 ***
## sMLH_msat39_mum -0.662822   0.841433  -0.788   0.4315
## Mum_Age          0.027679   0.032891   0.842   0.4008
## ---
## Signif. codes:  0 '***' 0.001 '**' 0.01 '*' 0.05 '.' 0.1 ' ' 1
##
## Residual standard error: 1.277 on 281 degrees of freedom
##   (567 observations deleted due to missingness)
## Multiple R-squared:  0.4504, Adjusted R-squared:  0.4327
## F-statistic: 25.58 on 9 and 281 DF,  p-value: < 2.2e-16
```

##### Residual check of model

```r
#~~ Model assumptions
testDispersion(m1growth)
```

```
##
##  DHARMa nonparametric dispersion test via sd of residuals fitted vs.
##  simulated
##
## data:  simulationOutput
## dispersion = 0.96696, p-value = 0.712
## alternative hypothesis: two.sided
```

```r
plotQQunif(m1growth)
plotResiduals(m1growth)
```

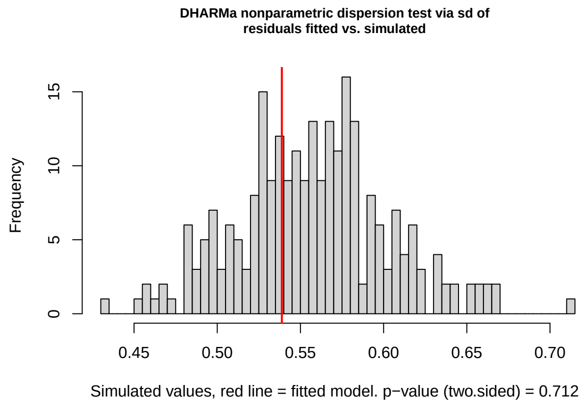

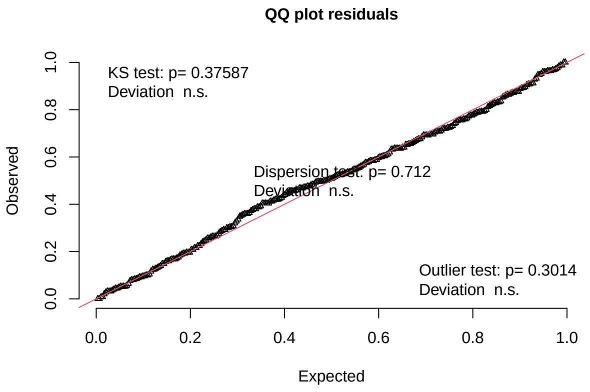

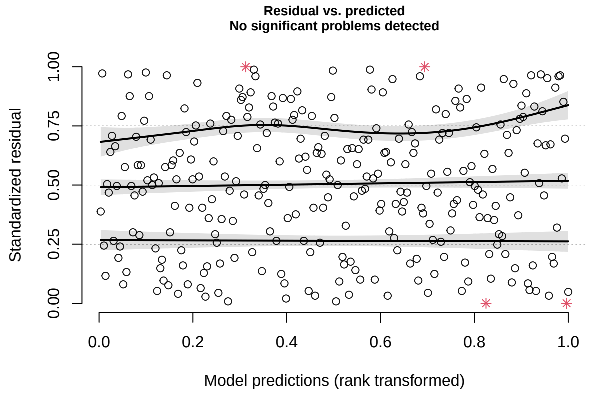

##### Model 2: excluding maternal effects

```r
#~~ Weight gain excl maternal effects
m2growth <- lm(WeightGain ~ sMLH_msat39_pup
                + Pup_Sex
                + Pup_BirthWeight
                + Age_Tag
                + Year,
                data = pup_alife)
```

```r
summary(m2growth)
```

```
##
## Call:
## lm(formula = WeightGain ~ sMLH_msat39_pup + Pup_Sex + Pup_BirthWeight +
##     Age_Tag + Year, data = pup_alife)
##
## Residuals:
##     Min      1Q  Median      3Q     Max
## -4.8136 -0.8687  0.0616  0.8039  4.6392
##
## Coefficients:
##                  Estimate Std. Error t value Pr(>|t|)
## (Intercept)     -1.476920   0.732784  -2.015  0.04423 *
## sMLH_msat39_pup  0.176693   0.569200   0.310  0.75633
## Pup_SexM         0.464347   0.117134   3.964 8.11e-05 ***
## Pup_BirthWeight  0.274592   0.090304   3.041  0.00245 **
## Age_Tag          0.055456   0.005252  10.559  < 2e-16 ***
## Year2018         1.753265   0.147036  11.924  < 2e-16 ***
## Year2019         1.617863   0.147068  11.001  < 2e-16 ***
## Year2020         1.390791   0.151884   9.157  < 2e-16 ***
## ---
## Signif. codes:  0 '***' 0.001 '**' 0.01 '*' 0.05 '.' 0.1 ' ' 1
##
## Residual standard error: 1.392 on 713 degrees of freedom
##   (137 observations deleted due to missingness)
## Multiple R-squared:  0.3884, Adjusted R-squared:  0.3823
## F-statistic: 64.67 on 7 and 713 DF,  p-value: < 2.2e-16
```

##### Residual check of model

```r
#~~ Model assumptions
testDispersion(m2growth)
```

```
##
##  DHARMa nonparametric dispersion test via sd of residuals fitted vs.
##  simulated
##
## data:  simulationOutput
## dispersion = 0.98774, p-value = 0.856
## alternative hypothesis: two.sided
```

```r
plotQQunif(m2growth)
plotResiduals(m2growth)
# The residuals vs. predicted quantile plot shows a small deviation for the 0.75
↪  quantile. However, the combined
# adjusted quantile test is not significant and also the Kolmogorov-Smirnov(KS)-test is
↪  non significant (see QQplot).
# So I conclude that the deviation is not very big, and no reason to reject the model.
```

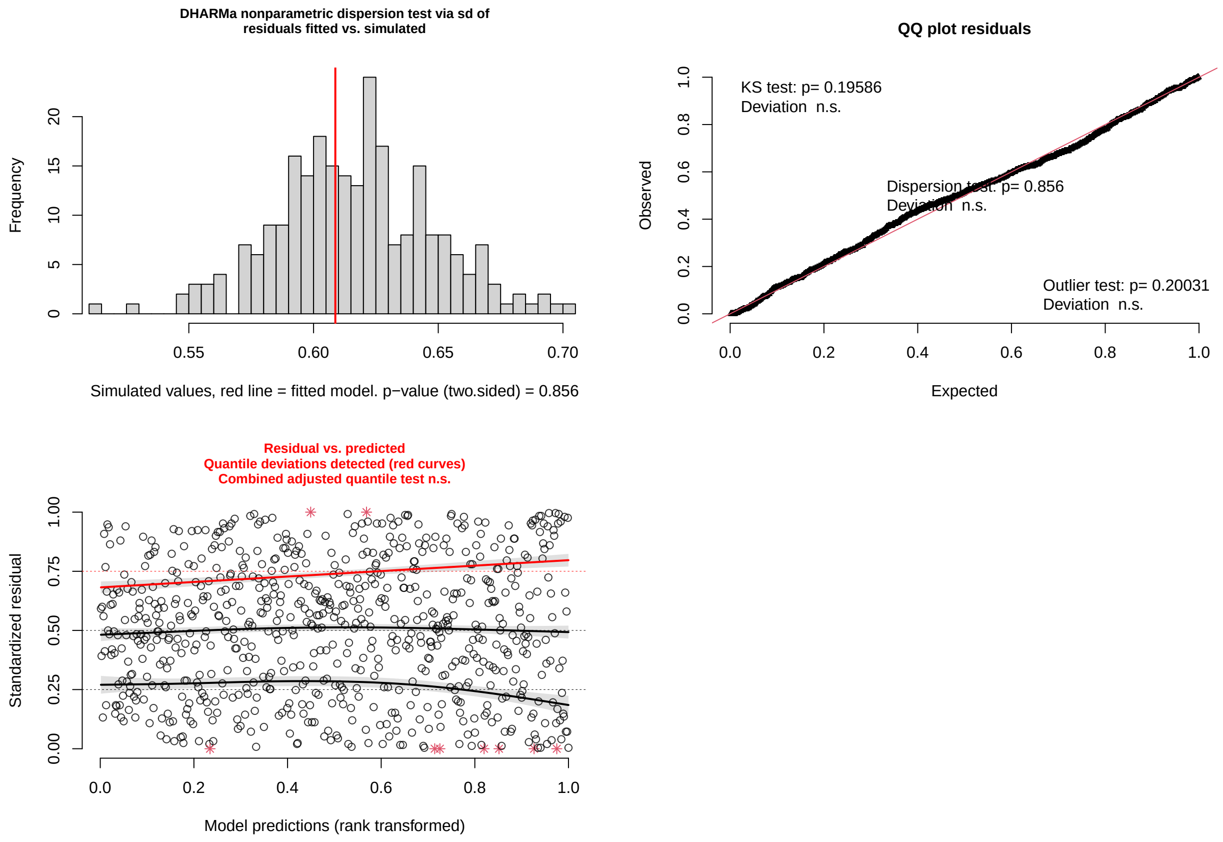

##### Parameter estimates pup growth models

Parameter estimates from: (a) linear model including maternal genetic diversity, and (b) excluding maternal effects. Estimates are shown together with confidence intervals (CI), significant *p*-values are in bold. For both models, total number of observations, as well as the variance explained by the predictors ($R^2$) and variance adjusted for the number of predictors ($R^2$ adjusted) are reported.

```
#~~ Labels
tab_labGrowth <- c(
  `(Intercept)` = "Intercept",
  sMLH_msat39_pup = "pup sMLH",
  Pup_BirthWeight = "pup birth mass",
  Pup_SexM = "pup sex [M]",
  sMLH_msat39_mum = "mother sMLH",
  Mum_Age = "mother age",
  Age_Tag = "pup age",
  Year2018 = "season [2019]",
  Year2019 = "season [2020]",
  Year2020 = "season [2021]")

#~~ Table
print(tab_model(m1growth, m2growth,
                pred.labels = tab_labGrowth,
                title = "Pup growth",
                dv.labels = c("(a) model incl. maternal effect",
                              "(b) model excl. maternal effect"),
```

```
                 #order.terms = c(1, 2, 3, 9, 4, 5, 6, 7, 8),
                 show.stat=T,
                 string.stat = "t value",
                 file = here("Tables", "Table_growth_full_model_vs_no_mat_NEW.html")))

# Makes a screenshot of saved htlm table and saves as a png
webshot::webshot(here("Tables", "Table_growth_full_model_vs_no_mat_NEW.html"),
                  file=here("Tables", "Table_growth_full_model_vs_no_mat_NEW.png"),
                   ↪  delay=2, vheight = 450, vwidth = 700)
```

**Pup growth**

| | **(a) model incl. maternal effect** | | | | **(b) model excl. maternal effect** | | | |
|---|---|---|---|---|---|---|---|---|
| *Predictors* | *Estimates* | *CI* | *t value* | *p* | *Estimates* | *CI* | *t value* | *p* |
| Intercept | -1.51 | -4.05 – 1.03 | -1.17 | 0.244 | -1.48 | -2.92 – -0.04 | -2.02 | **0.044** |
| pup sMLH | 1.97 | 0.30 – 3.64 | 2.32 | **0.021** | 0.18 | -0.94 – 1.29 | 0.31 | 0.756 |
| pup sex [M] | 0.42 | 0.06 – 0.79 | 2.29 | **0.023** | 0.46 | 0.23 – 0.69 | 3.96 | **<0.001** |
| pup birth mass | -0.01 | -0.31 – 0.29 | -0.05 | 0.963 | 0.27 | 0.10 – 0.45 | 3.04 | **0.002** |
| pup age | 0.05 | 0.04 – 0.07 | 5.71 | **<0.001** | 0.06 | 0.05 – 0.07 | 10.56 | **<0.001** |
| season [2019] | 1.67 | 1.21 – 2.12 | 7.25 | **<0.001** | 1.75 | 1.46 – 2.04 | 11.92 | **<0.001** |
| season [2020] | 2.02 | 1.62 – 2.43 | 9.82 | **<0.001** | 1.62 | 1.33 – 1.91 | 11.00 | **<0.001** |
| season [2021] | 1.51 | 1.09 – 1.93 | 7.14 | **<0.001** | 1.39 | 1.09 – 1.69 | 9.16 | **<0.001** |
| mother sMLH | -0.66 | -2.32 – 0.99 | -0.79 | 0.432 | | | | |
| mother age | 0.03 | -0.04 – 0.09 | 0.84 | 0.401 | | | | |
| Observations | 291 | | | | 721 | | | |
| $R^2$ / $R^2$ adjusted | 0.450 / 0.433 | | | | 0.388 / 0.382 | | | |

### Statistical models - SNP inbreeding

#### Birth weight analysis

In this section of the script factors affecting birth weight are investigated.

##### Filtered data for birth weight analysis

Mums that were no genetic match with their pups were removed (concerns pup H2 and H5, they were switched and suckled by the others mum).

##### Data visualization

Before fitting the models, the raw data is visualized to explore distribution.

```
#min/mean/max of birth weight
mean_birth_weight <- mean(Unique_Day60$Birth_weight, na.rm = T)
#5.61
#min_birth_weight <- min(Unique_Day60$Birth_weight, na.rm = T)
#3.2
#max_birth_weight <- max(Unique_Day60$Birth_weight, na.rm = T)
#7.8

birth_weight_raw <- ggplot(Unique_Day60, aes(x = Birth_weight)) +
  geom_histogram(binwidth = 0.2, fill = "orange", color = "black") +
  geom_vline(aes(xintercept = mean_birth_weight), color = "purple", linetype = "dashed",
  ↪  linewidth = 0.8) +
  labs(title = "Birth weight distribution",
       x = "Weight (kg)",
       y = "Frequency") +
  theme_minimal()
#Based on visual inspection, the birth weight looks fairly normally distributed

shapiro.test(Unique_Day60$Birth_weight) #the shapiro-wilk test tests for normality. The
↪  null-hypothesis is that the population is normally distributed. P value > 0.05
↪  implying that the distribution of the data are not significantly different from
↪  normal distribution. Therefore, we can assume normality.
```

```
##
##  Shapiro-Wilk normality test
##
## data:  Unique_Day60$Birth_weight
## W = 0.99537, p-value = 0.9855
```

```
#plots
plot_grid(birth_weight_raw, label_size = 12)
```

```
## Warning: Removed 1 rows containing non-finite values (`stat_bin()`).
```

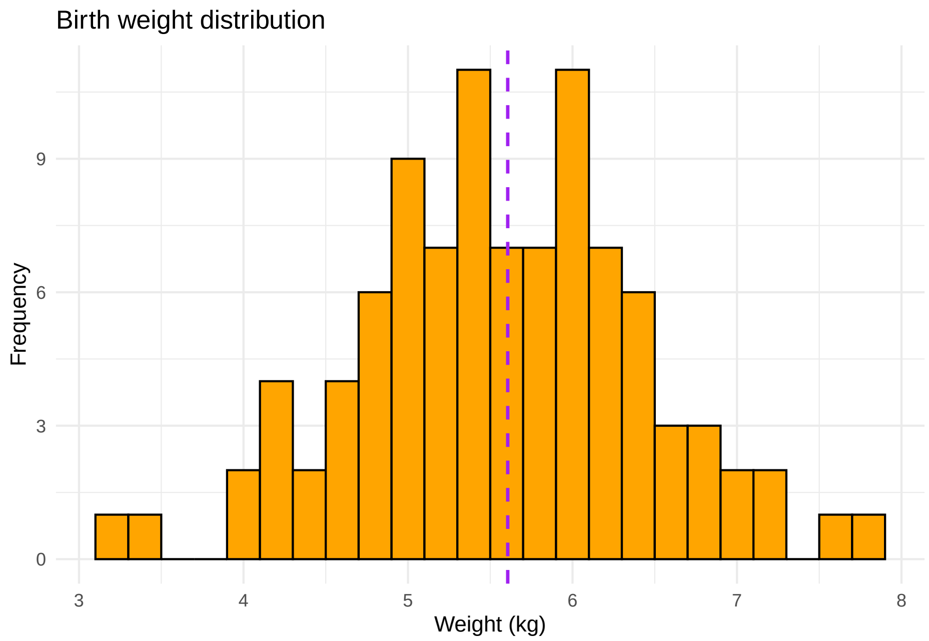

##### Model

The birth weight data is normally distributed, so the model is built with a Gaussian distribution.

```
# Model with Froh
BW.model.froh <- lm(Birth_weight ~ froh +
                      froh_mum +
                      Sex +
                      Season +
                      Beach,
                      data = Unique_Day60)

#plot(BW.model.froh)
summary(BW.model.froh)
```

```
##
## Call:
## lm(formula = Birth_weight ~ froh + froh_mum + Sex + Season +
##     Beach, data = Unique_Day60)
##
## Residuals:
##      Min       1Q   Median       3Q      Max
## -2.70382 -0.51808 -0.02644  0.46211  2.34856
##
## Coefficients:
##             Estimate Std. Error t value Pr(>|t|)
## (Intercept)   4.6903     0.8462   5.543 3.28e-07 ***
## froh          3.3658     8.1860   0.411   0.6820
## froh_mum      3.7677     7.7100   0.489   0.6263
## SexM          0.4369     0.1859   2.350   0.0211 *
## Season1920    0.2804     0.1766   1.588   0.1161
## BeachSSB      0.1172     0.1772   0.661   0.5103
## ---
```

```
## Signif. codes:  0 '***' 0.001 '**' 0.01 '*' 0.05 '.' 0.1 ' ' 1
##
## Residual standard error: 0.836 on 85 degrees of freedom
##   (7 observations deleted due to missingness)
## Multiple R-squared:  0.09185,    Adjusted R-squared:  0.03843
## F-statistic: 1.719 on 5 and 85 DF,  p-value: 0.1389
```

##### Residual check of model

```r
#Test model
testDispersion(BW.model.froh) #good fit
```

```
##
##  DHARMa nonparametric dispersion test via sd of residuals fitted vs.
##  simulated
##
## data:  simulationOutput
## dispersion = 0.94117, p-value = 0.656
## alternative hypothesis: two.sided
```

```r
plotQQunif(BW.model.froh) #deviation not significant
plotResiduals(BW.model.froh)

#Test model fit
#Predict values for your existing data
predicted_values <- predict(BW.model.froh, type = "response")
na_values <- rep(NA, 7) #to make the predicted no. of values equal to the observed data
predicted_values <- c(predicted_values, na_values)

#plot the densities of the predicted and observed data
BW_model <- ggplot(Unique_Day60, aes(x = Birth_weight)) +
  geom_density(fill = "orange", alpha = 0.5) +
  geom_density(aes(x = predicted_values), fill = "purple", alpha = 0.5) +
  labs(x = "Birth weight)", y = "Density") +
  ggtitle("Observed vs. Predicted birth weight")

BW_model #Note: densities does not match greatly
```

```
## Warning: Removed 1 rows containing non-finite values (`stat_density()`).

## Warning: Removed 7 rows containing non-finite values (`stat_density()`).
```

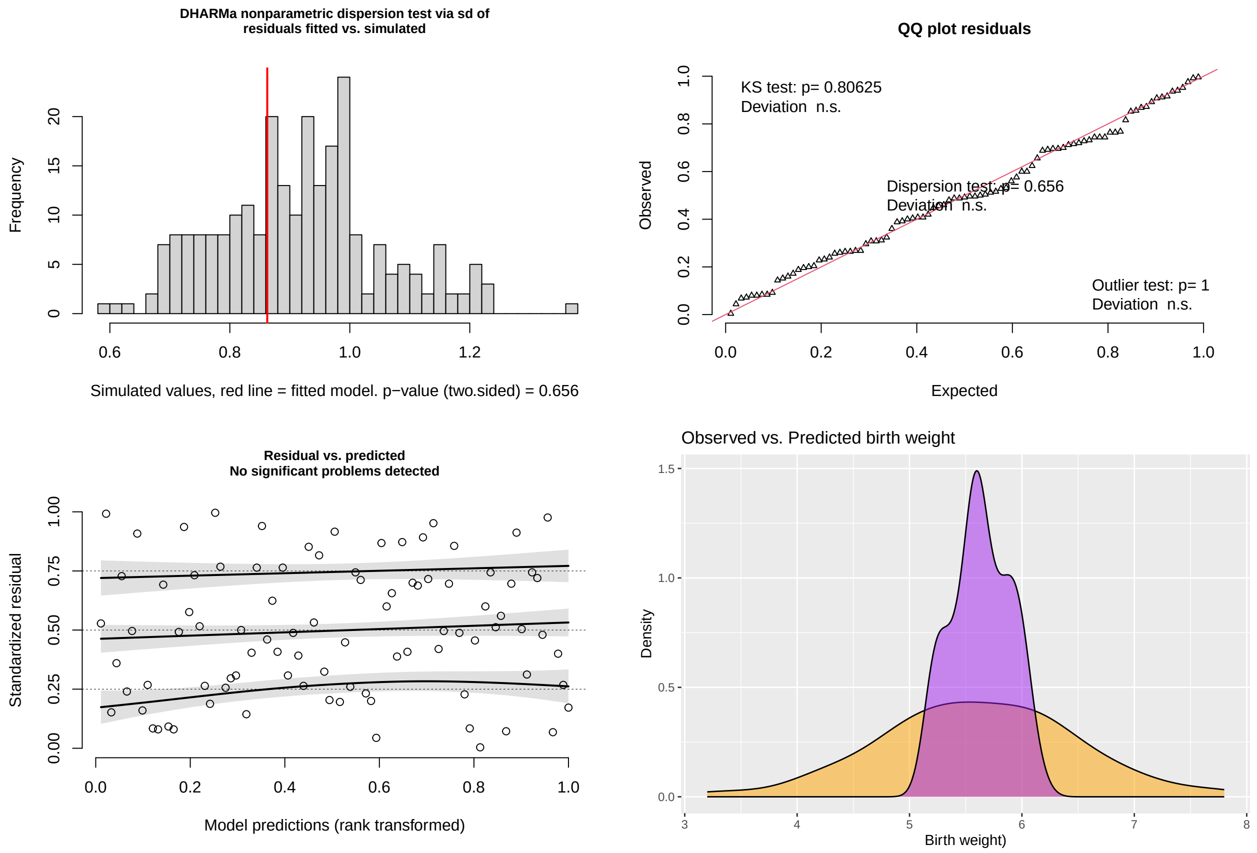

#### Paramter estimates pup birth weight model

```
# Table
lab_BW.model.froh <- c(
  `(Intercept)` = "Intercept",
  froh = "pup <i>F</i><sub>ROH</sub>", # for some reason, if I do not add the htlm
   ↪  space ( ), it puts Froh on the line below "pup". This fixes that
  froh_mum = "mother <i>F</i><sub>ROH</sub>",
  SexM = "pup sex [M]",
  BeachSSB = "colony [SSB]",
  Season1920 = "season [2020]")

print(sjPlot::tab_model(BW.model.froh,
                        title = "Pup birth mass",
                        dv.labels = "model incl. maternal effect",
                        pred.labels = lab_BW.model.froh,
                        show.stat=T,
                        string.stat = "t value",
                        file = here("Tables", "Table_BW_SNPs_Froh.html")))
webshot::webshot(here("Tables", "Table_BW_SNPs_Froh.html"),
                 file=here("Tables", "Table_BW_SNPs_Froh.png"), delay=2, vheight = 350,
                  ↪  vwidth = 450)
```

**Pup birth mass**

| | model incl. maternal effect | | | |
|---|---|---|---|---|
| *Predictors* | *Estimates* | *CI* | *t value* | *p* |
| Intercept | 4.69 | 3.01 – 6.37 | 5.54 | **<0.001** |
| pup $F_{ROH}$ | 3.37 | -12.91 – 19.64 | 0.41 | 0.682 |
| mother $F_{ROH}$ | 3.77 | -11.56 – 19.10 | 0.49 | 0.626 |
| pup sex [M] | 0.44 | 0.07 – 0.81 | 2.35 | **0.021** |
| season [2020] | 0.28 | -0.07 – 0.63 | 1.59 | 0.116 |
| colony [SSB] | 0.12 | -0.24 – 0.47 | 0.66 | 0.510 |
| Observations | 91 | | | |
| $R^2$ / $R^2$ adjusted | 0.092 / 0.038 | | | |

#### Survival analysis

First step for the survival analysis is to transform the 'Death' variable into a binary (0,1) variable and remove the two pups that were not cared for by their biological mum.

*Note: for the pups C20 and N1 their mums, F20 and FWB1, respectively, died during the sampling season as well.*

##### Binomial data for survival analysis

```
Unique_Day60$Survived <- ifelse(Unique_Day60$Death == 'N',1,0)
```

##### Model

The model is built with a binary response variable. The model includes includes $F_{ROH}$ values calculated from SNP data.

```
# Survival model with Froh
Survival.model.froh <- glm(Survived ~ froh +
                               froh_mum +
                               Sex +
                               Season +
                               Beach +
                               Birth_weight,
                               data = Unique_Day60, family = 'binomial')

summary(Survival.model.froh)
```

```
##
## Call:
## glm(formula = Survived ~ froh + froh_mum + Sex + Season + Beach +
```

```
##     Birth_weight, family = "binomial", data = Unique_Day60)
##
## Deviance Residuals:
##     Min       1Q   Median       3Q      Max
## -2.2299   0.2761   0.4397   0.6643   1.3074
##
## Coefficients:
##              Estimate Std. Error z value Pr(>|z|)
## (Intercept)   -4.2193     3.4102  -1.237   0.2160
## froh          28.9553    26.6320   1.087   0.2769
## froh_mum     -19.3476    26.6919  -0.725   0.4685
## SexM           0.1097     0.6558   0.167   0.8671
## Season1920    -0.1318     0.5818  -0.227   0.8208
## BeachSSB       1.3624     0.6211   2.193   0.0283 *
## Birth_weight   0.7982     0.3916   2.038   0.0415 *
## ---
## Signif. codes:  0 '***' 0.001 '**' 0.01 '*' 0.05 '.' 0.1 ' ' 1
##
## (Dispersion parameter for binomial family taken to be 1)
##
##     Null deviance: 93.248  on 90  degrees of freedom
## Residual deviance: 79.310  on 84  degrees of freedom
##   (7 observations deleted due to missingness)
## AIC: 93.31
##
## Number of Fisher Scoring iterations: 5
```

```
#Binary variable codes so 0 means the pup died that season and 1 means the pup survived.
#Birth weight and beach is significant
```

##### Residual check of model

```
#Test model
testDispersion(Survival.model.froh) #good fit
```

```
##
##  DHARMa nonparametric dispersion test via sd of residuals fitted vs.
##  simulated
##
## data:  simulationOutput
## dispersion = 1.0344, p-value = 0.864
## alternative hypothesis: two.sided
```

```
plotQQunif(Survival.model.froh) #deviation not significant
plotResiduals(Survival.model.froh)
# The residuals vs. predictred quantile plot shows a small deviation for the 0.75
↪ quantile. However, the combined adjusted quantile test is none significant and also
↪ the Kolmogorov-Smirnov(KS)-test is non significant (see QQplot). So we conclude that
↪ the deviation is not very big, and no reason to reject the model.
```

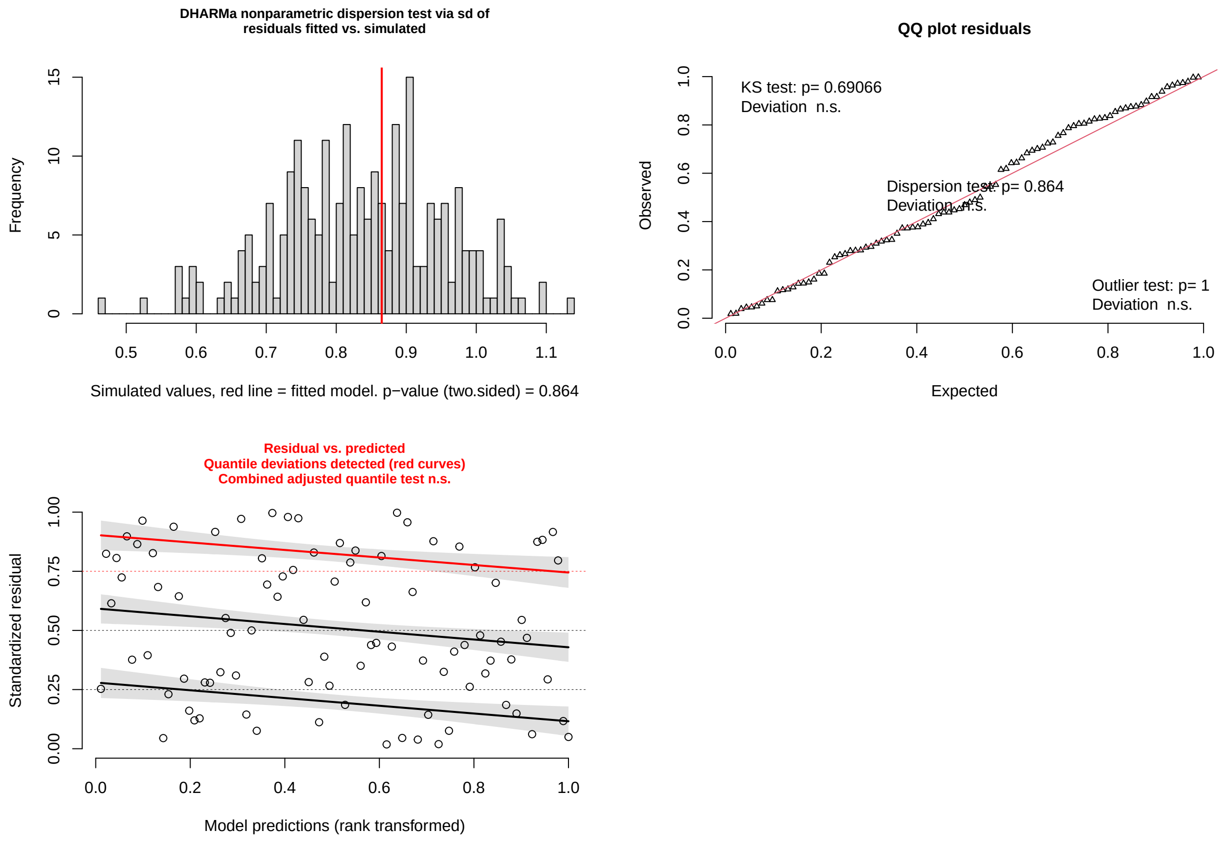

##### Paramter estimates pup survival model

```
lab_Survival.model.froh <- c(
  `(Intercept)` = "Intercept",
  froh = "pup <i>F</i><sub>ROH</sub>",
  froh_mum = "mother <i>F</i><sub>ROH</sub>",
  SexM = "pup sex [M]",
  Season1920 = "season [2020]",
  BeachSSB = "colony [SSB]",
  Birth_weight = "pup birth mass")

print(sjPlot::tab_model(Survival.model.froh,
                        title = "Pup survival",
                        dv.labels = "model incl. maternal effect",
                        pred.labels = lab_Survival.model.froh,
                        transform = NULL,
                        show.stat=T,
                        string.stat = "t value",
                        file = here("Tables", "Table_survival_SNPs_Froh.html")))
webshot::webshot(here("Tables", "Table_survival_SNPs_Froh.html"),
                 file=here("Tables", "Table_survival_SNPs_Froh.png"), delay=2, vheight =
                 ↪ 400, vwidth = 400)
```

**Pup survival**

| | model incl. maternal effect | | | |
|---|---|---|---|---|
| *Predictors* | *Log-Odds* | *CI* | *t value* | *p* |
| Intercept | -4.22 | -11.31 – 2.24 | -1.24 | 0.216 |
| pup $F_{\text{ROH}}$ | 28.96 | -21.78 – 83.83 | 1.09 | 0.277 |
| mother $F_{\text{ROH}}$ | -19.35 | -73.54 – 32.49 | -0.72 | 0.469 |
| pup sex [M] | 0.11 | -1.21 – 1.40 | 0.17 | 0.867 |
| season [2020] | -0.13 | -1.30 – 1.01 | -0.23 | 0.821 |
| colony [SSB] | 1.36 | 0.20 – 2.68 | 2.19 | **0.028** |
| pup birth mass | 0.80 | 0.09 – 1.65 | 2.04 | **0.042** |
| Observations | 91 | | | |
| $R^2$ Tjur | 0.153 | | | |

#### Growth curves with repeated measures

The dataset from FWB and SSB in season 1819/1920 contains repeated measures of growth at 6 different time points until tagging. This section contains the analysis of growth based on growth curves utilizing the repeated measures. *RM = repeated measures*

##### Data visualization

Before fitting the models, the raw data is visualized to explore distribution. Weight data is visually expected on its own and fitted against age in days to understand general distribution and pattern over time. The raw data is visually expected using ggplot2 and further explored using the Shapiro-Wilk test.

```
#mean of weight_kg
mean_weight <- mean(SurvivorsRM_Day60$Weight_kg, na.rm = T)
#7.684354
#max_tagging_weight <- max(SurvivorsRM_Day60$Last_weight, na.rm = T)
#16.3

weight_raw <- ggplot(SurvivorsRM_Day60, aes(x = Weight_kg)) +
  geom_histogram(binwidth = 0.4, fill = "orange", color = "black") +
  geom_vline(aes(xintercept = mean_weight), color = "purple", linetype = "dashed",
   ↪ linewidth = 0.8) +
  labs(title = "Weight distribution",
       x = "Weight (kg)",
       y = "Frequency") +
  theme_minimal()
#Based on visual inspection, data is slightly right-skewed
```

```
ggplot(SurvivorsRM_Day60, aes(x = log(Weight_kg))) +
  geom_histogram(binwidth = 0.1, fill = "orange", color = "black") +
  labs(title = "Weight distribution (log weight)",
       x = "log(Weight (kg))",
       y = "Frequency") +
  theme_minimal()
```

```
## Warning: Removed 2 rows containing non-finite values (`stat_bin()`).
```

```
shapiro.test(SurvivorsRM_Day60$Weight_kg) #the shapiro-wilk test tests for normality. The
↪ null-hypothesis is that the population is normally distributed. The test is
↪ significant, so we reject the null-hypothesis. The weight_kg data is not normally
↪ distributed. By scaling the outcome variable, we can make it more suitable to built
↪ models with, even though it does not necessarily normalize the distribution.
```

```
##
##  Shapiro-Wilk normality test
##
## data:  SurvivorsRM_Day60$Weight_kg
## W = 0.96611, p-value = 8.89e-09
```

```
#shapiro.test(scale(SurvivorsRM_Day60$Weight_kg)) still not normally distributed, however
↪ if it allows for normally distributed residuals down stream, it is valid.

#SurvivorsRM_Day60$Weight_kg_scale <- scale(SurvivorsRM_Day60$Weight_kg)
SurvivorsRM_Day60$Age_Days_scale <- scale(SurvivorsRM_Day60$Age_Days)
```

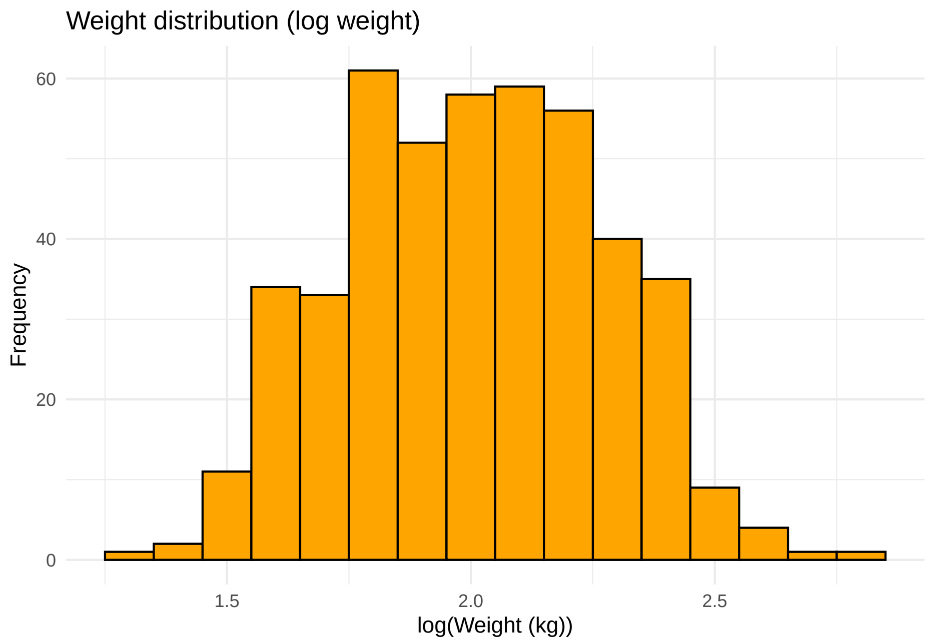

```
weight_over_time <- ggplot(SurvivorsRM_Day60, aes(x = Age_Days, y = Weight_kg)) +
  geom_point() +
  labs(title = "Weight gain over time",
       x = "Days",
       y = "Weight (kg)") +
  theme_minimal()
```

```r
#plots
plot_grid(weight_raw, weight_over_time, labels = "AUTO", label_size = 12)
```

```
## Warning: Removed 2 rows containing non-finite values (`stat_bin()`).

## Warning: Removed 2 rows containing missing values (`geom_point()`).
```

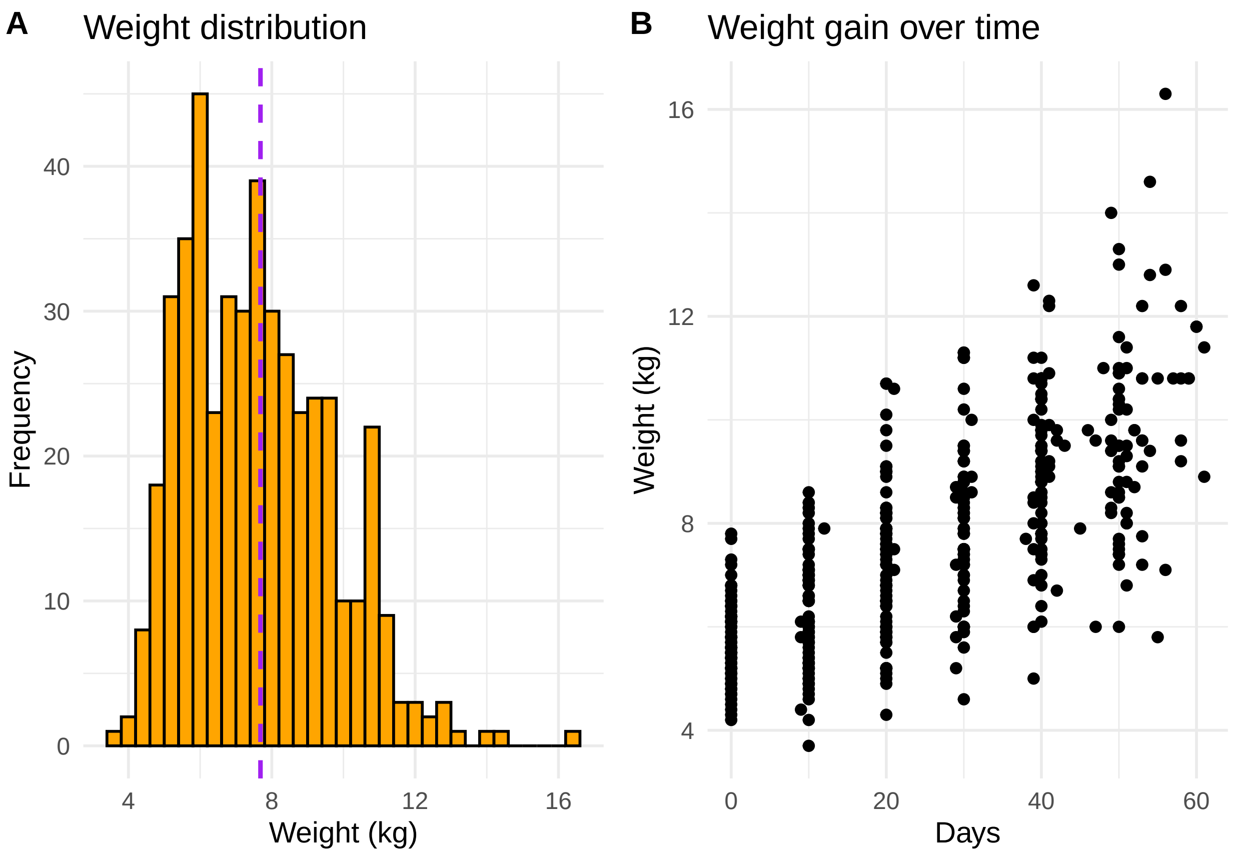

Based on both visual inspection and the shapiro-wilk test, the weight data does not follow a normal distribution but is slightly right-skewed.

##### Exploration of growth curve fit

We can explore different growth curves. In the larger dataset birth weight ranges from 2.45 to 7.7 kg and weight at day 60 from 6.5 to 14.6 kg. In this dataset with repeated measures, lowest birth weight is 4.2 and highest last weight is 16.3.

```r
#For the logistic and gompertz models, the parameters K, r and t are used to fit the
↪ model:
#K is the max weight the pups can reach, set to 20kg, as heaviest pup was 16.3
#r is the growth rate, the average growth rate is used: 0.080821
#t is the inflection point; the time at which the pups growth most rapidly.
#set at 30days, as this is the mid-point

#logistic growth model
logistic.model <- nls(Weight_kg ~ K / (1 + exp(-r * (Age_Days - t))),
                      data = SurvivorsRM_Day60,
                      start = list(K = 20, r = 0.080821, t = 30))
```

```r
#gompertz growth model
gompertz.model <- nls(Weight_kg ~ K * exp(-exp(-r * (Age_Days - t))),
                      data = SurvivorsRM_Day60,
                      start = list(K = 20, r = 0.080821, t = 30))

#For the linear model, a and b are parameters used to fit the model:
#a is the linear growth rate; the average growth rate is used: 0.080821
#b is the birth weight, here set at 2.45kg, as lightest measured pup was 2.45kg.

#Linear
linear.model <- nls(Weight_kg ~ a * Age_Days + b,
                    data = SurvivorsRM_Day60,
                    start = list(a = 0.080821, b = 2.45))

# Compare the models using AIC
models.1 <- list(linear.model, logistic.model, gompertz.model)
mod.names.1 <- c('linear.model', 'logistic.model', 'gompertz.model')
AIC_growth_curves <- aictab(cand.set = models.1, modnames = mod.names.1)

AIC_growth_curves
```

```
##
## Model selection based on AICc:
##
##                K    AICc Delta_AICc AICcWt Cum.Wt      LL
## linear.model   3 1623.24       0.00   0.52   0.52 -808.59
## logistic.model 4 1624.81       1.57   0.24   0.76 -808.36
## gompertz.model 4 1624.82       1.58   0.24   1.00 -808.36
```

```r
#in this basic model format, the linear model is the best fit

# Visualize the fits
ggplot(SurvivorsRM_Day60, aes(x = Age_Days, y = Weight_kg)) +
  geom_point() +
  geom_smooth(method = "nls", formula = y ~ a * x + b, se = FALSE, color = "orange",
              method.args = list(start = coef(linear.model))) +
  geom_smooth(method = "nls", formula = y ~ K / (1 + exp(-r * (x - t))), se = FALSE,
   ↪ color = "red",
              method.args = list(start = coef(logistic.model))) +
  geom_smooth(method = "nls", formula = y ~ K * exp(-exp(-r * (x - t))), se = FALSE,
   ↪ color = "purple",
              method.args = list(start = coef(gompertz.model))) +
  labs(title = "Growth Curve Models Comparison",
       x = "Days",
       y = "Weight") +
  theme_minimal()
```

```
## Warning: Removed 2 rows containing non-finite values (`stat_smooth()`).
## Removed 2 rows containing non-finite values (`stat_smooth()`).
## Removed 2 rows containing non-finite values (`stat_smooth()`).

## Warning: Removed 2 rows containing missing values (`geom_point()`).
```

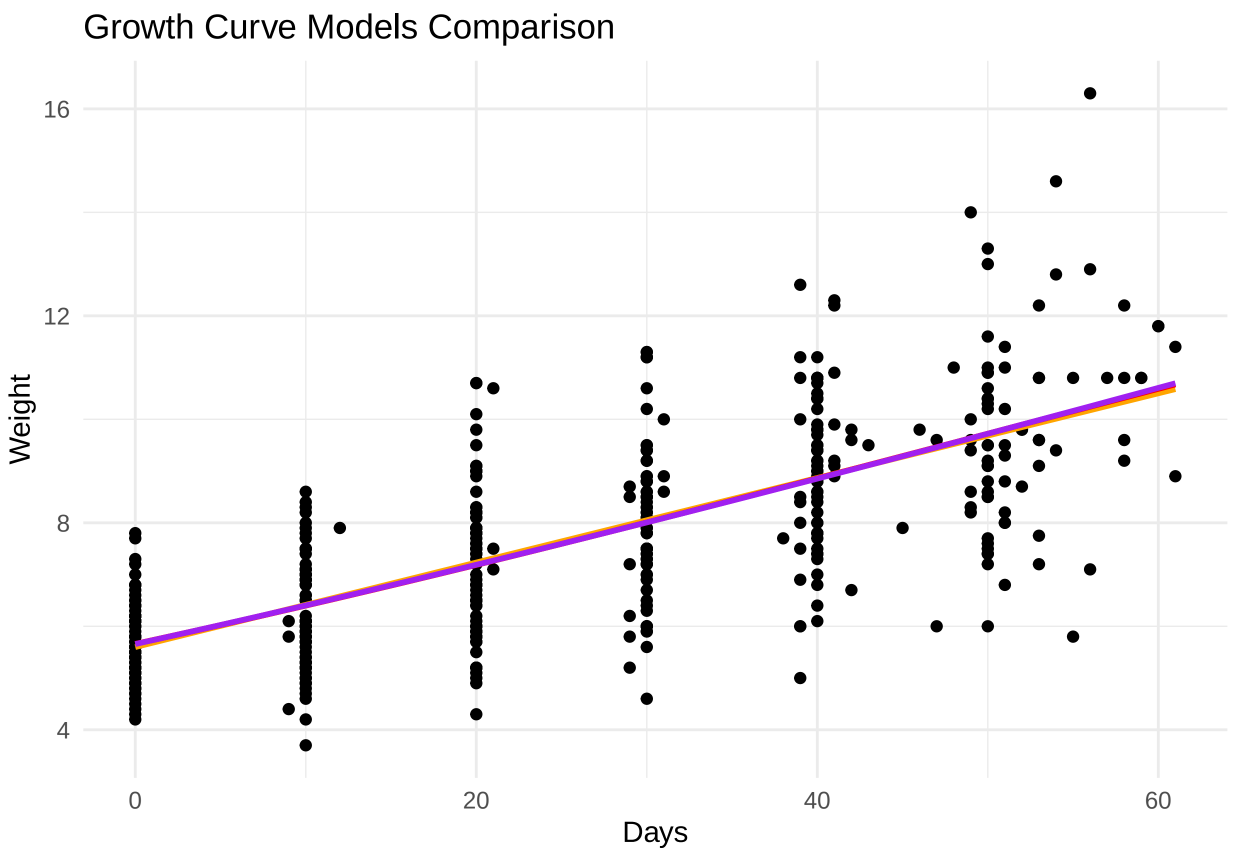

```
#clean up
rm(linear.model, logistic.model, gompertz.model, models.1, mod.names.1)
```

The different growth curves all lie within a delta AIC of 2, which means the general difference between the fit of the different growth curves to the data is not strong. Therefore, we proceed with the linear model structure.

##### Model: weight gain

As the growth curves turn out to be linear, we made a model of weight gain based on last weight - first weight.

```
##
##  Shapiro-Wilk normality test
##
## data:  UniqueSurvivors_Day60$Total_weight_gain
## W = 0.9756, p-value = 0.1561
```

```
##
## Call:
## lm(formula = Total_weight_gain ~ froh + froh_mum + Sex + Season +
##     Beach + Birth_weight + Last_day, data = UniqueSurvivors_Day60)
##
## Residuals:
##      Min       1Q   Median       3Q      Max
## -3.00439 -0.79453 -0.02775  0.89505  2.98155
##
```

```
## Coefficients:
##               Estimate Std. Error t value Pr(>|t|)
## (Intercept)   -2.12427    2.84783  -0.746  0.45844
## froh         -23.94247   17.24380  -1.388  0.16981
## froh_mum      46.61559   14.49353   3.216  0.00204 **
## SexM           2.05529    0.37199   5.525 6.45e-07 ***
## Season1920    -0.59785    0.34349  -1.741  0.08657 .
## BeachSSB       0.13263    0.33511   0.396  0.69358
## Birth_weight   0.05517    0.22675   0.243  0.80854
## Last_day       0.06352    0.03559   1.784  0.07910 .
## ---
## Signif. codes:  0 '***' 0.001 '**' 0.01 '*' 0.05 '.' 0.1 ' ' 1
##
## Residual standard error: 1.375 on 64 degrees of freedom
##   (4 observations deleted due to missingness)
## Multiple R-squared:  0.4371, Adjusted R-squared:  0.3755
## F-statistic: 7.098 on 7 and 64 DF,  p-value: 2.87e-06
```

##### Residual check of model

```
testDispersion(Growth_model) #good fit
```

```
##
##  DHARMa nonparametric dispersion test via sd of residuals fitted vs.
##  simulated
##
## data:  simulationOutput
## dispersion = 0.89925, p-value = 0.568
## alternative hypothesis: two.sided
```

```
plotQQunif(Growth_model) #deviation not significant
plotResiduals(Growth_model)

# Test model fit
# Predict values for your existing data
predicted_values <- predict(Growth_model, type = "response")
na_values <- rep(NA, 4) #to make the predicted no. of values equal to the observed data
predicted_values <- c(predicted_values, na_values)

#plot the densities of the predicted and observed data
growth_model <- ggplot(UniqueSurvivors_Day60, aes(x = Total_weight_gain)) +
  geom_density(fill = "orange", alpha = 0.5) +
  geom_density(aes(x = predicted_values), fill = "purple", alpha = 0.5) +
  labs(x = "Weight gain", y = "Density") +
  ggtitle("Observed vs. Predicted Weight gain")

plot_grid(growth_model, label_size = 12)
```

```
## Warning: Removed 1 rows containing non-finite values (`stat_density()`).
```

```
## Warning: Removed 4 rows containing non-finite values (`stat_density()`).
```

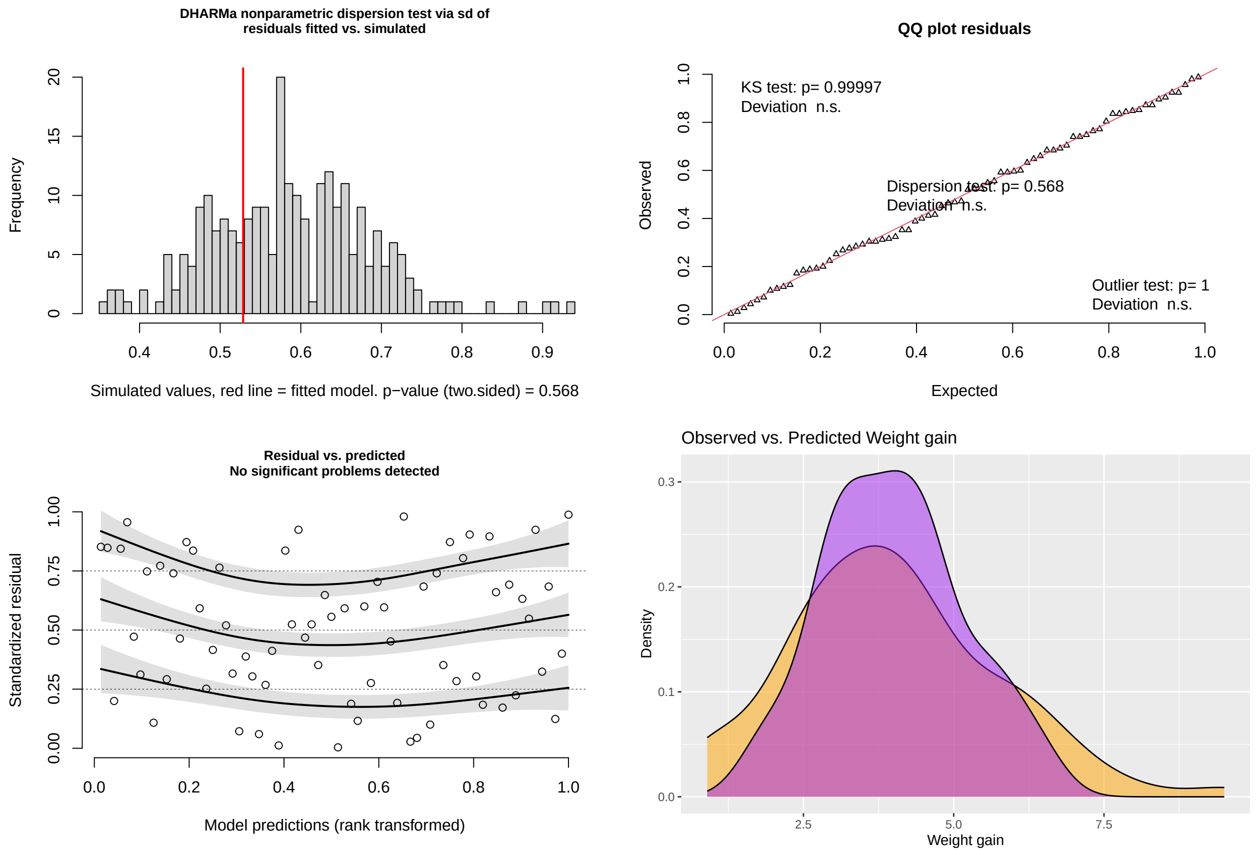

##### Parameter estimates pup growth model

```
lab_weight_gain_froh <- c(
  `(Intercept)` = "Intercept",
  froh = "pup <i>F</i><sub>ROH</sub>",
  froh_mum = "mother <i>F</i><sub>ROH</sub>",
  SexM = "pup sex [M]",
  Season1920 = "season [2020]",
  BeachSSB = "colony [SSB]",
  Birth_weight = "pup birth mass",
  Last_day = "pup age")

print(sjPlot::tab_model(Growth_model,
                        title = "Pup growth",
                        dv.labels = "model incl. maternal effect",
                        pred.labels = lab_weight_gain_froh,
                        show.stat=T,
                        string.stat = "t value",
                        file = here("Tables", "Table_Weight_gain_Froh.html")))
webshot::webshot(here("Tables", "Table_Weight_gain_Froh.html"),
                 file=here("Tables", "Table_Weight_gain_Froh.png"), delay=2, vheight =
                 ↪  450, vwidth = 500)
```

**Pup growth**

| | model incl. maternal effect | | | |
|---|---|---|---|---|
| *Predictors* | *Estimates* | *CI* | *t value* | *p* |
| Intercept | -2.12 | -7.81 – 3.56 | -0.75 | 0.458 |
| pup $F_{ROH}$ | -23.94 | -58.39 – 10.51 | -1.39 | 0.170 |
| mother $F_{ROH}$ | 46.62 | 17.66 – 75.57 | 3.22 | **0.002** |
| pup sex [M] | 2.06 | 1.31 – 2.80 | 5.53 | **<0.001** |
| season [2020] | -0.60 | -1.28 – 0.09 | -1.74 | 0.087 |
| colony [SSB] | 0.13 | -0.54 – 0.80 | 0.40 | 0.694 |
| pup birth mass | 0.06 | -0.40 – 0.51 | 0.24 | 0.809 |
| pup age | 0.06 | -0.01 – 0.13 | 1.78 | 0.079 |
| Observations | 72 | | | |
| $R^2$ / $R^2$ adjusted | 0.437 / 0.375 | | | |

##### Individual growth curves

For the models with repeated measures, a random effects term 'ID' is added to inform the model that some observations are clustered within the same individual. The base is built around a generalized linear mixed-effects model (GLMM) with Gamma(link = "identity") data distribution to account for the repeated measures and the right-skewed data.

```
# Unconditional means model aka null model (no predictors)
RM0 <- glmer(Weight_kg ~ 1 + (1 | ID),
             data = SurvivorsRM_Day60,
             family = Gamma(link="identity"))
#summary(RM0)

# Growth varying per day, each individual has different intercepts, but the same slope
RM <- glmer(Weight_kg ~ Age_Days + (1 | ID),
            data = SurvivorsRM_Day60,
            family = Gamma(link="identity"))
#summary(RM)

# Growth varying per day, each individual has different slopes ((Age_Days | ID))
#the -1 in (Age_Days_scale - 1 | ID) suppress the intercept
RM1 <- glmer(Weight_kg ~ Age_Days + (Age_Days - 1 | ID),
             data = SurvivorsRM_Day60,
             family = Gamma(link="identity"))
#summary(RM1)
```

```r
# Growth varying per day, both intercept and slope vary per individual
RM2 <- glmer(Weight_kg ~ Age_Days + (1 + Age_Days | ID),
             data = SurvivorsRM_Day60,
             family = Gamma(link="identity"))
#summary(RM2)

# Explanation: Fixed effects estimate: Intercept is average birth weight
                                    # Age_Days is one day change in weight
            # Random effects: associated variance
            # Intercept: how much variance in birth weight between pups
            # Age_Days: difference in slope. 0.00060 might not sound as much
            # Corr: the correlation between the slope and intercept: 0.36
            # it's positive, which indicates that pups with higher intercept
            # on average has a steeper slope as well.
            # Residual: 0.5593029. Refers to the variance not explained by the variables
            ↪  in the model
            # additionally, looking at confint(RM2), the confidence interval
            # for the intercept and for the Age_Days does not cross 0.
#ICC = repeatability of weight across individuals

models <- list(RM0, RM, RM1, RM2)
mod.names <- c('null-model', 'intercept', 'slope', 'i+s')
AIC_model_structure <- aictab(cand.set = models, modnames = mod.names)
AIC_model_structure
```

```
##
## Model selection based on AICc:
##
##            K    AICc Delta_AICc AICcWt Cum.Wt      LL
## i+s        6 1112.42       0.00      1      1 -550.12
## intercept  4 1247.39     134.97      0      1 -619.65
## slope      4 1286.04     173.61      0      1 -638.98
## null-model 3 1819.12     706.70      0      1 -906.54
```

```r
# The RM2 model, which allows both intercept and slope to vary per individual performs
↪  best.
#testDispersion(RM2)
#plotQQunif(RM2)

# Clean up
rm(RM0, RM1, RM, models, mod.names)
```

### Manuscript figures

#### Figure 1: map and seasonal data

```
#~~~~~~~~~~~~~~~~#
#  Load data  ####
#~~~~~~~~~~~~~~~~#

seasonal_data <- read.table(here("Data", "Raw", "seasonal_data.txt"), sep = "\t",
↪  stringsAsFactors = F, header = T)

#~~~~~~~~~~~~~~~~~~~~#
#  Seasonal data  ####
#~~~~~~~~~~~~~~~~~~~~#

source(here("Rcode", "anneke_theme.R"))

#~~ Make a list for the theme so it is the same for all figures
gglayer_theme <- list(
  scale_x_discrete(labels = c(`2017-2018` = "2018", `2018-2019` = "2019", `2019-2020` =
   ↪  "2020", `2020-2021` = "2021")),
  theme_anneke(),
  theme(axis.line.x = element_line(colour = 'black', linetype='solid'),
        axis.line.y = element_line(colour = 'black', linetype='solid'),
        plot.title = element_text(size = rel(1)))
)

#~~ Make sub plots
# Plot a is blank canvas + title where later the map gets added
p_a <- ggplot(seasonal_data %>% filter(variable=="SSB ESTIMATED NUMBER OF FEMALE
↪  BREEDERS"),
              aes(x = season, y = mean)) +
  #geom_pointrange(aes(ymin = CI95_low, ymax = CI95_high)) +
  #geom_point(shape = 22, size = 4, fill = "#eb7f86") +
  labs(title="(a) Map of Bird Island", x= "", y="") +
  gglayer_theme +
  theme(axis.line.x = element_blank(),
        axis.line.y = element_line(colour = 'white', linetype='solid'),
        axis.text = element_text(colour = "white"),
        panel.grid=element_blank(),
        panel.grid.major=element_blank(),
        panel.grid.minor=element_blank())

# Breeding females
p_breeders <-  ggplot(seasonal_data%>% filter(variable=="SSB ESTIMATED NUMBER OF FEMALE
↪  BREEDERS"),
                      aes(x = season, y = mean)) +
  geom_pointrange(aes(ymin = CI95_low, ymax = CI95_high)) +
  geom_point(shape = 22, size = 4, fill = "dimgrey") +
  labs(title="(b) Female breeders", x= "Year", y="No. of breeders") +
  gglayer_theme
```

```r
# Female pup birth mass
p_bm <-  ggplot(seasonal_data%>% filter(variable=="SSB FEMALE PUP BIRTH MASS (kg)"),
               aes(x = season, y = mean)) +
  geom_pointrange(aes(ymin = CI95_low, ymax = CI95_high)) +
  geom_point(shape = 22, size = 4, fill = "dimgrey") +
  labs(title="(c) Female pup birth mass", x= "Year", y="Birth mass (kg)") +
  gglayer_theme

# Female foraging trip duration
p_foraging <-  ggplot(seasonal_data %>% filter(variable=="FWB FEMALE FORAGING TRIP
 ↪  DURATION (days)"),
                     aes(x = season, y = mean)) +
  geom_pointrange(aes(ymin = CI95_low, ymax = CI95_high)) +
  geom_point(shape = 22, size = 4, fill = "dimgrey") +
  labs(title="(d) Female foraging trip duration", x="Year", y="Time at sea (days)") +
  gglayer_theme

#~~~~~~~~~~~~~~~~~~~~~~~#
#  Bird Island map  ####
#~~~~~~~~~~~~~~~~~~~~~~~#

#~~ Bird island maps
bi_coast <- st_read(here("Rcode", "Bird Island Map", "Map_Old",
 ↪  "BI_Coast_Projected_new.shp"), quiet = TRUE)
# bi_r <- st_read(here("Rcode", "Bird Island Map", "rivers_lines", "sg_bird_rivers.shp"),
 ↪  quiet = TRUE) #rivers
# bi_c <- st_read(here("Rcode", "Bird Island Map", "contours", "sg_bird_contours.shp"),
 ↪  quiet = TRUE) # contours
# bi <- st_read(here("Rcode", "Bird Island Map", "coastline", "sg_bird_coast.shp"), quiet
 ↪  = TRUE) #surface

#~~ Outline of SSB and FWB, made in Google Earth
ssb <- st_read(here("Rcode", "Bird Island Map", "beachs", "SSB.kml", "doc.kml"), quiet =
 ↪  TRUE)
fwb <- st_read(here("Rcode", "Bird Island Map", "beachs", "FWB.kml", "doc.kml"), quiet =
 ↪  TRUE)

#### mapping bird island ----
plot.bi.color <- ggplot() +
  geom_sf(data = bi_coast, fill = "dimgrey") +  #"#ADADAD"
  #geom_sf(data = bi_r, color = "blue") +
  #geom_sf(data = bi_c) +
  geom_sf(data=ssb, fill = "#872ca2") + #ssb
  geom_sf(data=fwb, fill = "#fa7876") + #fwb
  theme(legend.position="none") +
  theme_void()

#### mapping study colonies ----
# Adds box around study colonies on bird island map
plot.bi.color. <- plot.bi.color +
```

```
  annotate(geom = "rect",
           xmin = -38.060,
           xmax = -38.045,
           ymin = -54.014,
           ymax = -54.0065,
           fill = NA, # transparent bg
           color = "black" )

# Adds beach location in color
plot.bi.beaches.color <- ggplot() +
  geom_sf(data = bi_coast, fill = "NA") +
  geom_sf(data = bi_coast, fill = "#ADADAD") +  # "#eaeaea"
  geom_sf(data = fwb, fill = "#fa7876") +
  geom_sf(data = ssb, fill = "#872ca2") +
  theme(legend.position = "none") +
  theme_void()

# Add text
plot.bi.beaches.color <- plot.bi.beaches.color +
  coord_sf(xlim = c(-38.060, -38.045),
           ylim = c(-54.014, -54.0065),
           expand = FALSE) +
  annotation_scale(aes(location="br", style = "ticks")) +
  theme(panel.border = element_rect(colour = "black", fill=NA, linewidth=1)) +
  annotate(geom = "text",
           x = -38.05,
           y = -54.0092,
           label = "FWB",
           color = "#fa7876",
           fontface = "bold")  +
  annotate(geom = "text",
           x = -38.05,
           y = -54.011,
           label = "SSB",
           color = "#872ca2",
           fontface = "bold")  #+

# Combine
map <- ggdraw(p_a) + # empty canvas with title to match other plots
  draw_plot(plot.bi.beaches.color, x= 0.07, scale = .8) + # study colonies
  draw_plot(plot.bi.color., 0.07, .48, .5, .5, scale = 1.3) # Bird Iland

#map

#~~~~~~~~~~~~~~~~~#
#  Final plot  ####
#~~~~~~~~~~~~~~~~~#

P_seasonal <- plot_grid(map, p_breeders, p_bm, p_foraging)

P_seasonal
```

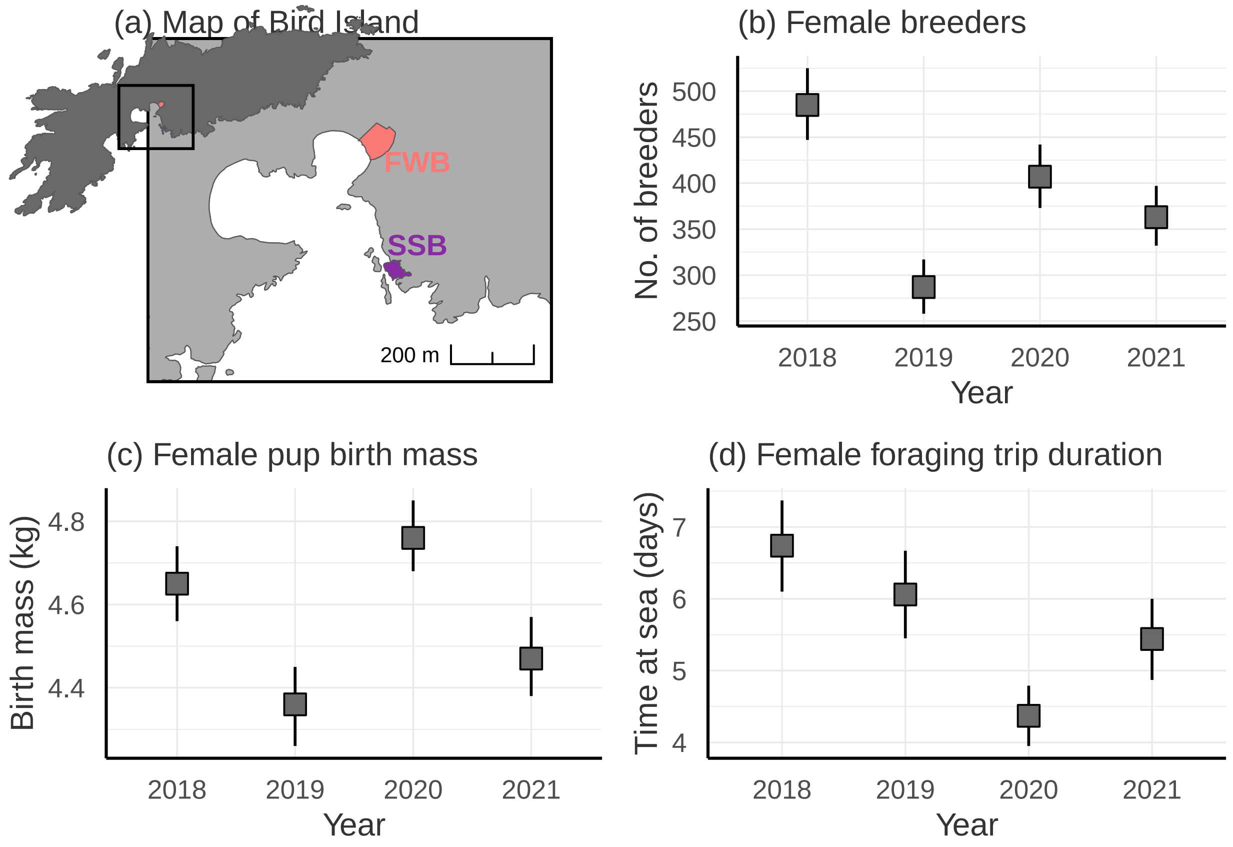

##### Figure 2: forest plots microsatellite models

```
#~~~~~~~~~~~~~~~~~~~~~~~~~~#
#  Make general theme  ####
#~~~~~~~~~~~~~~~~~~~~~~~~~~#

#~~ Use Martin Stoffel's GGplot theme as a base
source(here("Rcode", "anneke_theme.R"))

#~~ Make a list for the theme so it is the same for all figures
gglayer_theme <- list(
  geom_point(shape = 22, size = 3, fill = "black"),
  theme_anneke(),
  theme(axis.line.x = element_line(colour = 'black', linetype='solid'),
        axis.line.y = element_line(colour = 'black', linetype='solid'),
        axis.text.y = element_text(colour = 'black'),
        plot.title = element_text(size = rel(1)))
)

plot_label <- c(
  `(Intercept)` = "Intercept",
  sMLH_msat39_pup = "pup sMLH",
  Pup_SexM = "pup sex [M]",
  Pup_BirthWeight = "pup birth mass",
  sMLH_msat39_mum = "mother sMLH",
```

```r
  Mum_Age = "mother age",
  Age_Tag = "pup age",
  Year2018 = "season [2019]",
  Year2019 = "season [2020]",
  Year2020 = "season [2021]")

#~~~~~~~~~~~~~~~~~~~~~~~~~~~~~~~~~~~~~~~~~~~~~#
#  Apply custom plot function to 3 models  ####
#~~~~~~~~~~~~~~~~~~~~~~~~~~~~~~~~~~~~~~~~~~~~~#

source(here("Rcode", "custom_forest_plot.R"))

#~~ Create labels
lab1 <- paste0("(a) Pup birth mass\nIncl. maternal effects\n n = ", nobs(m1birthmass))
lab2 <- paste0("(b) Pup survival\nIncl. maternal effects\n n = ", nobs(m1survival))
lab3 <- paste0("(c) Pup growth\nIncl. maternal effects\n n = ", nobs(m1growth))

lab4 <- paste0("(d) Pup birth mass\nExcl. maternal effects\n n = ", nobs(m2birthmass))
lab5 <- paste0("(e) Pup survival\nExcl. maternal effects\n n = ", nobs(m2survival))
lab6 <- paste0("(f) Pup growth\nExcl. maternal effects\n n = ", nobs(m2growth))

#~~ Make plots
p.bw <- plot_data_models(m1birthmass, lab1, gglayer_theme)
```

```
## Scale for y is already present.
## Adding another scale for y, which will replace the existing scale.

## Warning: Removed 7 rows containing missing values (`geom_point()`).
```

```r
p.surv <- plot_data_models(m1survival, lab2, gglayer_theme)
```

```
## Scale for y is already present.
## Adding another scale for y, which will replace the existing scale.
## Waiting for profiling to be done...
##
## Waiting for profiling to be done...

## Warning: Removed 8 rows containing missing values (`geom_point()`).
```

```r
p.wg <- plot_data_models(m1growth, lab3, gglayer_theme)
```

```
## Scale for y is already present.
## Adding another scale for y, which will replace the existing scale.

## Warning: Removed 9 rows containing missing values (`geom_point()`).
```

```r
p2.bw <- plot_data_models(m2birthmass, lab4, gglayer_theme)
```

```
## Scale for y is already present.
## Adding another scale for y, which will replace the existing scale.

## Warning: Removed 5 rows containing missing values (`geom_point()`).
```

```r
p2.surv <- plot_data_models(m2survival, lab5, gglayer_theme)
```

```
## Scale for y is already present.
## Adding another scale for y, which will replace the existing scale.
## Waiting for profiling to be done...
##
## Waiting for profiling to be done...

## Warning: Removed 6 rows containing missing values (`geom_point()`).
```

```
p2.wg <- plot_data_models(m2growth, lab6, gglayer_theme)
```

```
## Scale for y is already present.
## Adding another scale for y, which will replace the existing scale.

## Warning: Removed 7 rows containing missing values (`geom_point()`).
```

```
# nb warnings are because I am removing dots and adding squares in the function!

#plot(p.bw)

#~~ Save plots
all_plots <- cowplot::plot_grid(p.bw, p.surv, p.wg,
                                p2.bw, p2.surv, p2.wg,
                                nrow = 2)

all_plots
```

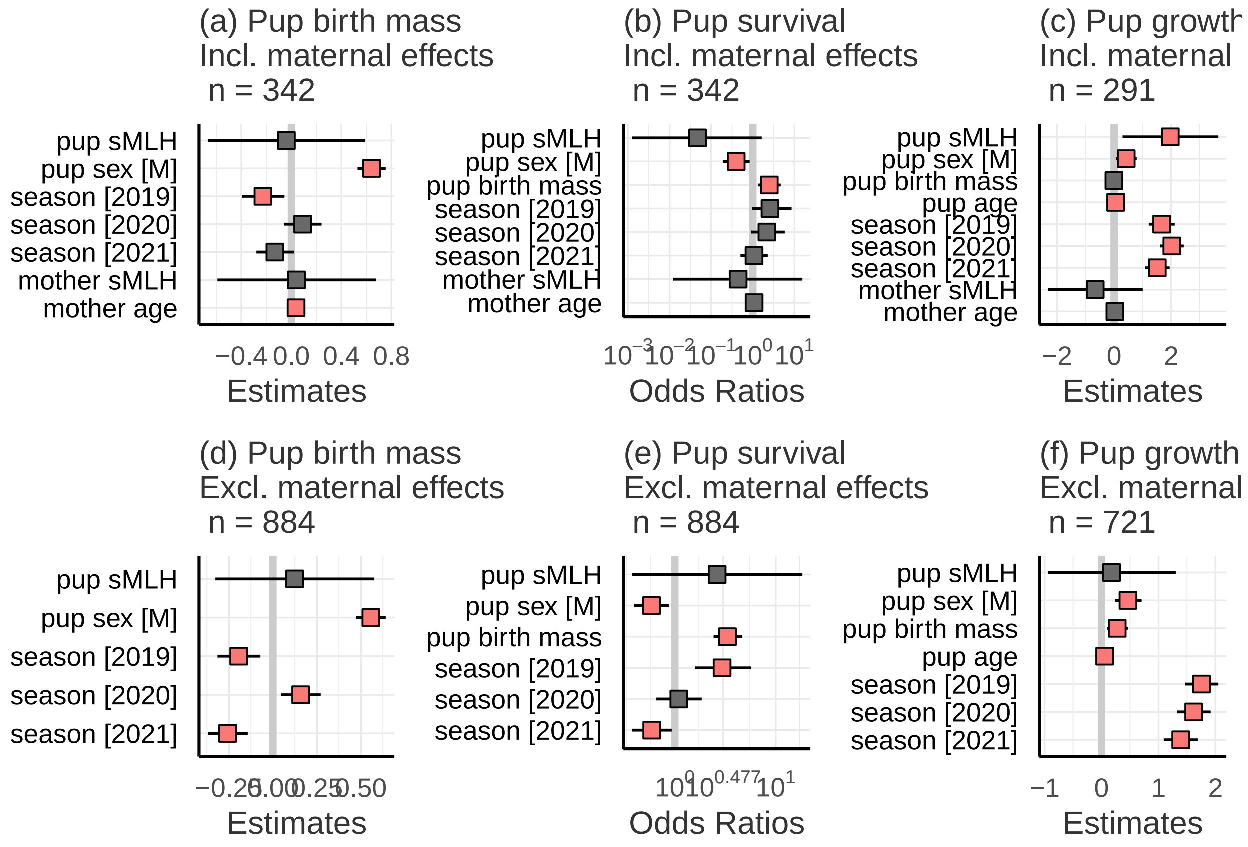

#### Figure 3: forest plots SNP models

```
# Color non sig effects
col1 = "dimgrey"
# Color sig effects
col2 = "#fa7876"

# Use Martin Stoffel's GGplot theme as a base
source("anneke_theme.R")

# Make a list for the theme so it is the same for all figures
gglayer_theme <- list(
  geom_point(shape = 22, size = 3, fill = "black"),
  theme_anneke(),
  theme(axis.line.x = element_line(colour = 'black', linetype='solid'),
        axis.line.y = element_line(colour = 'black', linetype='solid'),
        axis.text.y = element_text(colour = 'black'),
        plot.title = element_text(size = rel(1)))
)

gglayer_theme_alt <- list(
  geom_point(shape = 15, size = 2),
  theme_anneke(),
  theme(axis.line.x = element_line(colour = 'black', linetype='solid'),
        axis.line.y = element_line(colour = 'black', linetype='solid'),
        axis.text.y = element_text(colour = 'black'),
        plot.title = element_text(size = rel(1)))
)

gglayer_theme_alt2 <- list(
  theme_anneke(),
  theme(axis.line.x = element_line(colour = 'black', linetype='solid'),
        axis.line.y = element_line(colour = 'black', linetype='solid'),
        axis.text.y = element_text(colour = 'black'),
        plot.title = element_text(size = rel(1)))
)

plot_label <- c(
  `(Intercept)` = "intercept",
  Last_day = "pup age",
  Age_Days_scale = "age days (scaled)",
  sMLH_SNP.new_scale = "pup sMLH SNP",
  sMLH_SNP.new = "pup sMLH SNP",
  SexM = "pup sex [M]",
  sMLH_SNP.new_mum_scale = "mother sMLH SNP",
  sMLH_SNP.new_mum = "mother sMLH SNP",
  Season1920 = "season [2020]",
  BeachSSB = "colony [SSB]",
  Birth_weight = "pup birth mass",
  froh_mum_scale = expression("mother " ~ italic("F")[ROH] ~ "(scaled)"),
  froh_scale = expression("pup " ~ italic("F")[ROH] ~ "(scaled)"),
  froh = expression("pup " ~ italic("F")[ROH]),
  froh_mum = expression("mother " ~ italic("F")[ROH])) # expression("mother " ~
   ↪  italic("F")[ROH])) #"mother Froh")
```

```r
# Function for custom forest plots
source(here("Rcode", "custom_forest_plot.R"))

lab7 <- paste0("(a) Pup birth mass\nIncl. maternal effects\n n = ", nobs(BW.model.froh))
lab8 <- paste0("(b) Pup survival\nIncl. maternal effects\n n = ",
 ↪  nobs(Survival.model.froh))
lab9 <- paste0("(c) Pup growth\nIncl. maternal effects\n n = ", nobs(Growth_model))

#birth weight
p.BW.froh <- plot_data_models(BW.model.froh, lab7, gglayer_theme)
```

```
## Warning in sjmisc::word_wrap(axis.labels, wrap = wrap.labels): Word wrap is not
## available for expressions.

## Warning in is.na(x): is.na() applied to non-(list or vector) of type
## 'expression'

## Warning: Removed 5 rows containing missing values (`geom_point()`).
```

```r
#survival
p.survival.froh <- plot_data_models(Survival.model.froh, lab8, gglayer_theme)
```

```
## Warning in sjmisc::word_wrap(axis.labels, wrap = wrap.labels): Word wrap is not
## available for expressions.

## Warning in is.na(x): is.na() applied to non-(list or vector) of type
## 'expression'

## Warning: Removed 6 rows containing missing values (`geom_point()`).
```

```r
#growth
p.growth.froh <- plot_data_models(Growth_model, lab9, gglayer_theme)
```

```
## Warning in sjmisc::word_wrap(axis.labels, wrap = wrap.labels): Word wrap is not
## available for expressions.

## Warning in is.na(x): is.na() applied to non-(list or vector) of type
## 'expression'

## Warning: Removed 7 rows containing missing values (`geom_point()`).
```

Plots for figure

```r
AllCurves <- ggplot(data = SurvivorsRM_Day60, aes(x = Age_Days, y = Weight_kg, group =
 ↪  ID)) +
  geom_smooth(method = "lm", se = F, colour = "grey", linewidth = 0.8, alpha = 0.5) +
  geom_point(shape = 15, fill = "black", size = 1.5) +
  geom_abline(slope = 0.081732, intercept = 5.597941, colour = "black", linewidth = 1.1)
   ↪  + #average
  #geom_abline(slope = 0.080226, intercept = 5.502678, colour = "orange", size = 2) +
  #annotate(geom="text", y = 15.5, x = 15, size = 5, label = "FWB: y = 0.080226*age +
   ↪  5.502678", color = "orange") +
  #geom_abline(slope = 0.080821, intercept = 5.511745, colour = "purple", size = 2) +
  #annotate(geom="text", y = 16, x = 15, size = 5, label = "SSB: y = 0.080821*age +
   ↪  5.511745", color = "purple") +
  theme_bw(base_size = 18) + #removes background color
  theme(panel.border = element_blank()) +  #removes border lines
```

```r
  theme(axis.line = element_line(colour = "black")) + #adds in axis lines
  xlab("Age (days)") + #name of x lab
  ylab("Weight (kg)") + #name of y lab
  ggtitle("(e) Growth curves of all pups") #title of plot

#https://quantdev.ssri.psu.edu/tutorials/growth-modeling-basics

p.allcurves <- AllCurves +
  gglayer_theme_alt2  #+
  #theme(axis.title.y=element_text(angle=0, vjust = 0.5))

#Plot for poster with illustrative examples
PlotIllu <- subset(SurvivorsRM_Day60, ID %in% c('H14', 'H1', 'C12', 'T4', 'H7'))

SubIllu <- ggplot(data = PlotIllu, aes(x = Age_Days, y = Weight_kg, colour = ID)) +
  geom_line(linetype = "dashed", linewidth = 1, alpha = 0.8) +
  geom_smooth(method = "lm", se = F, linewidth = 1.2, alpha = 1) +
  #scale_color_brewer(palette="PuOr") +
  scale_color_carto_d(palette = "ag_Sunset") + #colorblind friendly palette
  theme_bw(base_size = 18) + #removes background color
  theme(panel.border = element_blank()) +  #removes border lines
  theme(axis.line = element_line(colour = "black")) + #adds in axis lines
  #theme(legend.position = c(0.15, 0.92)) +
  theme(legend.background = element_rect(fill="NA")) +
  xlab("Age (days)") + #name of x lab
  xlim(0,60) + #x axis limits
  ylab("Weight (kg)") + #name of y lab
  ylim(3.5,16.4) +
  #guides(fill=guide_legend(title="ID")) + #name of legend
  ggtitle("(d) Representative growth curves of 5 pups") #title of plot

p.IDGrowth <- SubIllu + gglayer_theme_alt + theme(legend.position = c(0.12, 0.73)) +
  #theme(legend.background = element_rect(fill="NA")) +
  guides(fill=guide_legend(title="ID"))  #+
  #theme(axis.title.y=element_text(angle=0, vjust = 0.5))

#png(file = here("Growthplot.png"),   # The directory you want to save the file in
   # width = 100, # The width of the plot in inches
   # height = 50)

#plot_grid(SubIllu, AllCurves, labels = "AUTO", label_size = 20)

#dev.off()
```

Final figure

```r
top_row <- cowplot::plot_grid(p.BW.froh, p.survival.froh, p.growth.froh,
                                          ncol = 3, align = 'hv', axis = 'l')

bottom_row <- cowplot::plot_grid(p.IDGrowth, p.allcurves,
                                          nrow = 1 , align = 'hv')
```

```
## `geom_smooth()` using formula = 'y ~ x'
```

```
## `geom_smooth()` using formula = 'y ~ x'
## Warning: Removed 2 rows containing non-finite values (`stat_smooth()`).
## Warning: Removed 2 rows containing missing values (`geom_point()`).
```

```r
Final_figure_froh_h <- cowplot::plot_grid(top_row, bottom_row, nrow = 2)

Final_figure_froh_h
```

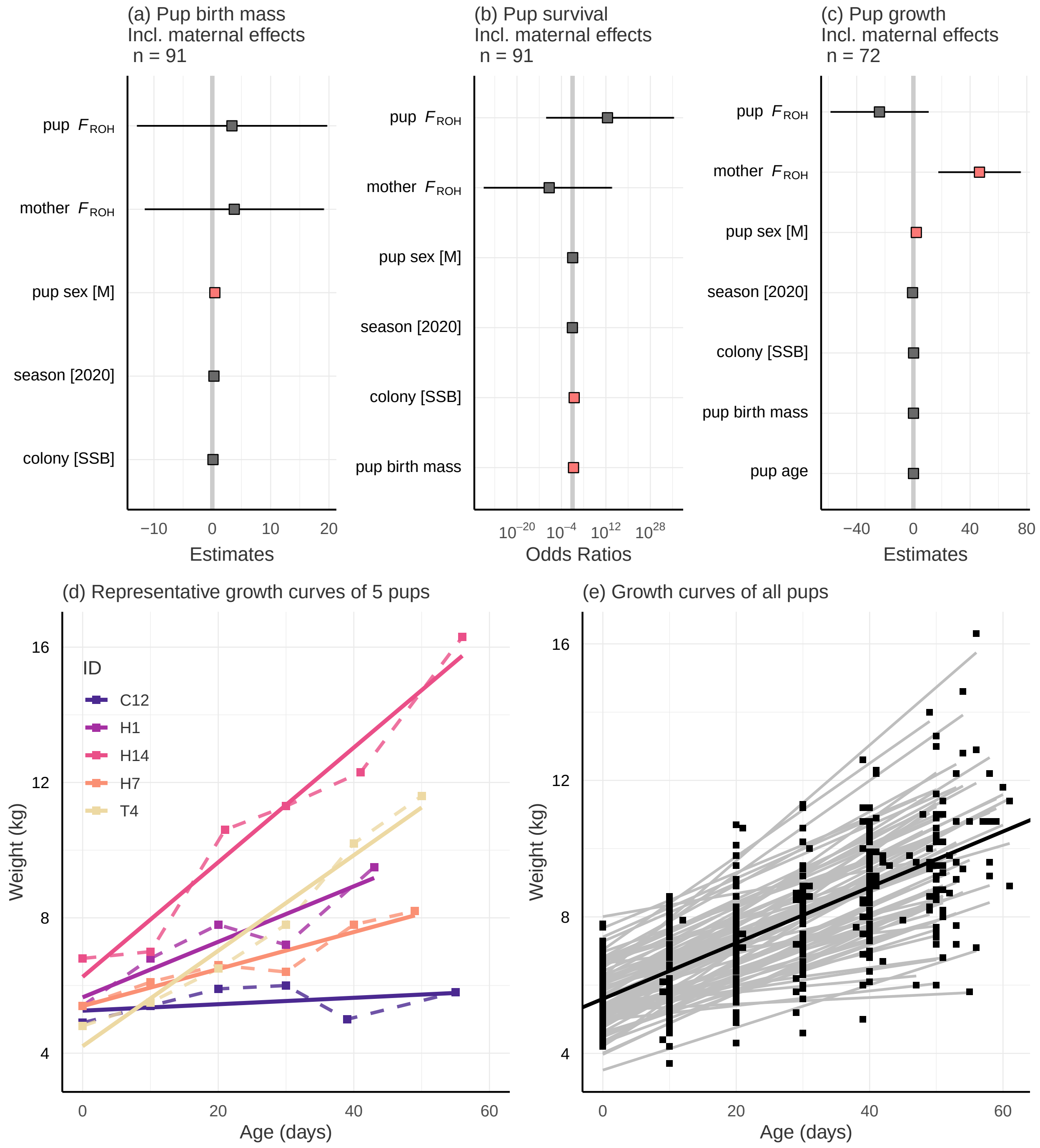

```
#ggsave(here("Figs", "F3_figure_froh_h.jpg"), Final_figure_froh_h, width = 8, height = 8)
```

```
## - Session info ---------------------------------------------------------------
##  setting  value
##  version  R version 4.0.2 (2020-06-22)
##  os       Windows 10 x64
##  system   x86_64, mingw32
##  ui       RTerm
##  language (EN)
##  collate  English_World.1252
##  ctype    English_World.1252
##  tz       Europe/Berlin
##  date     2024-01-12
##
## - Packages -------------------------------------------------------------------
##  ! package      * version    date       lib source
##  P abind          1.4-5      2016-07-21 [?] CRAN (R 4.0.0)
##  P AICcmodavg   * 2.3-2      2023-03-20 [?] CRAN (R 4.0.2)
##  P assertthat     0.2.1      2019-03-21 [?] CRAN (R 4.0.2)
##  P backports      1.2.1      2020-12-09 [?] CRAN (R 4.0.3)
##  P bayestestR     0.13.1     2023-04-07 [?] CRAN (R 4.0.2)
##  P bitops         1.0-7      2021-04-24 [?] CRAN (R 4.0.5)
##  P boot           1.3-25     2020-04-26 [?] CRAN (R 4.0.2)
##  P broom          0.7.6      2021-04-05 [?] CRAN (R 4.0.5)
##  P callr          3.7.0      2021-04-20 [?] CRAN (R 4.0.5)
##  P car          * 3.0-10     2020-09-29 [?] CRAN (R 4.0.5)
##  P carData      * 3.0-4      2020-05-22 [?] CRAN (R 4.0.3)
##  P cellranger     1.1.0      2016-07-27 [?] CRAN (R 4.0.2)
##  P chron        * 2.3-61     2023-05-02 [?] CRAN (R 4.0.2)
##  P class          7.3-17     2020-04-26 [?] CRAN (R 4.0.2)
##  P classInt       0.4-3      2020-04-07 [?] CRAN (R 4.0.2)
##  P cli            3.6.1      2023-03-23 [?] CRAN (R 4.0.2)
##  P codetools      0.2-16     2018-12-24 [?] CRAN (R 4.0.2)
##  P colorspace     2.0-1      2021-05-04 [?] CRAN (R 4.0.5)
##  P CompQuadForm   1.4.3      2017-04-12 [?] CRAN (R 4.0.3)
##  P cowplot      * 1.1.1      2020-12-30 [?] CRAN (R 4.0.5)
##  P crayon         1.4.1      2021-02-08 [?] CRAN (R 4.0.5)
##  P curl           4.3.1      2021-04-30 [?] CRAN (R 4.0.5)
##  P data.table   * 1.14.0     2021-02-21 [?] CRAN (R 4.0.5)
##  P datawizard     0.8.0      2023-06-16 [?] CRAN (R 4.0.2)
##  P DBI            1.1.1      2021-01-15 [?] CRAN (R 4.0.5)
##  P dbplyr         2.1.1      2021-04-06 [?] CRAN (R 4.0.5)
##  P DHARMa       * 0.4.3      2021-07-07 [?] CRAN (R 4.0.5)
##  P digest         0.6.27     2020-10-24 [?] CRAN (R 4.0.3)
##  P doParallel     1.0.16     2020-10-16 [?] CRAN (R 4.0.3)
##  P dotwhisker   * 0.7.4      2021-09-02 [?] CRAN (R 4.0.5)
##  P dplyr        * 1.0.6      2021-05-05 [?] CRAN (R 4.0.2)
##  P e1071          1.7-6      2021-03-18 [?] CRAN (R 4.0.5)
##  P effectsize     0.8.5      2023-08-09 [?] CRAN (R 4.0.2)
##  P ellipsis       0.3.2      2021-04-29 [?] CRAN (R 4.0.5)
##  P emmeans        1.6.3      2021-08-20 [?] CRAN (R 4.0.5)
##  P estimability   1.3        2018-02-11 [?] CRAN (R 4.0.3)
##  P evaluate       0.22       2023-09-29 [?] CRAN (R 4.0.2)
##  P fansi          0.4.2      2021-01-15 [?] CRAN (R 4.0.5)
##  P farver         2.1.0      2021-02-28 [?] CRAN (R 4.0.5)
##  P fastmap        1.1.0      2021-01-25 [?] CRAN (R 4.0.5)
```

```
##  P forcats      * 0.5.1      2021-01-27 [?] CRAN (R 4.0.5)
##  P foreach        1.5.1      2020-10-15 [?] CRAN (R 4.0.3)
##  P foreign        0.8-80     2020-05-24 [?] CRAN (R 4.0.2)
##  P fs             1.5.0      2020-07-31 [?] CRAN (R 4.0.2)
##  P gap            1.2.3-1    2021-04-21 [?] CRAN (R 4.0.5)
##  P generics       0.1.0      2020-10-31 [?] CRAN (R 4.0.3)
##  P ggeffects      1.1.1      2021-07-29 [?] CRAN (R 4.0.5)
##  P ggmap          3.0.2      2023-03-14 [?] CRAN (R 4.0.2)
##  P ggplot2      * 3.4.3      2023-08-14 [?] CRAN (R 4.0.2)
##  P ggsn         * 0.5.0      2019-02-18 [?] CRAN (R 4.0.5)
##  P ggspatial    * 1.1.9      2023-08-17 [?] CRAN (R 4.0.2)
##  P ggstance       0.3.6      2022-11-16 [?] CRAN (R 4.0.2)
##  P ggtext       * 0.1.2      2022-09-16 [?] CRAN (R 4.0.2)
##  P glue           1.4.2      2020-08-27 [?] CRAN (R 4.0.2)
##  P gridtext       0.1.5      2022-09-16 [?] CRAN (R 4.0.2)
##  P gtable         0.3.0      2019-03-25 [?] CRAN (R 4.0.2)
##  P haven          2.4.1      2021-04-23 [?] CRAN (R 4.0.5)
##  P here         * 1.0.1      2020-12-13 [?] CRAN (R 4.0.5)
##  P hms            1.0.0      2021-01-13 [?] CRAN (R 4.0.5)
##  P htmltools      0.5.2      2021-08-25 [?] CRAN (R 4.0.5)
##  P httpuv         1.6.1      2021-05-07 [?] CRAN (R 4.0.5)
##  P httr           1.4.2      2020-07-20 [?] CRAN (R 4.0.2)
##  P inbreedR     * 0.3.2      2016-09-09 [?] CRAN (R 4.0.2)
##  P insight        0.19.3     2023-06-29 [?] CRAN (R 4.0.2)
##  P iterators      1.0.13     2020-10-15 [?] CRAN (R 4.0.3)
##  P jpeg           0.1-10     2022-11-29 [?] CRAN (R 4.0.2)
##  P jsonlite       1.7.2      2020-12-09 [?] CRAN (R 4.0.5)
##  P KernSmooth     2.23-17    2020-04-26 [?] CRAN (R 4.0.2)
##  P knitr          1.44       2023-09-11 [?] CRAN (R 4.0.2)
##  P labeling       0.4.2      2020-10-20 [?] CRAN (R 4.0.3)
##  P later          1.2.0      2021-04-23 [?] CRAN (R 4.0.5)
##  P lattice        0.20-41    2020-04-02 [?] CRAN (R 4.0.2)
##  P lavaan       * 0.6-16     2023-07-19 [?] CRAN (R 4.0.2)
##  P lifecycle      1.0.3      2022-10-07 [?] CRAN (R 4.0.2)
##  P lme4         * 1.1-26     2020-12-01 [?] CRAN (R 4.0.5)
##  P lmerTest     * 3.1-3      2020-10-23 [?] CRAN (R 4.0.5)
##  P lubridate      1.7.10     2021-02-26 [?] CRAN (R 4.0.5)
##  P magrittr       2.0.1      2020-11-17 [?] CRAN (R 4.0.5)
##  P maptools       1.1-1      2021-03-15 [?] CRAN (R 4.0.5)
##  P MASS           7.3-51.6   2020-04-26 [?] CRAN (R 4.0.2)
##  P Matrix       * 1.2-18     2019-11-27 [?] CRAN (R 4.0.2)
##  P merDeriv     * 0.2-4      2022-03-11 [?] CRAN (R 4.0.5)
##  P mgcv           1.8-31     2019-11-09 [?] CRAN (R 4.0.2)
##  P mime           0.10       2021-02-13 [?] CRAN (R 4.0.4)
##  P minqa          1.2.4      2014-10-09 [?] CRAN (R 4.0.5)
##  P mnormt         2.1.1      2022-09-26 [?] CRAN (R 4.0.2)
##  P modelr         0.1.8      2020-05-19 [?] CRAN (R 4.0.2)
##  P munsell        0.5.0      2018-06-12 [?] CRAN (R 4.0.2)
##  P mvtnorm        1.1-1      2020-06-09 [?] CRAN (R 4.0.0)
##  P nlme         * 3.1-148    2020-05-24 [?] CRAN (R 4.0.2)
##  P nloptr         1.2.2.2    2020-07-02 [?] CRAN (R 4.0.5)
##  P nonnest2     * 0.5-6      2023-08-13 [?] CRAN (R 4.0.2)
##  P numDeriv       2016.8-1.1 2019-06-06 [?] CRAN (R 4.0.0)
##  P openxlsx       4.2.3      2020-10-27 [?] CRAN (R 4.0.3)
```

```
##  P parameters     0.21.1     2023-05-26 [?] CRAN (R 4.0.2)
##  P pbapply        1.7-2      2023-06-27 [?] CRAN (R 4.0.2)
##  P pbivnorm       0.6.0      2015-01-23 [?] CRAN (R 4.0.3)
##  P performance    0.10.4     2023-06-02 [?] CRAN (R 4.0.2)
##  P pillar         1.6.0      2021-04-13 [?] CRAN (R 4.0.5)
##  P pkgconfig      2.0.3      2019-09-22 [?] CRAN (R 4.0.2)
##  P plyr           1.8.6      2020-03-03 [?] CRAN (R 4.0.2)
##  P png            0.1-8      2022-11-29 [?] CRAN (R 4.0.2)
##  P processx       3.5.2      2021-04-30 [?] CRAN (R 4.0.5)
##  P promises       1.2.0.1    2021-02-11 [?] CRAN (R 4.0.5)
##  P proxy          0.4-25     2021-03-05 [?] CRAN (R 4.0.5)
##  P ps             1.6.0      2021-02-28 [?] CRAN (R 4.0.5)
##  P purrr        * 0.3.4      2020-04-17 [?] CRAN (R 4.0.2)
##  P qgam           1.3.3      2021-04-27 [?] CRAN (R 4.0.5)
##  P quadprog       1.5-8      2019-11-20 [?] CRAN (R 4.0.0)
##  P R6             2.5.0      2020-10-28 [?] CRAN (R 4.0.3)
##  P rcartocolor  * 2.1.1      2023-05-13 [?] CRAN (R 4.0.2)
##  P Rcpp           1.0.11     2023-07-06 [?] CRAN (R 4.0.2)
##  P readr        * 1.4.0      2020-10-05 [?] CRAN (R 4.0.3)
##  P readxl       * 1.3.1      2019-03-13 [?] CRAN (R 4.0.2)
##  P reprex         2.0.0      2021-04-02 [?] CRAN (R 4.0.5)
##  P RgoogleMaps    1.4.5.3    2020-02-12 [?] CRAN (R 4.0.5)
##  P rio            0.5.26     2021-03-01 [?] CRAN (R 4.0.5)
##  P rlang          1.1.1      2023-04-28 [?] CRAN (R 4.0.2)
##  P rmarkdown      2.25       2023-09-18 [?] CRAN (R 4.0.2)
##  P rprojroot      2.0.2      2020-11-15 [?] CRAN (R 4.0.2)
##  P rstudioapi     0.13       2020-11-12 [?] CRAN (R 4.0.3)
##  P rvest          1.0.0      2021-03-09 [?] CRAN (R 4.0.5)
##  P sandwich     * 3.0-2      2022-06-15 [?] CRAN (R 4.0.2)
##  P scales         1.2.1      2022-08-20 [?] CRAN (R 4.0.2)
##  P sessioninfo    1.1.1      2018-11-05 [?] CRAN (R 4.0.2)
##  P sf           * 1.0-14     2023-07-11 [?] CRAN (R 4.0.2)
##  P shiny          1.6.0      2021-01-25 [?] CRAN (R 4.0.5)
##  P sjlabelled     1.1.8      2021-05-11 [?] CRAN (R 4.0.5)
##  P sjmisc         2.8.7      2021-05-12 [?] CRAN (R 4.0.5)
##  P sjPlot       * 2.8.15     2023-08-17 [?] CRAN (R 4.0.2)
##  P sjstats        0.18.2     2022-11-19 [?] CRAN (R 4.0.2)
##  P sp             1.4-5      2021-01-10 [?] CRAN (R 4.0.5)
##  P statmod        1.4.36     2021-05-10 [?] CRAN (R 4.0.2)
##  P stringi        1.6.1      2021-05-10 [?] CRAN (R 4.0.2)
##  P stringr      * 1.4.0      2019-02-10 [?] CRAN (R 4.0.2)
##  P survival       3.1-12     2020-04-10 [?] CRAN (R 4.0.2)
##  P tibble       * 3.1.1      2021-04-18 [?] CRAN (R 4.0.5)
##  P tidyr        * 1.1.3      2021-03-03 [?] CRAN (R 4.0.5)
##  P tidyselect     1.1.1      2021-04-30 [?] CRAN (R 4.0.5)
##  P tidyverse    * 1.3.1      2021-04-15 [?] CRAN (R 4.0.5)
##  P units          0.7-1      2021-03-16 [?] CRAN (R 4.0.5)
##  P unmarked       1.3.2      2023-07-08 [?] CRAN (R 4.0.2)
##  P utf8           1.2.1      2021-03-12 [?] CRAN (R 4.0.5)
##  P vctrs          0.6.3      2023-06-14 [?] CRAN (R 4.0.2)
##  P VGAM           1.1-9      2023-09-19 [?] CRAN (R 4.0.2)
##  P webshot        0.5.5      2023-06-26 [?] CRAN (R 4.0.2)
##  P withr          2.5.0      2022-03-03 [?] CRAN (R 4.0.5)
##  P xfun           0.40       2023-08-09 [?] CRAN (R 4.0.2)
```

```
##  P xml2           1.3.2      2020-04-23 [?] CRAN (R 4.0.2)
##  P xtable         1.8-4      2019-04-21 [?] CRAN (R 4.0.2)
##  P yaml           2.2.1      2020-02-01 [?] CRAN (R 4.0.2)
##  P zip            2.1.1      2020-08-27 [?] CRAN (R 4.0.2)
##  P zoo            1.8-9      2021-03-09 [?] CRAN (R 4.0.5)
##
## [1] C:/Uni/10_Growth_msats-2017-2020/renv/library/R-4.0/x86_64-w64-mingw32
## [2] C:/Users/localadmin/AppData/Local/Temp/Rtmp2P1F8D/renv-system-library
## [3] C:/Program Files/R/R-4.0.2/library
##
##  P -- Loaded and on-disk path mismatch.
```
